# The m^6^A motif is an anti-3′ splice site

**DOI:** 10.64898/2026.10.16.759739

**Authors:** Kevin R. Roy, Guillaume F. Chanfreau

**Affiliations:** Department of Chemistry and Biochemistry, University of California, Los Angeles, Los Angeles, CA 90095, USA

**Keywords:** m^6^A, METTL3–METTL14, pre-mRNA splicing, 3′ splice site, DRACH motif, GLORI, Nanopore sequencing, AT–AC introns, allele-specific methylation, polypyrimidine tract

## Abstract

N^6^-methyladenosine (m^6^A) is the most abundant internal mRNA modification and can inhibit splicing when present on a 3′ splice site (3′SS). Whether methylation of splicing signals contributes broadly to splicing regulation is unknown. We analyzed in vitro and in vivo m^6^A maps from 31 studies to investigate m^6^A deposition at splicing signals. We show that 3′SS-like substitutions block in vitro methylation by the main m^6^A writer and that intronic m^6^A can be broadly detected in vivo but spares canonical 3′SS adenosines, as opposed to compatible adenosines at other splicing signals, including the 3′SS of AT–AC introns. Human variant alleles compatible with the writer motif are modified at higher rates, and allelic comparisons revealed a preference for A at −4, where most 3′SS carry a pyrimidine. These results show that the specificity of the main m^6^A writer spares the 3′SS from methylation, which reconciles cotranscriptional methylation with efficient splicing.

## Introduction

N^6^-methyladenosine (m^6^A), the most abundant internal modification of mRNA, is written primarily by the METTL3–METTL14 complex and is deposited extensively on chromatin-associated, unspliced pre-mRNA.^1–4^ On this pre-mRNA, the writer can reach the adenosines on which splicing depends, among them the A of the near-invariant AG at every 3′ splice site (3′SS). In *C. elegans*, the METTL16 ortholog METT-10 methylates this adenosine in the *sams-3* and *sams-4* pre-mRNAs and regulates their splicing,^5,6^ and a methylated 3′SS adenosine lowers the binding affinity of U2AF1 for splice sites in vitro.^5,7^ Splicing inhibition by a methylated 3′SS adenosine was also demonstrated in vitro using HeLa nuclear extracts for introns with a weak polypyrimidine tract, and several mouse pre-mRNAs carry the METTL16 motif in a predicted stem-loop at the 3′SS, which raised the possibility that METTL16 regulates splicing more broadly.^5^ A writer with broad access to nascent RNA could therefore impair splicing widely unless the 3′SS adenosine escapes methylation.

Transcriptome-wide maps established the RRACU and DRACH consensus used by METTL3–METTL14 (D = A/G/U, R = A/G, H = A/C/U, Y = C/U; Supplemental Note 1),^8–10^ and splice-site consensus sequences were proposed to rule out direct marking of splice sites,^3^ a reasoning later extended to branch points.^11^ Current models of m^6^A deposition emphasize a positional exclusion zone near exon–exon junctions, attributed to the exon junction complex^12–14^ and to complexes bound before splicing.^4^ None of these studies examined the splice-site adenosines themselves, and METTL16 can methylate outside the METTL3–METTL14 consensus.^5^ Apart from the 3′SS of AT–AC introns, which end in AC, three positions at or near splicing signals can place an adenosine in a DRACH, namely position 4 of the 5′ splice site (5′SS A4) where position 5 is C, the branch point and the first exonic nucleotide (exon +1). If the specificity of the main m^6^A writer governs splice-site methylation, canonical 3′SS adenosines should be spared and compatible adenosines at the other signals methylated. We tested this prediction with a splice-site census of 35 genomes, published in vitro methylation assays on randomized substrates, and single-base m^6^A maps from 31 studies, including realigned libraries using GLORI m^6^A mapping and 16 million HEK293 reads of nanopore direct RNA sequencing (DRS), which reads the modified base itself, allele by allele. Canonical 3′SS adenosines were modified no more often than other YAG adenosines in single-molecule direct RNA reads and stayed below each library’s background in newly synthesized RNA, whereas compatible adenosines at other splicing signals were methylated, showing that writer specificity governs which splicing signals are methylated.

## Results

### Canonical 3′ splice sites cannot match the main m^6^A writer’s consensus

The consensus motifs reported for METTL3-family writers, DRACH in mammals, the stricter RRACU, the yeast RGAC motif and RRACH in plants, all place a C at +1, the position after the methylated A.^8,10,15,16^ The invariant G after the A of a 3′SS YAG therefore excludes the canonical 3′SS adenosine from all of them, whereas the same AG can complete a DRACH motif centered on the next exonic A (Figure 1A). Documented exceptions at +1 U include a writer that methylates GAU in planarians^17^ and a GAU site class in plants for which no writer has been assigned.^18^ Across 35 genomes, only 8.9 % of 3′SS carry a purine before the AG, and about 96 % would need two or more substitutions to form a DRACH (Figure 1B; Table S1). The exceptions are the 3′SS of AT–AC introns, whose terminal C satisfies the +1 requirement (Figure 1B). METTL16 methylates the A of a CAG in U6 snRNA, so the exclusion is specific to the METTL3–METTL14 consensus rather than to this 3′SS-like context (Figure 1C).^5,19^

**Figure 1.**
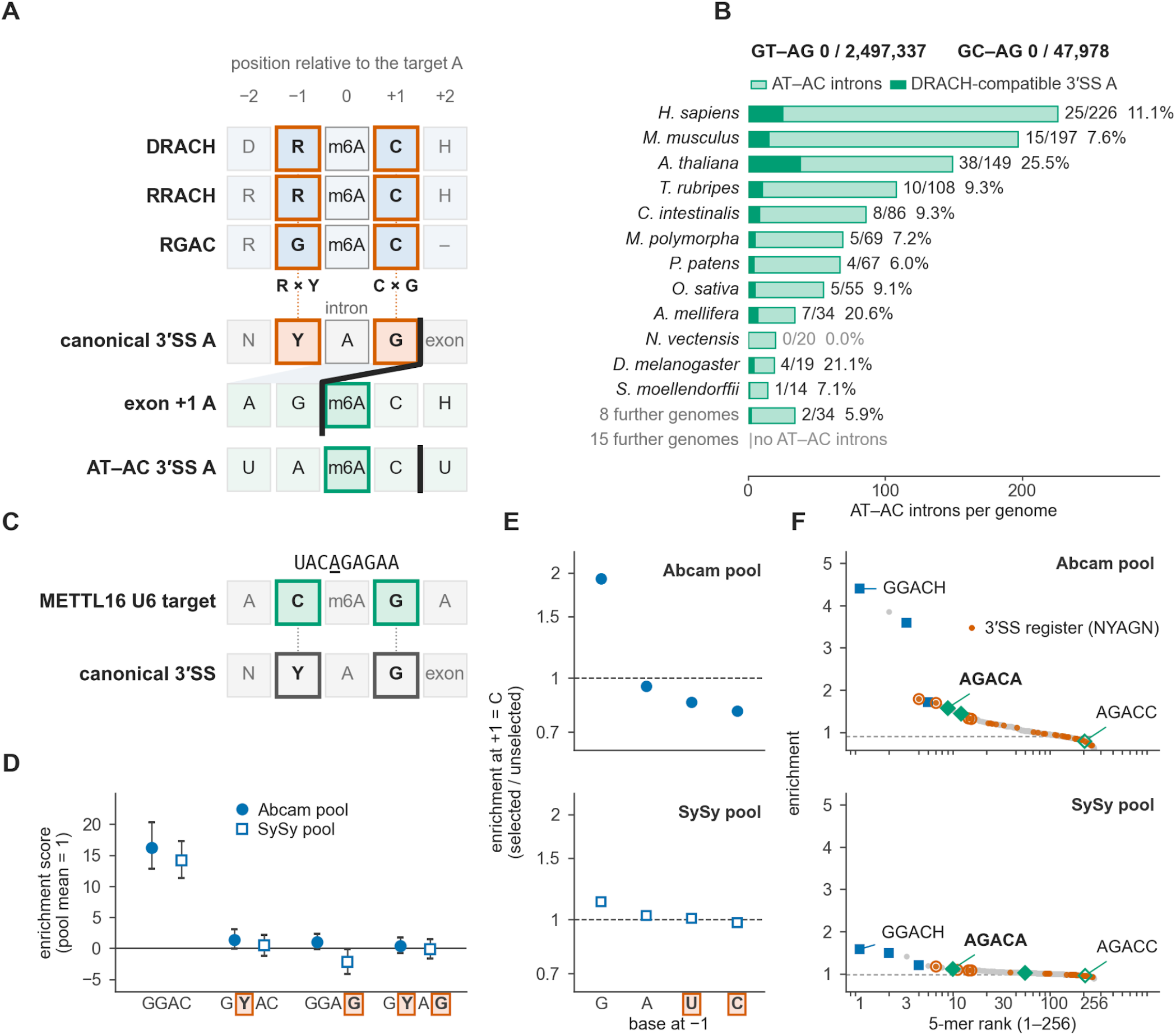
Sequence incompatibility between the METTL3-family m^6^A consensus and canonical 3′ splice sites. **(A)** Methylation and 3′SS motifs aligned on the target A. Rows show three METTL3-family motifs and canonical 3′SS, exon +1 and AT–AC contexts. **(B)** 3′SS A compatibility across 35 eukaryotic genomes (Ensembl protein-coding annotations, which list fewer AT–AC introns than the GENCODE annotation used for the AT–AC analyses). No GT–AG or GC–AG 3′SS A is DRACH-compatible. Bars show AT–AC introns per genome, with compatible sites in the dark segment. **(C)** The METTL16 U6 target UAC<u>A</u>GAGAA carries a YAG sequence around its methylated A, aligned here with the canonical 3′SS. **(D)** In vitro methylation of single-A substrates from randomized libraries selected with two antibodies (Abcam or Synaptic Systems, SySy):^20^ enrichment over the mock library, background-subtracted and scaled to a pool mean of 1, with 95 % intervals. Here and in (E), boxes mark the bases a canonical 3′SS carries, as in (A), and Y stands for C or U. **(E)** Graded preference for the −1 base in the same libraries, on one logarithmic scale: with +1 fixed at C, Abcam enrichment is 1.93 for −1 G and 0.95 for −1 A, neither pyrimidine is enriched over mock, and both pools give the order G > A > U > C. **(F)** Full-pool enrichment ranks of all A-centered 5-mers: blue squares, GGACH; vermilion points, the 3′SS context (open, ACAGN); green diamonds, AGACA and AGACU (open, AGACC); dashed line, the pool median. The median of the 32 5-mers in the 3′SS context is close to the pool median (0.98 versus 0.99); a full-pool rank credits every A-centered 5-mer on a selected molecule, so for AGACA, AGACU and AGACC, which carry an A at −2, the rank is not that 5-mer’s own activity (STAR Methods).

### 3′SS-like substitutions block in vitro methylation

We tested whether the writer itself rejects a 3′SS-like context by reanalyzing a published in vitro assay in which a randomized RNA pool was incubated with purified METTL3–METTL14 and sequenced after m^6^A enrichment with two antibodies (Figures S1A and S1B).^20^ On a background-subtracted scale, GGAC was enriched 14-to 16-fold over the pool mean (Figure 1D). Replacing the −1 G with a pyrimidine cut enrichment by 92 to 97 %, and replacing the +1 C with a G left the substrate at the level of the mock library on the raw scale (1.00- and 0.89-fold, against 3.04- and 1.44-fold for GGAC), so YAG sequences, which carry both substitutions, were not enriched (Figure 1D). With +1 fixed as C, enrichment at −1 fell in the order G > A > U > C, with neither pyrimidine enriched over mock (Figure 1E), and the 32 NYAGN 5-mers centered on the 3′SS A sat near the pool median (Figure 1F). Because antibody selection returns whole molecules and cannot identify which adenosine is methylated (STAR Methods), these libraries only set a ceiling on A at −1 and cannot score the first exonic A, which we test in vivo below. Purified METTL3–METTL14 thus largely rejects sites that carry either base separating a canonical 3′ splice site from DRACH.

### Intronic m^6^A spares canonical 3′SS adenosines

To ask whether cells obey the same rule, we used two mapping methods that do not require a sequence motif to call a site. GLORI scores methylation as the fraction of reads in which an adenosine escapes a chemical deamination that m^6^A resists, above each library’s unconversion background,^21,22^ while nanopore DRS reads the modified base itself (Table S2; Figures S2A–S2D; STAR Methods). We first examined chromatin-associated RNA from A549 cells, counting every position inside an annotated intron (STAR Methods). For all intronic sequences, the m^6^A rate per available DRACH A rose toward the 3′SS and remained detectable to within three nucleotides of the intron end (Figure 2A; Figure S4A), as in newly synthesized polyadenylated RNA.^23^ Because part of that intronic signal comes from positions that are exonic in other transcripts (Supplemental Note 2), we also considered positions that are intronic in every annotated isoform. Sites called in both replicates were too sparse there to resolve the last 15 nucleotides, so we examined these positions in direct RNA reads. Among 9,404 intronic DRACH sites called in both replicates, G at −1 was 3.1-fold more frequent per available site than A, which matches the in vitro preference (Figure 1E). In cells, intronic methylation thus follows the writer’s in vitro preferences to within a few nucleotides of the 3′ splice site.

**Figure 2.**
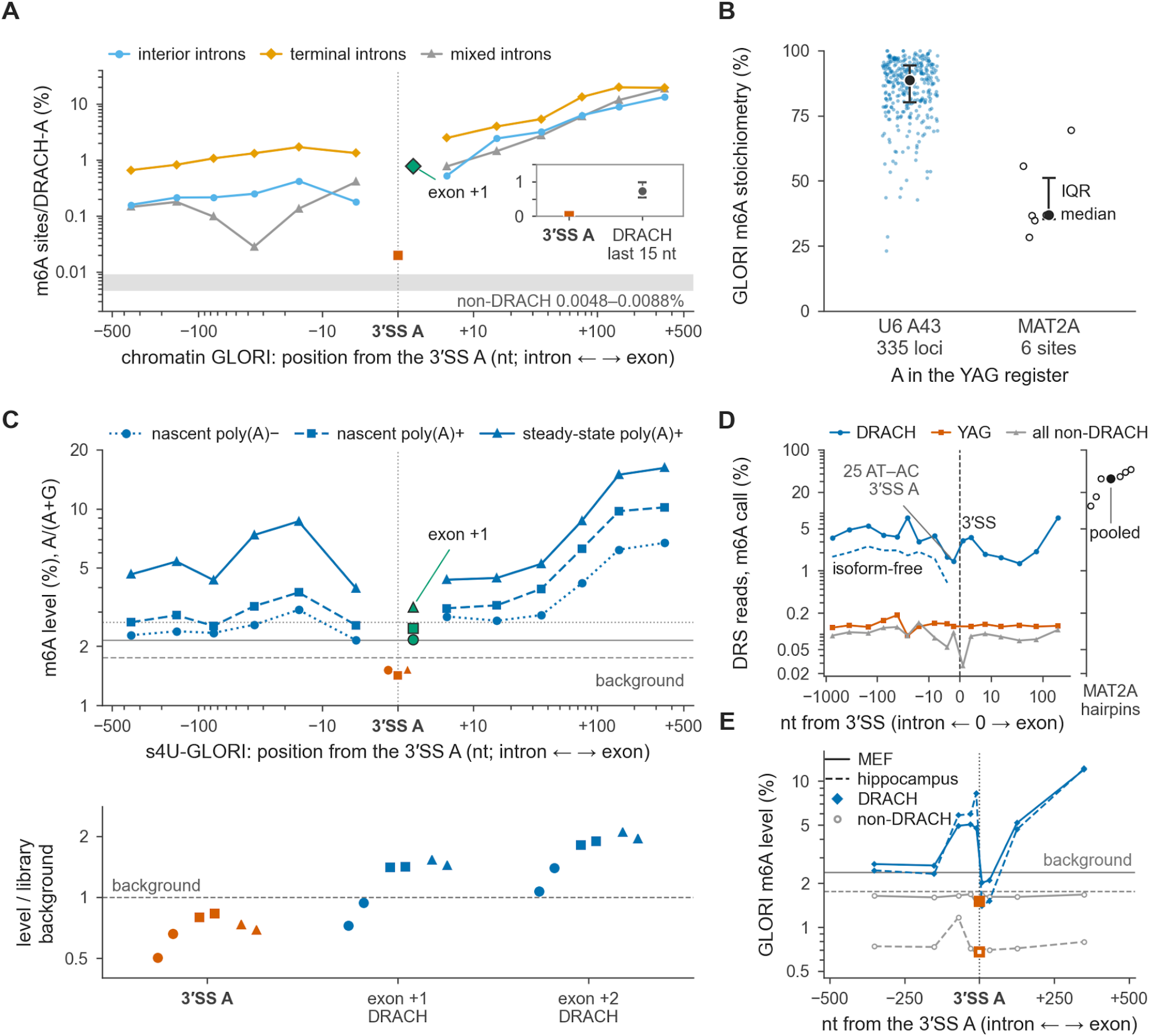
Methylation reaches the intron’s last few nucleotides but rarely the 3′ splice-site A. **(A)** Chromatin GLORI of A549 cells (METTL3-depleted, rescued with wild-type METTL3; GSE272467),^22^ on m^6^A sites called in both replicates, around the 3′SS A of interior introns (n = 39,940), terminal introns (7,819) and mixed introns (6,093), on a symmetric logarithmic axis of distance from the 3′SS A. Curves count detected sites per available DRACH adenosine. Positions inside any annotated intron of a protein-coding gene are counted on the intron side. The inset compares the 3′SS A (11 of 54,819 covered residues) with DRACH adenosines 3 to 15 nt upstream (43 of 5,857; Wilson 95 % interval). The gray band is the intronic non-DRACH rate. The first exonic base reaches 0.78 % (11 of 1,408 boundaries with a spanning DRACH). **(B)** m^6^A stoichiometry in the libraries of (A) at all 335 U6 A43 loci and the 6 *MAT2A* hairpins, with the median and interquartile range: medians 89 % and 37 %. **(C)** The same profile in s^4^U-GLORI of HeLa nascent poly(A)−, nascent poly(A)+ and steady-state poly(A)+ RNA (GSA-human HRA006964; two replicates each),^23^ with the intron classes pooled. Gray lines mark each fraction’s mean background. The 3′SS A lies below every library’s background, and exon +1 rises above it in both poly(A)+ fractions. Below, each library’s m^6^A level at the 3′SS A and at exon +1 and exon +2 DRACH adenosines, relative to that library’s background. **(D)** Direct RNA sequencing (DRS) of HEK293 RNA (ENA PRJEB111137) on a symmetric logarithmic axis centered on the intron–exon boundary. YAG rates stay near 0.12 % everywhere, including at the 3′SS, where the 25 DRACH-compatible AT–AC residues are marked. The six METTL16-methylated *MAT2A* hairpin adenosines are called modified in 33 % of their pooled reads. The dashed curve keeps positions intronic in every isoform. Rates in (C–E) are fractions of reads, not calibrated stoichiometries. **(E)** The same profile in 12 realigned mouse GLORI libraries, pooled by tissue: four mouse embryonic fibroblast (MEF) libraries (GSE210563) and eight dorsal hippocampus libraries (GLORI 3.0; GSE270643). The 3′SS A sits below each tissue’s background, at 1.51 % in MEF and 0.68 % in hippocampus.

The 3′SS adenosine itself followed the same rule. Among GLORI sites called in both replicates, 11 of 54,819 covered 3′SS A residues carried a signal, 37 times less often than DRACH adenosines 3 to 15 nucleotides upstream in the same introns (Figure 2A), and the 3′SS A was called 38 to 57 times less often than those adenosines in polyadenylated RNA from HeLa and HEK293T cells (STAR Methods).^21^ Ten of the 11 chromatin sites lie inside an exon of another transcript, eight of them where another transcript splices to a 3′SS upstream, and only one lies at a position that is intronic in every isoform (Figure S4B). By contrast, GLORI can detect m^6^A in the same YAG context at the METTL16-dependent targets U6 A43 and the *MAT2A* hairpins (Figure 2B; Figure S4C).^19,24,25^ Across 25 further motif-unrestricted GLORI libraries, the canonical 3′SS A was called about as often as the adenosines at positions 3 and 4 of the 5′SS, which the motif also excludes.^26–29^ In primary human bladder tumor and adjacent tissue, the 3′SS A stayed at the background of its YAG context in 17 of 18 GLORI libraries and 0.02 percentage points above it in the remaining tumor library, and in all 18 on reads that run into the intron.^26^ In realigned GLORI libraries of HeLa RNA labeled by a 5-minute pulse of 4-thiouridine (s^4^U), which include nascent RNA,^23^ the canonical 3′SS A was unconverted in 1.40 to 1.54 % of reads, below each library’s background of 1.7 to 3.1 %, whereas DRACH adenosines at exon +2 rose above it in every library (3.1 to 4.4 %), as did those at exon +1 in the four poly(A)+ libraries (2.4 to 3.2 %; Figure 2C). The below-background signal at the 3′SS adenosine therefore reflects little or no methylation rather than a failure to detect it.

In DRS of polyadenylated HEK293 RNA (STAR Methods),^30^ across 43,533 YAG 3′SS covered by unspliced reads, the 3′SS A was modified in 0.120 % of reads, no higher than YAG adenosines elsewhere in the same introns (0.127 %) and far below interior intronic DRACH adenosines (4.37 %; Figure 2D). Intronic DRACH positions that no other transcript’s exon covers are called modified in 2.0 % of reads, less than the pooled rate but far above YAG residues, and they stay methylated up to the intron end (dashed line in Figure 2D). By contrast, the same libraries detected the six *MAT2A* m^6^A sites in the same CAG context (Figure 2D). Read-level m^6^A-SAC-seq, another single-base method, also found no excess at these adenosines over the same 5-mers elsewhere (1.06 % against 0.97 %; odds ratio 1.14, 95 % CI 0.49 to 2.69; STAR Methods).^31^ Nor did adenosine-to-inosine editing account for the low signal, because removing every 3′SS A edited in at least 5 % of reads left the rate at 0.120 % (Figure S4D; STAR Methods).

In 12 realigned mouse GLORI libraries, m^6^A at canonical 3′SS was below background in every library and intron class (1.11 % pooled, and 0.43 to 0.98 % on intron-containing reads in three libraries tested; Figure 2E; Figures S5A–S5D). Of the 12 mouse 3′SS proposed as METTL16 targets,^5^ only *Sorbs1*, whose use rose in *Mettl16*-null embryos, had enough intron-containing reads to test, and none of its 31 such reads was methylated, whereas the METTL16 sites of the *Mat2a* hairpins in the same libraries read 44 to 73 % methylated (Table S3A). Because DRACH adenosines remain methylated to within three nucleotides of the intron end, sequence rather than position protects the 3′SS adenosine from the writer.

### Some DRACH-compatible AT–AC 3′ splice sites are methylated

AT–AC introns offer a natural test of whether a 3′SS adenosine is methylated when its sequence allows. In GENCODE, 39 of 271 human and 18 of 209 mouse AT–AC introns have a DRACH-compatible 3′SS adenosine (STAR Methods). m^6^A was detected at several AT–AC 3′SS in GLORI of HEK293T and HeLa cells and by m^6^A-SAC-seq of HeLa cells (Figures 3A and 3B),^21,32^ including the minor-intron genes *TMEM231*, *PIGB* and *STX10* and the terminal introns of *TSHZ3* and *DSN1*. Most of these 3′SS adenosines also lie inside an exon of another transcript, but the *DSN1* mark already reaches 24 % in nascent poly(A)− RNA (Figure 3F). The DRACH-compatible human *TMEM231* 3′SS was methylated at 41 to 52 % in HEK293T cells, whereas m^6^A was not called at the incompatible mouse ortholog in any of 11 realigned libraries (upper bound 18 %; one-sided exact binomial test against the human level, P = 2 × 10⁻⁷; Figure 3C). *DSN1* 3′SS methylation fell about twofold under METTL3 depletion, remained after METTL14 depletion, and fell in two bladder cancer cell lines depleted of VIRMA (Figure 3D; Tables S3B, S3C and S3E).^21,26^ DRACH-compatible AT–AC 3′SS adenosines were methylated more often than coverage-matched incompatible ones in HEK293T GLORI (3 of 28 against 0 of 95 per replicate; STAR Methods), although only 1 of the 10 compatible sites that no other transcript’s exon covers was methylated, and their methylation was not established in mouse (Figure S5C). In human cells, a 3′SS adenosine whose sequence fits the motif can therefore be methylated.

**Figure 3.**
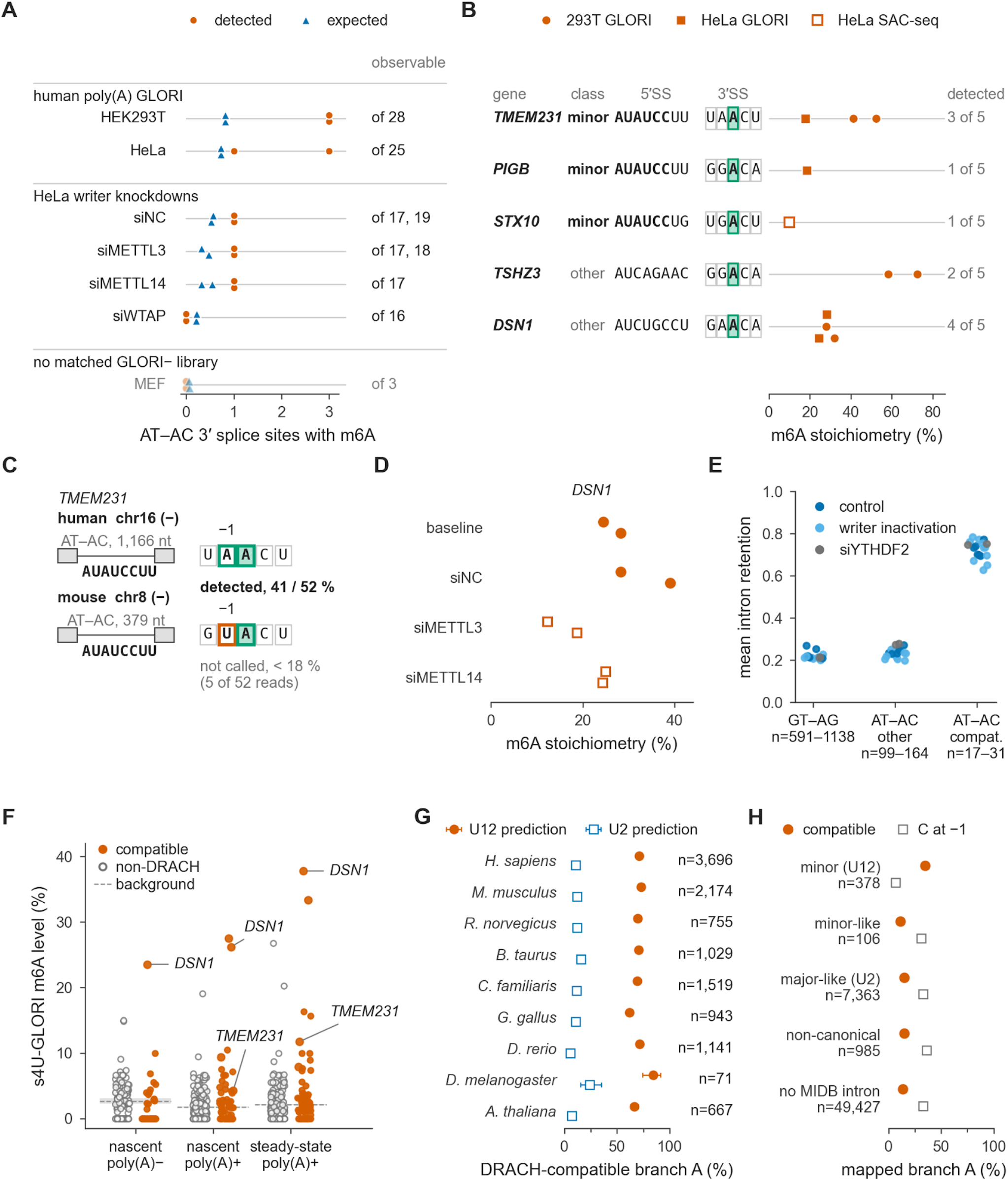
m^6^A at DRACH-compatible AT–AC 3′ splice sites and the DRACH compatibility of minor-intron branch points. **(A)** AT–AC 3′SS m^6^A signals detected by GLORI,^21^ one point per library: vermilion, detected 3′SS A residues; blue, the coverage-matched interior-DRACH expectation. Numbers at right count the compatible sites with at least 10 reads in the library’s matched unconverted (GLORI−) library. MEF libraries have no matched GLORI− library, so their counts are estimated, and they are grouped last. **(B)** m^6^A signals at five genes, one point per library with a detected site; symbols indicate cell line and assay (GLORI or m^6^A-SAC-seq),^32^ the class column marks minor-intron genes^33^ and the right column counts libraries with a detected site. **(C)** Human and mouse *TMEM231* share a minor-class U12 intron, but only the human UAACU 3′SS is DRACH-compatible and methylated. m^6^A at the mouse site is not detected in 11 realigned libraries (5 of 52 reads unconverted, 4 of them in one 7-read library). **(D)** *DSN1* 3′SS methylation in HeLa GLORI, untreated (baseline) and after control siRNA or siRNA against writer subunits (two replicates each; filled, controls; open, knockdowns). **(E)** Intron retention by 3′SS class in 18 unconverted (GLORI−) HEK293T and HeLa libraries (GSE210563) in control conditions (untreated or control siRNA), or after writer inactivation (by siRNA or the METTL3 inhibitor STM2457) or reader depletion (siYTHDF2). **(F)** The same sites in the six s^4^U-GLORI HeLa libraries.^23^ The two classes are level in nascent poly(A)− RNA, and compatible sites are higher in both polyadenylated fractions; the gray band spans each fraction’s two replicate backgrounds. Rates are fractions of unconverted reads, not calibrated stoichiometries. **(G)** DRACH compatibility of the branch A predicted in minor introns of the Minor Intron Database across nine genomes, with Wilson 95 % intervals; n counts introns. It is 62–85 % at U12-predicted and 5.8–24 % at U2-predicted positions. **(H)** Mapped human branch points by intron class; n counts branch A. The branch A is DRACH-compatible at 34.7 % of minor and 15.0 % of major-like branch points, and C, which DRACH excludes at −1, precedes 6.6 % and 33 % of them.

Introns with a DRACH-compatible AT–AC 3′SS were threefold more retained than incompatible ones in every condition (Figure 3E; STAR Methods). Read-level m^6^A-SAC-seq added four more AT–AC genes (Figures S6A–S6D; Tables S3D and S3E). The methylated *DSN1* intron was almost entirely unspliced, and reducing its methylation by writer depletion did not improve its splicing. Therefore, methylation more likely reflects the longer nuclear residence of a slowly spliced intron than direct inhibition of splicing by the mark, as at *STX10* (Supplemental Note 3).

### DRACH-compatible adenosines at other splicing signals are methylated

Exons that begin with AC frequently place their first adenosine in a DRACH context, because of the preceding intronic AG (Figure 1A). In DRS reads of unspliced molecules, exon +1 adenosines in a boundary-spanning DRACH context were modified two orders of magnitude more frequently than non-DRACH exon +1 adenosines (3.12 % against 0.027 %; Figure 4A). While the signal was stronger at 3′ splice sites inside exons of other annotated isoforms, the excess did not come from the mRNAs of those isoforms, because precursor molecules at these sites were marked at least as often (Supplemental Note 4; Figures S7A–S7D). In chromatin GLORI, 11 of 1,408 covered exon +1 adenosines in a spanning DRACH (0.78 %) carried a signal called in both replicates, close to the 0.73 % of exon positions +2 to +15 pooled over all intron classes (71 of 9,678; Figures S4A and S4E). In 16 GLORI libraries of human CD8 T cells,^34^ 22 of 4,862 covered exon +1 adenosines reached 50 % m^6^A stoichiometry, and the exon +1 signal fell 1.80-fold under METTL3 inhibition, whereas the 3′SS A signal did not change (1.03-fold). Methylated exon +1 sites had weaker polypyrimidine tracts, with about one fewer uridine than sites matched for retention and depth (137 sites; logistic regression adjusted for retention, depth and intron length, Benjamini–Hochberg adjusted P = 0.008; Figures S8A and S8B). From unspliced to spliced reads, the exon +1 signal fell 7.7-fold more where splicing destroys the motif than where the upstream exon recreates it, suggesting that some exon +1 methylation is added after splicing (Figures S8C and S8D; Figure S9A). Because the caller’s recall depends on the surrounding sequence, part of this difference could reflect detection rather than methylation. Introns with a methylated exon +1 were more retained only where another transcript’s exon covers the 3′SS, so the mark gives no sign of impairing splicing (Figures S7D, S8E, and S8F).

**Figure 4.**
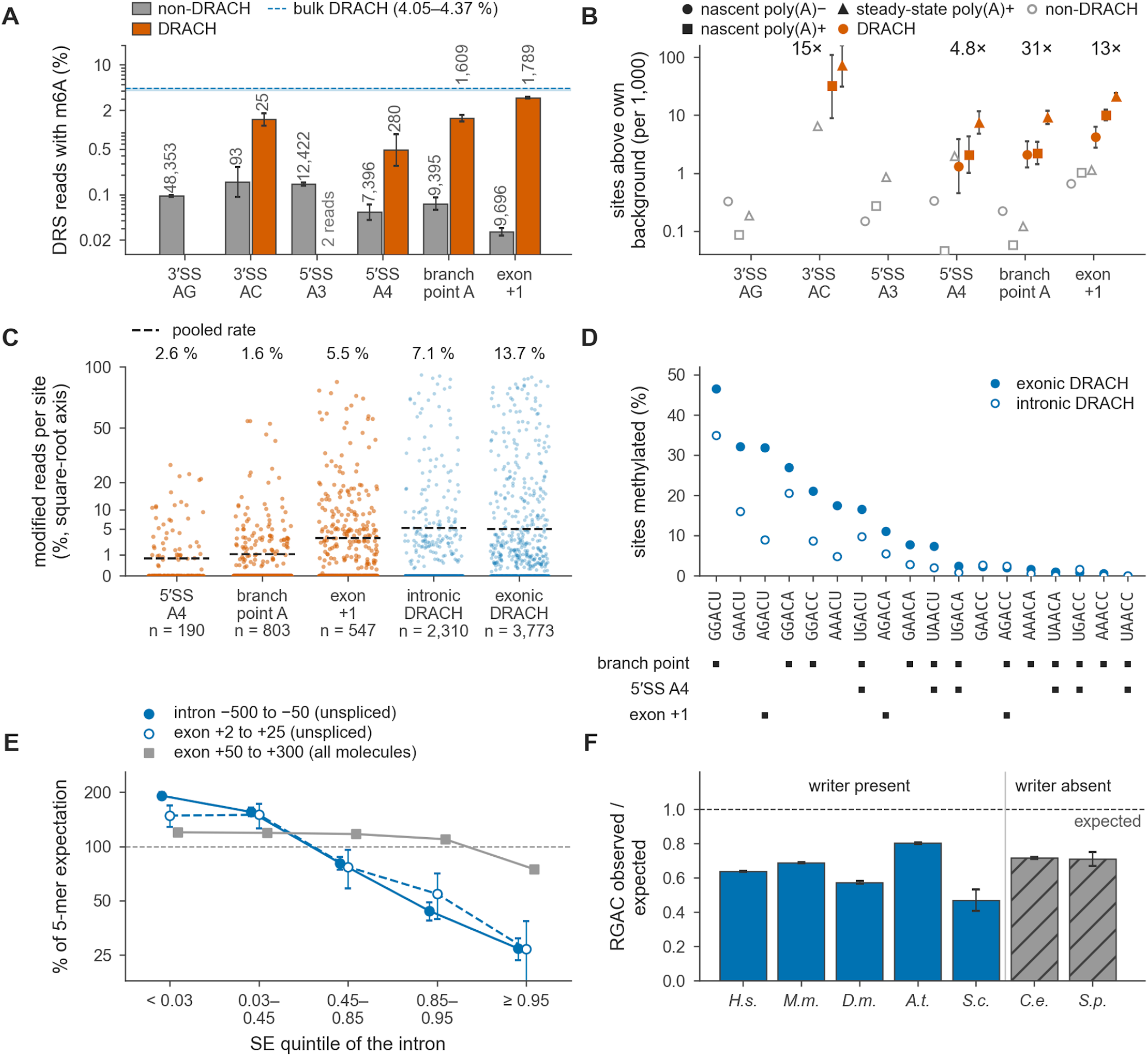
Compatible splicing signals are methylated at levels that depend on their 5-mer and on splicing efficiency. **(A)** m^6^A at five splicing-signal positions in HEK293 direct RNA sequencing, by DRACH context. Bars are modified-read fractions with Wilson 95 % intervals and site counts, and the 5′SS A4 and branch-point columns use positions intronic in every isoform (all positions, 0.71 % and 1.17 %). An AG 3′SS A is never DRACH. **(B)** The same positions site by site in six s^4^U-GLORI HeLa libraries.^23^ Points are sites above their library’s background per 1,000 covered observations, with Wilson 95 % intervals, and numbers are DRACH to non-DRACH ratios on intron-only positions (31- and 4.8-fold at branch points and 5′SS A4, 41- and 13-fold over all their positions; Fisher’s exact test, P < 10⁻¹⁰ except at AT–AC 3′SS A). **(C)** Direct RNA reads site by site for every DRACH-context 5′SS A4, branch-point and exon +1 adenosine with ≥ 20 reads, beside intronic and exonic DRACH references 50–500 nt from a 3′SS. The number above each strip is the percentage of sites methylated in 20 % or more of reads; most carry none (66 % of exon +1 sites). **(D)** The 18 DRACH 5-mers ranked by methylated share in the references of (C), with the 5-mers each signal can reach beneath. The exonic–intronic offset reflects read depth, not biology. **(E)** Methylated share of DRACH sites over the 5-mer expectation by intron splicing efficiency (SE), falling several-fold from the most retained to the best spliced introns on unspliced molecules and 1.6-fold on all molecules, with binomial bootstrap 95 % intervals. **(F)** RGAC motifs −55 to −4 nt from the 3′SS (YAG excluded), counted against a dinucleotide-preserving shuffle, are depleted as much without (gray) as with (blue) a METTL3-family writer. Whiskers show one SD of the null. *H.s.*, *Homo sapiens*; *M.m.*, *Mus musculus*; *D.m.*, *Drosophila melanogaster*; *A.t.*, *Arabidopsis thaliana*; *S.c.*, *Saccharomyces cerevisiae*; *C.e.*, *Caenorhabditis elegans*; *S.p.*, *Schizosaccharomyces pombe*.

We then asked whether adenosines at the 5′SS A4 and the branch point were also methylated when they fit the DRACH motif. For A4 positions that are intronic in every annotated isoform, the DRACH-compatible subset was modified in 0.49 % of DRS reads (12 of 2,471), against 0.05 % for non-DRACH A4 (Figure 4A). Among 86,615 mapped human branch points,^35,36^ 13.9 % of those at an adenosine were DRACH-compatible, twice the intronic-A background and more often in minor than in major introns (Figures 3G and 3H). At positions that are intronic in every isoform, compatible branch adenosines were modified in 1.52 % of DRS reads, against 0.07 % for non-DRACH ones (Figure 4A). We also tested every covered site in newly synthesized RNAs against its own library’s background (STAR Methods). Individually methylated sites were 31-fold more frequent at DRACH-compatible branch points than at incompatible ones, 4.8-fold at 5′SS A4 and 13-fold at exon +1 (Figure 4B), whereas the canonical 3′SS A gave 0.18 elevated observations per 1,000, within the range of incompatible adenosines at the other positions. Mouse branch points behaved similarly (12.8-fold across 12 GLORI libraries), whereas mouse 5′SS A4 gave no signal once positions exonic in another transcript were excluded (Figure S5E). No *S. cerevisiae* branch point or 3′SS is compatible with the stricter yeast RGAC requirement (STAR Methods). Across splicing signals, therefore, sequence rather than the type of signal decides which adenosines are methylated.

### Compatible splicing signals are methylated at or below the level their sequence predicts

We next asked what sets methylation levels at compatible sites. The modified fraction at most compatible sites was modest, as DRS detected m^6^A on at least 20 % of reads for only 2.6 % of 5′SS A4, 1.6 % of branch-point and 5.5 % of exon +1 sites, against 7.1 % of all intronic and 13.7 % of exonic DRACH adenosines (Figure 4C; Figure S10A). DRACH 5-mers differ widely in methylated share (0 to 46.5 %; Figure 4D), and intronic methylation rises several-fold with retention (Figure 4E; Supplemental Note 2), so we compared each class with the expectation from its own 5-mers in the intronic reference at fixed splicing efficiency. At matched sequence and splicing efficiency, branch points reached 67 % of the expected level (95 % CI 32 to 111 %) and exon +1 fell below it (60 %), although not significantly once reads that could be another transcript’s mRNA were left out (STAR Methods; 79 %, 95 % CI 52 to 113 %), and 5′SS A4 was too imprecise to resolve (133 %; Figures S10B and S10C). In HeLa s^4^U-GLORI, exon +1 on intron-retaining molecules reached 52 % of the level at neighboring DRACH adenosines (95 % CI 21 to 100 %; STAR Methods). RGAC motifs are depleted upstream of 3′SS as much in species without a METTL3-family writer as in species with one, so this depletion is not a safeguard against methylation (Figure 4F). Compatible branch points and first exonic adenosines are methylated at or below the level their sequence predicts, whereas 5′SS A4 remains unresolved, so a compatible sequence permits methylation without fixing its level.

### Human genetic variants change methylation according to their match to DRACH

If sequence dictates methylation, sequence variation within or near DRACH sequences should change methylation accordingly. Long-read and single-base sequencing can resolve allele-specific m^6^A, including variants inside the motif, as shown in F1 hybrid mouse cells, a human lymphoblastoid cell line and F1 hybrid mouse brain.^37,38^ We therefore tested this at positions that complete the motif (Figure 5A). DRS reads of the DRACH allele were called modified 9.1 times more often than reads of the non-DRACH allele in HEK293 cells (25 sites, 95 % CI 4.5 to 77) and 15.8 times more often in HG002 cells (143 sites, 95 % CI 7.9 to 63; Figure S11A). In GM12878 cells sequenced on a single flow cell, the DRACH allele was called modified 38.5 times more often (43 sites, 95 % CI 14 to 219; Figures S11A and S11B). GLORI provided similar results in HEK293T cells (11.1-fold, 54 sites, 95 % CI 6.2 to 18.2; Figures S11B and S11C) and in HeLa cells, where DRACH alleles carried 5.4 times more methylation than non-DRACH ones (116 sites, 95 % CI 2.3 to 16). In both assays variants lay at −1 and +1, the two positions at which a canonical 3′ splice site departs from DRACH. Variants that kept the motif or lay 3 to 5 nucleotides away did not change m^6^A levels in HEK293 and HG002 cells (DRS ratios 0.91 to 1.01). In GM12878 cells, the ratio was 0.45 at the 28 sites where a variant kept the motif (95 % CI 0.20 to 0.87) and 1.00 at the 159 sites with a variant 3 to 5 nucleotides away. In HeLa cells, inhibiting METTL3 or depleting WTAP lowered methylation of the DRACH alleles from 50 % to 14 and 23 %, whereas depleting the reader YTHDF2 left it at 50 % (Figure S11D).

**Figure 5.**
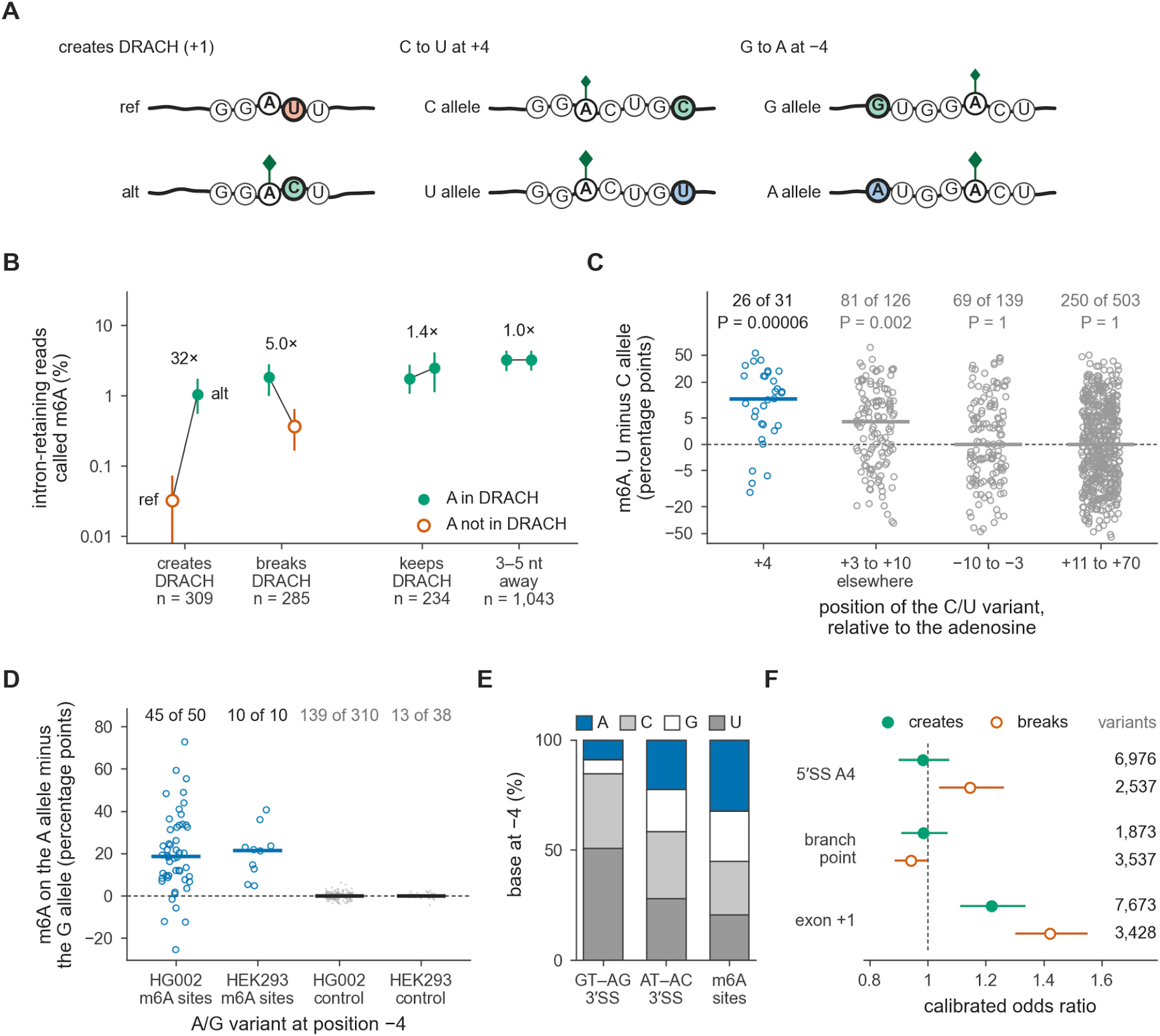
Allelic variants change methylation according to the match to the writer’s motif. **(A)** Heterozygous variants that complete a DRACH motif (left, here at +1), change a C to a U four nucleotides downstream of a DRACH adenosine (middle) or change a G to an A four nucleotides upstream (right). Diamonds mark m^6^A, and the larger diamond the more methylated allele. The varied base is red where it blocks methylation, green where it permits it and blue where it raises it. **(B)** HG002 direct RNA reads called m^6^A on each allele of heterozygous variants within 5 nt of an adenosine, in introns that are exonic only in retained-intron models, counting intron-retaining reads only. Variants at −2 to +2 create or break a DRACH, and controls keep the motif or lie 3 to 5 nt away. Whiskers are 95 % site-bootstrap intervals. **(C)** HEK293T GLORI methylation of the U allele minus the C allele at heterozygous C/U variants near an m^6^A site, on a pseudo-log axis. Points are sites and bars are medians, and counts give the sites with more m^6^A on the U allele, with P from a two-sided sign test. Sites were not selected on the difference. **(D)** HG002 and HEK293 direct RNA reads called m^6^A on the A allele minus the G allele at heterozygous A/G variants four nucleotides upstream of an m^6^A site, beside control adenosines that are unmethylated and form no DRACH on either allele. Points are sites and bars are medians, and counts give the sites with more m^6^A on the A allele (two-sided sign test, P = 4 × 10⁻⁹ in HG002 and P = 0.002 in HEK293). **(E)** The base four nucleotides upstream of the 3′SS A at 250,300 human GT–AG 3′ splice sites (Ensembl) and 271 AT–AC 3′ splice sites (GENCODE v46), and of the methylated adenosine at 213,750 m^6^A sites mapped by GLORI in HEK293T cells. A pyrimidine occupies this position at 85 % of GT–AG and 58 % of AT–AC 3′ splice sites, and an A at 9 %, 23 % and 32 % of the three sets. **(F)** Variants that create DRACH (filled) or break it (open) at 5′SS A4, branch points and exon +1 in 76,215 human genomes (gnomAD v4.1.1). Points are calibrated odds ratios of observed among possible substitutions against motif-neutral ones, with 95 % CIs, and numbers are observed variants. A ratio below 1 would indicate purifying selection.

These results also extended to intronic sites on pre-mRNA. In intron-retained RNAs from HG002 cells, variants that created a DRACH motif raised m^6^A levels 32-fold (309 sites), whereas variants that broke the motif lowered them fivefold (285 sites), and the same controls stayed unchanged (Figure 5B; Figures S11E and S11F). Every site with an allele difference lay outside splicing signals. At the signals themselves, only five variants, all in HG002, were covered deeply enough to test, and they carried almost no m^6^A calls on either allele (4 in 1,278 reads). In human cells, a single substitution at −1 or +1 is enough to change methylation of the adenosine it flanks, consistent with in vitro results (Figures 1D and 1E).

### Allelic variants reveal two writer preferences outside the core motif

At C/U variants four nucleotides downstream of an m^6^A mapped by GLORI in HEK293T cells, the U allele carried more m^6^A at 26 of 31 sites and the C allele at 4 (median difference, 11 percentage points; two-sided sign test, P = 6 × 10⁻⁵), whereas C/U variants 3 to 10 nucleotides upstream showed no bias (69 against 70 sites; Figure 5C; Figure S11G). This preference was confirmed in HeLa cells, where the +4 U alleles carried more m^6^A at 30 of 34 such sites (P = 6 × 10⁻⁶), and it survived the same writer and reader perturbations (1.35 to 1.63 against 1.50 in the controls; Figure S11H), suggesting that the +4 U affects methylation directly rather than by increasing stability of the methylated transcript. Because each comparison is made between two alleles of one site, this result extends to human cells the preference for uridine at +4 found in yeast and mouse cells.^39,40^ Downstream of the motif, a U was also more methylated than an A (28 of 37 sites, P = 0.003), whereas A and C did not differ, so the effect is a preference for uridine rather than an aversion to cytidine. The same studies also inferred a preference for adenosine four nucleotides upstream.^39,40^ At A/G variants at −4, direct RNA reads of the A allele were more methylated at 45 of 50 sites in HG002 cells and at all 10 in HEK293 cells (two-sided sign test, P = 4 × 10⁻⁹ and P = 0.002), while false-positive calls at unmethylated adenosines showed no such bias (Figure 5D). GLORI, which can read A/C and A/U variants, agreed at all 11 such sites in HEK293T and HeLa cells (P = 0.001). The two preferences combine multiplicatively, as expected if they act independently (STAR Methods).

The polypyrimidine tract departs from the writer’s preference for A at −4, so a pyrimidine lies four nucleotides upstream of the 3′SS A in 85 % of 250,300 human GT–AG 3′ splice sites and an A at only 9 %, whereas an A occupies that position at 32 % of the sites methylated in HEK293T cells (Figure 5E).

### Allelic differences in methylation are tolerated

At the 490 exonic variants in HG002 whose alleles differ in methylation by at least 10 percentage points, the more methylated allele was similar in abundance to the other. Across all 11,959 exonic variants, a 50-point difference in methylation changed allelic abundance by 0.99-fold (95 % CI 0.94 to 1.03), and METTL3 inhibition did not shift allelic balance in HEK293T cells. In other studies, the more highly methylated allele was the more abundant one in hybrid mouse embryonic stem cells and at several genes in GM12878 cells,^37^ but the less abundant one in hybrid mouse postnatal brain^38^ and in yeast and human–mouse interspecies hybrids.^40^ Averaged over thousands of human sites, methylation at a single site therefore does not measurably change the transcript abundance of the allele that carries it, although individual genes can depart from this average.

Variants that create or break the DRACH motif at splicing signals are frequent in human populations, and 1.5 % of them reach an allele frequency of at least 1 % (402 of 26,028 variants in 76,215 genomes). No class showed a significant deficit against motif-neutral variants at the same positions, after calibrating on adenosines with no splicing role (Figure 5F). Creating or breaking a DRACH motif at splicing signals is therefore tolerated in human populations, as expected when 83 to 94 % of these sites are methylated in less than 5 % of reads (Figure 4C).

## Discussion

The motif preferred by the main m^6^A writer complex is an anti-3′ splice site. The purine and C that the DRACH motif of METTL3–METTL14 requires sit at the two positions where a canonical 3′SS carries a pyrimidine and a G (Figure 1A). In vitro, either mismatch alone removed most of the enrichment for methylated substrates (Figure 1D). Canonical 3′SS adenosines were modified no more often than other YAG adenosines in single-molecule direct RNA reads, stayed below each library’s background in newly synthesized RNA and, among GLORI sites that both replicates called, were methylated far less often than DRACH adenosines near the end of the same introns (Figure 2). In CD8 T cells, the residual signal did not respond to METTL3 inhibition.^34^ Methylation of a canonical YAG 3′SS adenosine, where it occurs, would therefore point to a writer other than METTL3–METTL14, such as METT-10 at the *sams-3* and *sams-4* 3′SS of *C. elegans*.^5,6^ At the mouse 3′SS proposed as METTL16 substrates, only *Sorbs1* was covered well enough to test, and its intron-containing reads were unmethylated (Table S3A). By contrast, DRACH-compatible adenosines at the first exonic nucleotide and at branch points were methylated, at or below the level their sequence predicts (Figure 4; Figure S10C), and so were some DRACH-compatible AT–AC 3′SS (Figure 3). These patterns fit the proposal that m^6^A is a default mark of immature RNA that promotes nuclear retention and decay.^41,42^ A writer with broad access to nascent RNA can mark intron-containing transcripts throughout their length, AT–AC 3′SS included, whereas the 3′SS adenosine of GT–AG and GC–AG introns, whose methylation can impair U2AF1 binding, escapes methylation because its sequence is incompatible (Figures 6A and 6B).^5^

**Figure 6.**
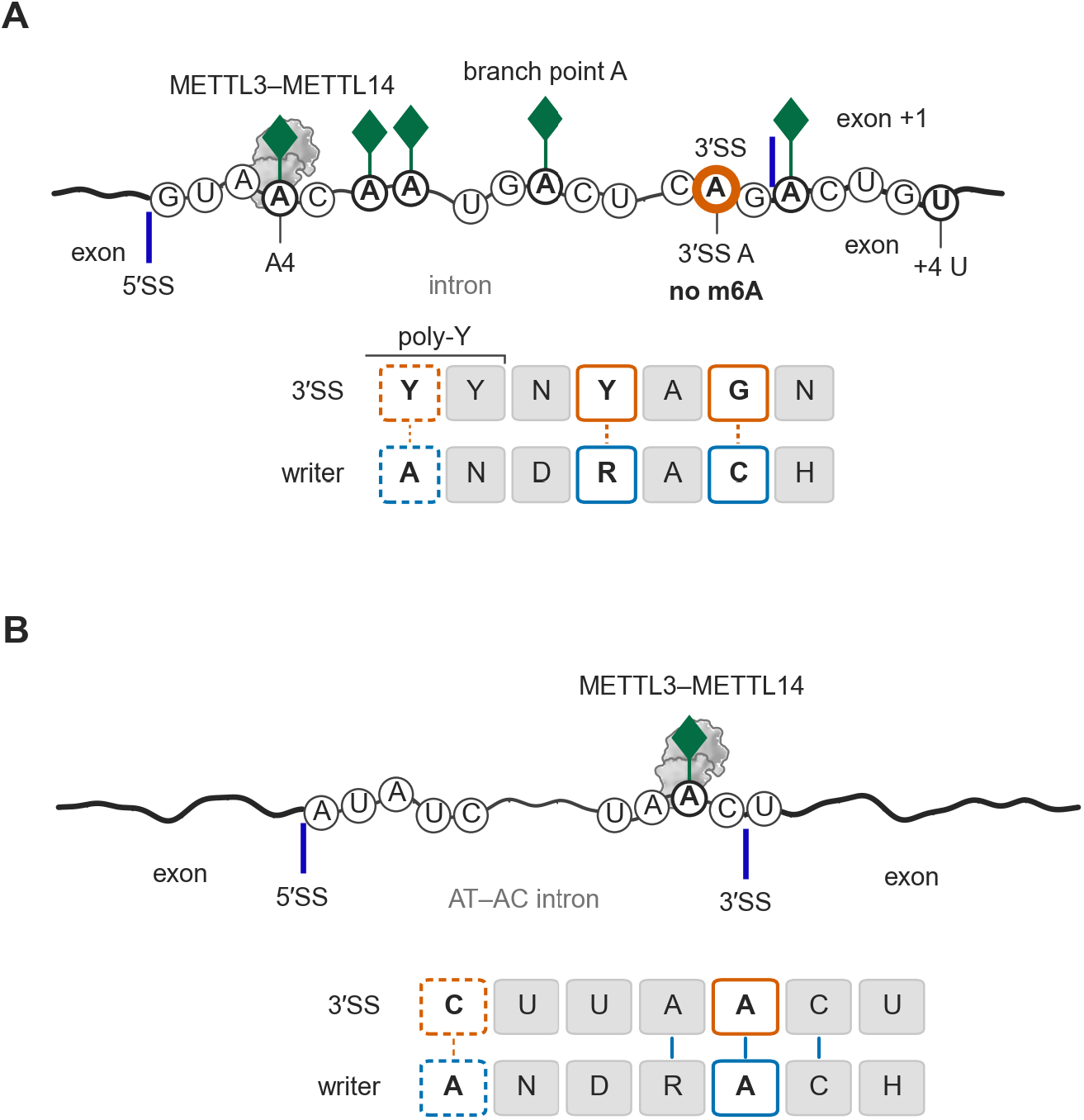
Sequence specificity separates METTL3–METTL14 substrates from canonical 3′ splice sites. **(A)** METTL3–METTL14 can methylate DRACH sites in unspliced pre-mRNA, including the 5′SS A4 of the subset whose position 5 is C rather than the consensus G, intronic A residues, compatible branch points and the first exonic A when the intron’s AG completes the motif, shown with a U at +4, which raises methylation (Figure 5C). The marked positions are examples of compatible sites, not modifications carried together on one molecule. The canonical YAG 3′SS is incompatible: its −1 pyrimidine and +1 G differ from the purine and C required by DRACH, and its A (vermilion ring) carries no m^6^A. The register extends to −4, where the polypyrimidine tract usually places a pyrimidine and the writer prefers an A (Figures 5D and 5E); dashed boxes mark preferences, solid boxes requirements. **(B)** The same writer at an AT–AC 3′ splice site whose own sequence admits the motif. AT–AC introns end in AC rather than AG, so a DRACH can close on the 3′SS A itself, and in *TMEM231* the 3′SS context is UAACU and the A is methylated. *TMEM231* carries a C at −4 and is methylated nonetheless. Compatible AT–AC introns are also spliced less efficiently (Figure 3E), and the cause of that association is unresolved. Discs spell the AT–AC donor A U A U C and the 3′SS U A A C | U. The two panels are schematics of different introns, not a controlled sequence comparison. The protein surface is the deposited structure of METTL3–METTL14 (PDB 5IL2),^43^ oriented by its cofactor-marked catalytic face because that structure lacks substrate RNA. The transcript is drawn as a single strand with heavier lines for exons. Transcript positions are schematic. The schematic was drawn by plotting code written with Claude Code (Anthropic; Claude Opus 5, Opus 5.5, Fable 5 and Fable 5.1).

Methylation near splice sites can promote splicing, as it does for adenoviral late transcripts,^44^ and how methylation of 5′SS A4, branch points and exon +1 affects recognition by the spliceosome remains uncertain. Position 4 of the 5′SS pairs with U1 and then U6 snRNA, and the branch-point adenosine bulges from its helix with U2 snRNA, so methylation at either position could alter recognition by the snRNAs.^45–47^ Structural models place the exon +1 methyl group against the U2AF1 backbone in its preferred orientation and clear of it in the other (Figures S12A–S12D).^7^ Loop 1 of the U5 snRNA, which base-pairs with exon sequences at both splice sites in yeast, could also be sensitive to the mark at the second step.^48^ Intron retention gives no sign that m^6^A at exon +1 affects splicing (Figures S7D, S8E, and S8F). Part of the exon +1 signal may be deposited after splicing, where the upstream exon recreates the motif, although the exon junction complex deposited at that junction may limit it (Figures S8C and S8D; Figure S9A).^12,13^ Whether methylating an AT–AC 3′SS affects U12-type splicing, in which ZRSR2/Urp contacts the 3′SS, is unknown.^49^ These sites nonetheless show that a 3′SS adenosine can be methylated where its sequence permits, as at the minor-intron 3′SS of *TMEM231* and the AT–AC 3′SS of *DSN1* (Figures 3B–3D).

Because the exclusion is written into the YAG sequence itself, it should extend to alternative and cryptic 3′SS, which share that context, so new 3′SS can arise without being methylated. This exclusion comes from the writer’s specificity, unlike the depletion of m^6^A near exon junctions, attributed to the exon junction complex once the junction forms and to complexes bound before splicing (Figure S4A).^4,12–14^ Those studies scored DRACH motifs, which by definition carry a purine at −1 and C at +1, so the canonical 3′SS A was never part of them, and a regional depletion near the junction does not by itself explain why that nucleotide is a poor substrate. A boundary-spanning DRACH at exon +1 is nonetheless methylated inside that region, where methylation levels are lower. The same rule makes the effect of sequence variation on m^6^A predictable. A heterozygous variant that completes the motif next to an adenosine brings methylation to its own allele (Figure 5B; Figure S11). At the splicing signals where the motif can occur, variants that create or break it are frequent and tolerated in human populations (Figure 5F). No single substitution, by contrast, can make a canonical 3′ splice site fit the motif without destroying its splice site identity (Figures 1A and 1B). Allelic comparisons also resolve two preferences beyond DRACH, for U at +4 and A at −4 (Figures 5C and 5D). At most 3′ splice sites the polypyrimidine tract places a pyrimidine at −4, a third departure from the writer’s preferences. Whether the exclusion of YAG from methyltransferase specificity was inherited or narrowed by selection cannot be established. The MT-A70 family also includes bacterial DNA methyltransferases,^50^ and the last eukaryotic common ancestor already had a complex spliceosome.^51^ Whatever its origin, this exclusion reconciles broad cotranscriptional methylation with efficient splicing by sparing the one adenosine whose methylation can impair the removal of AG-dependent introns.

### Limitations of the study

This study reanalyzes public data and adds no new perturbation experiments. The inhibition of human splicing by a 3′SS m^6^A was shown in HeLa nuclear extracts and with a reporter expressed in HeLa cells^5^ and was not tested for endogenous introns here. The randomized-substrate in vitro methylation assays analyzed selected entire RNA molecules with an antibody and cannot identify the methylated adenosine within the randomized sequence. DRS m^6^A rates are fractions of reads called modified by a model rather than calibrated stoichiometries, they bound rather than measure methylation at YAG adenosines, and a motif-agnostic caller can still carry the writer’s 5-mer preference. GLORI background levels differ between libraries, and several public GLORI libraries carry no unique molecular identifiers, so we compared classes within each library or set rather than across sets, and in chromatin-associated A549 RNA we used only sites called in both replicates, which favors highly methylated and deeply covered sites. Among such sites in HEK293T and HeLa cells, canonical 3′SS adenosines were called 38 to 57 times less often than DRACH adenosines near the end of the same introns but about twice as often as other YAG adenosines in those introns, and matching for distance from the intron end left that comparison inconclusive (1.4-to 1.6-fold, with intervals that include both parity and a two-to threefold excess). The conclusion about the 3′SS adenosine therefore rests most directly on direct RNA reads, in which each read is one molecule, and on newly synthesized RNA, where removing duplicate reads did not change it. The mouse, CD8 T-cell and read-level m^6^A-SAC-seq counts were not deduplicated. Finally, the libraries analyzed did not sample the gonads, where METTL16 is most abundant, nor the early embryos in which two of the proposed mouse 3′SS responded to its loss, so METTL16 activity at 3′SS in those tissues remains untested.

## Resource availability

### Lead contact

Requests for further information and resources should be directed to and will be fulfilled by the lead contact, Guillaume F. Chanfreau.

### Materials availability

This study did not generate new unique reagents.

### Data and code availability

● This paper analyzes existing, publicly available data, whose accession numbers are listed in the key resources table and Table S2; the HG002 direct RNA data are the Oxford Nanopore open data release UHRR_HG002_2026.06 (s3://ont-open-data/UHRR_HG002_2026.06/). No new sequencing data were generated for this study. Processed per-site tables derived from these data, including the pooled table of three flow cells that underlies the direct RNA single-molecule analyses, have been deposited at Dryad and are publicly available as of the date of publication at https://doi.org/10.5061/dryad.xpnvx0kz4, and Tables S1 and S3 provide the splice-site census and the AT–AC 3′ splice-site results. Per-site tables of the HeLa allele analyses carry genotype-linked information and are not deposited, in keeping with the NIH HeLa Genome Data Use Agreement; their pooled values are included in the Dryad deposit, and the code deposit contains the scripts that rebuild them from the public HeLa GLORI reads.
● All original code has been deposited at Zenodo and is publicly available at https://doi.org/10.5281/zenodo.22950231 as of the date of publication. The deposit includes the analysis scripts, the software environment and a script that regenerates every figure from the tables deposited at Dryad.
● Any additional information required to reanalyze the data reported in this paper is available from the lead contact upon request.

## Acknowledgments

This work was supported by National Institutes of Health grant R35 GM130370 from the National Institute of General Medical Sciences to G.F.C.

## Author contributions

G.F.C. and K.R.R. conceived the study. K.R.R. performed the analyses. K.R.R. and G.F.C. wrote the manuscript.

## Declaration of generative AI and AI-assisted technologies in the manuscript preparation process

During the preparation of this work, K.R.R. used Claude Opus 5, Opus 5.5, Fable 5 and Fable 5.1 (Anthropic) to draft and revise the text and to write the code for the analyses and figures, including the Figure 6 schematic. After using these tools, the authors reviewed and edited the content as needed and take full responsibility for the content of the published article.

## Supplemental information

Document S1. Figures S1–S16, Supplemental Notes 1–4 and supplemental references

Table S1. 3′ splice-site compatibility census, related to Figure 1

Table S2. Studies, datasets and resources analyzed in this study, related to STAR Methods

Table S3. AT–AC 3′ splice-site methylation, writer dependence, the mouse *Tmem231* ortholog and the proposed METTL16 3′ splice-site targets, related to Figures 2 and 3

## STAR Methods

### Key resources table

| REAGENT or RESOURCE | SOURCE | IDENTIFIER |
| --- | --- | --- |
| **Deposited data** |  |  |
| Derived per-site tables, including the pooled three-flow-cell DRS table | This paper | Dryad:<br>10.5061/dryad.xpnvx0kz4 |
| Chromatin-associated RNA GLORI, A549 cells (METTL3 knockdown rescued with wild-type or 3KR METTL3) | Huang et al.<sup>22</sup> | GEO: GSE272467 |
| GLORI and unconverted (GLORI-) libraries of HEK293T, HeLa and MEF cells | Liu et al.<sup>21</sup> | GEO: GSE210563 (including the MEF GLORI- control GSM6432603) |
| m<sup>6</sup>A-SAC-seq, HeLa and other cell lines | Ge et al.<sup>32</sup> | GEO: GSE198246 |
| eTAM-seq | Xiao et al.<sup>52</sup> | GEO: GSE211303 |
| *S. cerevisiae* direct RNA sequencing, sporulating wild-type and *ime4Δ* cells | Leger et al.<sup>53</sup> | ENA: PRJEB44511 |
| RNA004 curlcake synthetic m<sup>6</sup>A standards (direct RNA, POD5) | Cruciani et al.<sup>54</sup> | ENA: PRJEB82528 (runs ERR13956435 unmodified, ERR13956436 m<sup>6</sup>A) |
| GM12878 whole-transcriptome in vitro transcribed RNA, direct RNA with depositor modification calls | European Nucleotide Archive (no linked publication) | ENA: PRJEB102644 |
| Native HEK293 and GM12878 direct RNA sequencing (RNA004) with modified-base BAMs | Ghohabi Esfahani et al.<sup>30</sup> | ENA: PRJEB111137 (HEK293 runs ERR17033219, ERR17033221, ERR17033222; GM12878 run ERR17033220) |
| TNT-seq m<sup>6</sup>A peaks in nascent RNA | Louloupi et al.<sup>55</sup> | GEO: GSE83561 |
| m<sup>6</sup>A-CLIP | Ke et al.<sup>3</sup> | GEO: GSE86336 |
| m<sup>6</sup>A-SAC-seq | Hu et al.<sup>31</sup> | GEO: GSE162340 (SuperSeries GSE162357) |
| m<sup>6</sup>A-SAC-seq | Hu et al.<sup>31</sup> | GEO: GSE162356 (SuperSeries GSE162357) |
| METTL16-AID degron GLORI, HCT116 cells | Flaherty et al.<sup>25</sup> | GEO: GSE296342 |
| HeLa siRNA RNA-seq panel (305 splicing factors and regulators) | Rogalska et al.<sup>56</sup> | ENA: ERP133481 (PRJEB49033; ArrayExpress E-MTAB-11202) |
| GLORI of activated human CD8 T cells (deposited alignments) | Gameiro et al.<sup>34</sup> | ArrayExpress: E-MTAB-15649 |
| miCLIP of activated human CD8 T cells (inspected, not reanalyzed) | Gameiro et al.<sup>34</sup> | ArrayExpress: E-MTAB-15643 |
| GLORI of bladder tumor, adjacent tissue and bladder cancer cell lines | Koch et al.<sup>26</sup> | GEO: GSE281749 |
| s<sup>4</sup>U-GLORI of HeLa nascent poly(A)-, nascent poly(A)+ and steady-state poly(A)+ RNA (reads) | Tang et al.<sup>23</sup> | GSA-human: HRA006964 (runs HRR1664501–HRR1664506) |
| s<sup>4</sup>U-GLORI site table (second supplementary table of the paper) | Tang et al.<sup>23</sup> | https://doi.org/10.1016/j.molcel.2024.07.017 (Table S2) |
| GLORI 2.0 and GLORI 3.0, HEK293T and mouse dorsal hippocampus | Sun et al.<sup>29</sup> | GEO: GSE270643 |
| Uli-Epic GLORI | He et al.<sup>57</sup> | GEO: GSE277359 |
| FOCAS GLORI, control-guide libraries of dCas13b-FTO cells | Zhang et al.<sup>27</sup> | GEO: GSE301419 |
| GLORI, A549, H9 and Jurkat cells | Huang et al.<sup>28</sup> | GEO: GSE290408 |
| m<sup>6</sup>A-SAC-seq | Dou et al.<sup>58</sup> | GEO: GSE205710 |
| m<sup>6</sup>A-SAC-seq | Huang et al.<sup>59</sup> | GEO: GSE280563 |
| m<sup>6</sup>A-SAC-seq | Mishra et al.<sup>60</sup> | GEO: GSE273614 |
| DART-seq | Tegowski et al.<sup>61</sup> | GEO: GSE180954 |
| miCLIP | Cho et al.<sup>62</sup> | GEO: GSE152633 |
| m<sup>6</sup>A-eCLIP | Alasar et al.<sup>63</sup> | GEO: GSE188580 |
| m<sup>6</sup>A-eCLIP | Akçaöz-Alasar et al.<sup>64</sup> | GEO: GSE223209 |
| DART-seq variants | Zhu et al.<sup>65</sup> | GEO: GSE196416 |
| miCLIP | Yang et al.<sup>66</sup> | GEO: GSE134103 |
| miCLIP | Huang et al.<sup>67</sup> | GEO: GSE121942 |
| m<sup>6</sup>A-CLIP | Ke et al.<sup>68</sup> | GEO: GSE71154 |
| meCLIP | Roberts et al.<sup>69</sup> | GEO: GSE147440 |
| m<sup>6</sup>A-SAC-seq, rice | Wang et al.<sup>18</sup> | GEO: GSE243722 |
| m<sup>6</sup>A-SAC-seq, *Arabidopsis* | Wang et al.<sup>18</sup> | GEO: GSE245738 |
| m<sup>6</sup>A-SAC-seq, mouse | Koo et al.<sup>70</sup> | GEO: GSE277229 |
| GLORI (examined, excluded from the 3'SS tests) | Gene Expression Omnibus (no linked publication) | GEO: GSE333647 |
| *S. cerevisiae* direct RNA, in vitro polyadenylated (coverage proxy only) | Gaglione et al.<sup>71</sup> | BioProject: PRJNA1365163 |
| IVM-seq: in vitro methylation of randomized substrates by METTL3–METTL14 | Zhang et al.<sup>20</sup> | Dryad: 10.5061/dryad.37pvmcvvb |
| HG002 direct RNA sequencing (two RNA004 poly(A) runs and eight SQK-DRB004 libraries) | Oxford Nanopore Technologies open data | Dataset UHRR_HG002_2026.06; https://ont-open-data.s3.amazonaws.com/UHRR_HG002_2026.06/ |
| HEK293 whole-genome sequence (Complete Genomics), parental line | Lin et al.<sup>72</sup> | ENA: ERR163011 |
| GIAB HG002 small-variant benchmark v4.2.1 (GRCh38) | Wagner et al.<sup>73</sup> | https://ftp-trace.ncbi.nlm.nih.gov/ReferenceSamples/giab/release/AshkenazimTrio/HG002_NA24385_son/NISTv4.2.1/GRCh38 |
| GIAB HG001 (GM12878) small-variant benchmark v4.2.1 (GRCh38) | Genome in a Bottle Consortium | https://ftp-trace.ncbi.nlm.nih.gov/ReferenceSamples/giab/release/NA12878_HG001/NISTv4.2.1/GRCh38 |
| gnomAD v4.1.1 genomes (76,215 individuals, FILTER PASS) | Chen et al.<sup>74</sup> | https://storage.googleapis.com/gcp-public-data--gnomad/release/4.1.1/vcf/genomes/; RRID:SCR_014964 |
| ENCODE eCLIP peaks, HepG2 and K562 | Van Nostrand et al.<sup>75</sup> | ENCODE portal, https://www.encodeproject.org (eCLIP, HepG2 and K562); RRID:SCR_015482 |
| ENCODE shRNA RNA-seq, RBM15 knockdown | Van Nostrand et al.<sup>75</sup> | ENCODE: ENCSR385UPQ (K562), ENCSR599PXD (HepG2) |
| ENCODE shRNA RNA-seq, FTO knockdown | Van Nostrand et al.<sup>75</sup> | ENCODE: ENCSR688GVV (K562), ENCSR389HFU (HepG2) |
| ENCODE shRNA RNA-seq, U2AF1 knockdown | Van Nostrand et al.<sup>75</sup> | ENCODE: ENCSR342EDG (K562), ENCSR372UWV (HepG2) |
| ENCODE shRNA RNA-seq, non-targeting controls | Van Nostrand et al.<sup>75</sup> | ENCODE: ENCSR067GHD, ENCSR264TUE, ENCSR585KOJ (HepG2); ENCSR143COQ, ENCSR344XID, ENCSR424QCW (K562) |
| ENCODE CRISPR RNA-seq, METTL3 guide | ENCODE Project Consortium (no linked publication) | ENCODE: ENCSR335KFN (K562), ENCSR884REL (HepG2) |
| ENCODE CRISPR RNA-seq, RNPC3 guide | ENCODE Project Consortium (no linked publication) | ENCODE: ENCSR631NIZ (K562) |
| ENCODE CRISPR RNA-seq, non-targeting guide controls | ENCODE Project Consortium (no linked publication) | ENCODE: ENCSR347NKX, ENCSR877JEM (K562); ENCSR704VPL (HepG2) |
| Human branch points mapped from lariat reads | Mercer et al.<sup>35</sup> | https://doi.org/10.1101/gr.182899.114 |
| Human branch points mapped from lariat intermediates | Zeng et al.<sup>36</sup> | https://doi.org/10.1016/j.molcel.2022.11.004 |
| Mouse branch points mapped from lariat reads (mm9; Tables S3 and S7 of the paper) | Taggart et al.<sup>76</sup> | https://doi.org/10.1101/gr.202820.115 |
| Minor Intron Database (MIDB) | Olthof et al.<sup>33</sup> | midb.pnb.uconn.edu (gene queries 2026-08-25; nine-species download 2026-09-03) |
| m<sup>6</sup>A-Atlas v2.0, human hg38 high-confidence sites | Liang et al.<sup>77</sup> | http://www.rnamd.org/m6a/download/high/hg38/hg38_CL_Tech.txt.gz |
| m<sup>6</sup>AConquer, human orthogonally validated sites (hg38) | Zhao et al.<sup>78</sup> | http://180.208.58.19/m6aconquer/Download/Human/HC_Overall/m6A_OrthogonallyValidatedSites_Combined_hg38.csv |
| DirectRMDb, *Homo sapiens* site table | Zhang et al.<sup>79</sup> | http://www.rnamd.org/directRMDb/download/HomoSapiens.zip |
| RMBase v3.0, human hg38 m<sup>6</sup>A sites | Xuan et al.<sup>80</sup> | http://bioinformaticsscience.cn/rmbase/download/modSingleSiteFiles/hg38/hg38.m6A.tar.gz |
| Structure: METTL3–METTL14 methyltransferase domain complex with SAH | Wang et al.<sup>43</sup> | PDB: 5IL2 |
| Structure: *S. pombe* U2AF1–RNA complex | Yoshida et al.<sup>7</sup> | PDB: 7Co6 |
| Structure: *S. pombe* U2AF1–RNA complex | Yoshida et al.<sup>7</sup> | PDB: 7Co7 |
| Structure: *S. pombe* U2AF1–RNA complex | Yoshida et al.<sup>7</sup> | PDB: 7Co8 |
| Structure: Postcatalytic spliceosome | Wilkinson et al.<sup>47</sup> | PDB: 6EXN |
| Structure: Postcatalytic spliceosome (P complex) | Liu et al.<sup>81</sup> | PDB: 6BK8 |
| Structure: Human postcatalytic spliceosome | Fica et al.<sup>82</sup> | PDB: 6QDV |
| Structure: Spliceosome after branching (catalytic step I) | Wan et al.<sup>46</sup> | PDB: 5GMK |
| Structure: Spliceosome after branching | Galej et al.<sup>83</sup> | PDB: 5LJ3 |
| **Software and algorithms** |  |  |
| Analysis and figure-generation code, software environment and figure regeneration script | This paper | Zenodo:<br>10.5281/zenodo.22950231 |
| GLORI-tools | Liu et al.<sup>21</sup> | https://github.com/liucongcas/GLORI-tools; commit def608b (def608b66ce4a32d18d76b4742df59dd3aaaa5c7, patched for current Python) |
| cutadapt | Martin<sup>84</sup> | Version 5.2; RRID:SCR_011841 |
| NGmerge | Gaspar<sup>85</sup> | Bioconda package 0.3 (binary reports version 0.2) |
| STAR | Dobin et al.<sup>86</sup> | Version 2.7.10b;<br>RRID:SCR_004463 |
| Bowtie (GLORI-tools transcriptome step) | Langmead et al.<sup>87</sup> | Version 1.3.1;<br>RRID:SCR_005476 |
| samtools | Danecek et al.<sup>88</sup> | Version 1.16.1 (version 1.23.1 for the curlcake caller validation);<br>RRID:SCR_002105 |
| BCFtools | Danecek et al.<sup>88</sup> | Version 1.16;<br>RRID:SCR_005227 |
| minimap2 | Li<sup>89</sup> | Version 2.28 (yeast); 2.30-r1287 (GM12878 and curlcake alignments; depositor alignments of PRJEB111137 and of UHRR_HG002_2026.06);<br>RRID:SCR_018550 |
| Dorado | Oxford Nanopore Technologies | https://github.com/nanoporetech/dorado; 2.1.1 with rna004_130bps_hac@v5.1.0 and inosine_m6A@v1 or m6A_DRACH@v1 (curlcake standards, this study); depositor basecalls: 1.0.0 sup with the 8-modification all-context model (PRJEB111137 HEK293, one flow cell), sup@v5.2.0 (PRJEB111137 GM12878), 2.0.0 sup@v6.0.0 with inosine_m6A_2OmeA@v1 (HG002), 0.8.1 four-modification model (PRJEB102644) |
| f5c | Gamaarachchi et al.<sup>90</sup> | Version 1.4 |
| Nanocompore | Leger et al.<sup>53</sup> | Version 1.0.4 |
| pyliftover | N/A (PyPI package) | https://pypi.org/project/pyliftover/; version 0.4.1 |
| MaxEntScan, Python implementation maxentpy | Yeo and Burge<sup>91</sup> | https://github.com/kepbod/maxentpy (master branch archive) |
| ViennaRNA (RNAplfold) | Lorenz et al.<sup>92</sup> | Version 2.5.1;<br>RRID:SCR_008550 |
| PyMOL (open-source) | Schrödinger, LLC | Version 3.1.0;<br>RRID:SCR_000305 |
| Python | Python Software Foundation | Version 3.11.15 (figures and tables); 3.9.23 (native-register readout); per-step versions of upstream steps in the code deposit STEPS.tsv;<br>RRID:SCR_008394 |
| NumPy | Harris et al.<sup>93</sup> | Version 2.4.3 (figures); 2.0.2 (native-register readout);<br>RRID:SCR_008633 |
| SciPy | Virtanen et al.<sup>94</sup> | Version 1.17.1 (figures); 1.13.1 (native-register readout);<br>RRID:SCR_008058 |
| Matplotlib | Hunter<sup>95</sup> | Version 3.10.8;<br>RRID:SCR_008624 |
| pandas | McKinney<sup>96</sup> | Version 2.3.3 (figures); per-step versions in STEPS.tsv;<br>RRID:SCR_018214 |
| pysam | N/A (GitHub: pysam-developers) | https://github.com/pysam-developers/pysam; versions 0.19.0, 0.22.1 and 0.23.3 by step |
| statsmodels | Seabold and Perktold<sup>97</sup> | Versions 0.14.5 and 0.15.0 by step |
| RECTIFY (plotting module only: gene models of the isoform-ambiguity figure) | This lab | https://github.com/k-roy/RECTIFY; tag manuscript/m6a-3ss-20260924 |
| Claude Code (Claude Opus 5, Opus 5.5, Fable 5 and Fable 5.1) | Anthropic | https://www.anthropic.com/claude-code |
| **Other** |  |  |
| Human genome GRCh38, primary assembly (GENCODE release 46 FASTA) | Genome Reference Consortium; GENCODE | https://ftp.ebi.ac.uk/pub/databases/gencode/Gencode_human/release_46/ (GRCh38.primary_assembly.genome.fa); RRID:SCR_014966 |
| Human genome GRCh38 analysis set without alternate contigs | Genome Reference Consortium (NCBI) | GenBank assembly GCA_000001405.15 (no-alt analysis set) |
| GENCODE human annotation, release 46 | GENCODE | https://ftp.ebi.ac.uk/pub/databases/genocode/Gencode_human/release_46/;<br>RRID:SCR_014966 |
| GENCODE human annotation and transcripts, release 29 (GLORI-tools reference) | GENCODE | https://ftp.ebi.ac.uk/pub/databases/genocode/Gencode_human/release_29 |
| Mouse genome GRCm38 (mm10) primary assembly with GENCODE mouse annotation release M25 | GENCODE | https://ftp.ebi.ac.uk/pub/databases/genocode/Gencode_mouse/release_M25 |
| Mouse genome GRCm39 with GENCODE mouse annotation release M35 (splice-site census of Table S1) | GENCODE | https://ftp.ebi.ac.uk/pub/databases/genocode/Gencode_mouse/release_M35/ |
| Ensembl release 115 and Ensembl Genomes release 62 genomes and protein-coding annotations (splice-site census of Table S1) | Ensembl | https://ftp.ensembl.org/pub/release-115;<br>[https://ftp.ensemblgenomes.ebi.ac.uk/pub (release 62)](https://ftp.ensemblgenomes.ebi.ac.uk/pub/release_62/);<br>RRID:SCR_002344 |
| Ensembl release 97 genome sequences (nine-species U12 census with MIDB introns) | Ensembl | Ensembl release 97;<br>RRID:SCR_002344 |
| *S. cerevisiae* reference genome R64, as Ensembl R64-1-1 (splice-site census of Table S1) and SGD R64-5-1 (yeast direct RNA comparison) | Ensembl | Ensembl assembly R64-1-1; SGD release R64-5-1;<br>RRID:SCR_002344 |
| MANE Select transcripts | MANE_Select tag of the GENCODE release 46 annotation | GENCODE release 46 GTF, tag MANE_Select |
| UCSC liftOver chain hg18ToHg38 | UCSC Genome Browser | https://hgdownload.soe.ucsc.edu/goldenPath/hg18/liftOver/hg18ToHg38.over.chain.gz |
| UCSC liftOver chain mm9ToMm10 | UCSC Genome Browser | https://hgdownload.soe.ucsc.edu/goldenPath/mm9/liftOver/mm9ToMm10.over.chain.gz |
| UCSC liftOver chain hg19ToHg38 (lifting deposited site tables and database records) | UCSC Genome Browser | https://hgdownload.soe.ucsc.edu/goldenPath/hg19/liftOver/hg19ToHg38.over.chain.gz |

### Experimental model and study participant details

This study generated no new data from cell lines, animals, plants, microbes or human participants, and all data were obtained from public repositories and are listed with their accessions in the key resources table and Table S2. The principal human cell lines in the reanalyzed data were HEK293 (female; RRID:CVCL_0045), HEK293T (female; RRID:CVCL_0063), HeLa (female; RRID:CVCL_0030), A549 (male; RRID:CVCL_0023), HCT116 (male; RRID:CVCL_0291), the lymphoblastoid cell lines GM12878 (female; RRID:CVCL_7526) and HG002 (GM24385, male; RRID:CVCL_1C78), and the bladder cancer cell lines T24 (female; RRID:CVCL_0554), RT4 (male; RRID:CVCL_0036) and UM-UC-3 (male; RRID:CVCL_1783), with sexes as recorded in Cellosaurus. Primary human CD8 T cells^34^ and bladder tumor and adjacent tissue^26^ were analyzed only as the sequencing data and site tables deposited by those studies, and donor characteristics are those reported there. Mouse data came from embryonic fibroblasts^21^ and dorsal hippocampus,^29^ whose deposits do not record sex, strain or age, and further mouse, rice and *Arabidopsis* site tables and the *S. cerevisiae* direct RNA data of sporulating wild-type and *ime4*Δ cells^53^ are listed in Table S2. Because every comparison in this study is made between positions or alleles within one sample or dataset, sex was not analyzed as a variable, and the conclusions are not tested for differences between female and male cells.

## Method details

### Genome annotations and splice-site census

The 35-genome census (Table S1A) used Ensembl protein-coding annotations and included 2,497,337 GT–AG 3′SS A residues, scored separately from GC–AG 3′ splice sites (Figure 1B). Human and mouse AT–AC censuses and methylation analyses, intron models and exon-start composition used GENCODE v46 (hg38) and vM25 (mm10). U12-type classifications and minor-intron-containing genes came from the Minor Intron Database (midb.pnb.uconn.edu, gene queries 2026-08-25),^33^ and a separate download on 2026-09-03 supplied the nine-species U12 branch-point census. The seven-species census of Table S1B used GENCODE vM35 on GRCm39 for mouse, Ensembl release 115 and Ensembl Genomes release 62 for the other non-human species, and Ensembl R64-1-1 for *S. cerevisiae*. Accessions and genome builds are in Table S2. Base composition at each position around the 3′SS A was tabulated over the 250,300 human and 220,592 mouse GT–AG 3′ splice sites of Table S1B.

We defined the 3′SS A as the A in the terminal AG or AC and extracted the five nucleotides centered on it in transcript orientation. We scored compatibility with DRACH (D = A/G/U, R = A/G, H = A/C/U), the stricter RRACH and the fungal RGAC consensus, counting a position as compatible only when every specified base matched, without partial or weighted matching.

To test for depletion of writer motifs near splice sites, we counted motifs in 60-nt windows and compared them with dinucleotide-preserving shuffles of up to 500 nt of the intron’s 3′ end and up to 300 nt of the downstream exon, each shuffled separately, and with a column permutation of the window, and report the range spanned by the two null models. Figure 4F shows RGAC from −55 to −4 nt against the dinucleotide-preserving shuffle alone, with whiskers of one standard deviation of that null; against the column permutation the seven ratios range from 0.57 to 1.04, with 0.91 in human and 1.04 in mouse.

### In vitro specificity

We reanalyzed randomized-substrate IVM-seq data deposited at Dryad under 10.5061/dryad.37pvmcvvb.^20^ The published protocol describes an N20 substrate; in the released Abcam and Synaptic Systems libraries, we analyzed 20- and 23-nt inserts, respectively, between the documented constant flanks. The mock libraries are antibody-free controls,^20^ so each wild-type/mock ratio compares antibody-selected with unselected material from the same methylation reaction. Two antibodies that differ only in capture efficiency give enrichment grids related by a single constant.

Fitting (Synaptic Systems − 1) = k × (Abcam − 1) through the origin over all 16 combinations of −1 and +1 bases gave k = 0.125 with a residual standard deviation of 0.020, and the two pools therefore differ in dynamic range rather than in sequence preference. The Synaptic Systems libraries prepared with the METTL14 R298P enzyme, a cancer-derived mutant that redirects the writer toward GGAU rather than reducing its activity,^20^ showed no −1 preference (0.99, 0.95, 0.97 and 1.00 for A, C, G and U; Figure S1C). The two antibody pools ranked the contexts concordantly, which does not exclude a capture bias shared by both. The −1 values of Figure 1E are raw ratios of the antibody-selected library to its no-antibody control, normalized for library size so that 1 denotes no enrichment.

The single-A analysis retained reads with exactly one A in the central window (Figures S13A and S13B). Enrichment was the frequency of a context in the wild-type library divided by its frequency in the corresponding deposited mock library. Contexts the enzyme does not use share one baseline ratio, which we estimated as the median ratio of the six −1/+1 combinations with G or U at +1, 0.87 in the Abcam pool and 0.97 in the Synaptic Systems pool; for comparison, the raw GGAC ratios were 3.04 and 1.44, and those of GGAG, which carries G in place of the +1 C, 1.00 and 0.89. The score subtracts that baseline from each ratio and divides by one minus the baseline, so that a context at the baseline scores 0 and a context with the pool-average enrichment scores 1; on this scale GGAC scored 16.3 in the Abcam pool and 14.2 in the Synaptic Systems pool, and the YAG contexts 0.03 and −0.01. Ranked on this score without reference to any motif, the three single-A GGAC 5-mers placed among the top four of the 81 single-A 5-mers in both pools (Figure S1D). This background-subtracted enrichment score is not a calibrated methylated fraction or catalytic activity. Reported 95 % confidence intervals propagate count uncertainty and treat the estimated background as fixed. The full-pool analysis used all reads, deduplicated them, and ranked the 256 A-centered 5-mers by wild-type-over-mock enrichment (Figure S1E). The second readout was used for the 3′SS register of Figure 1F; because every A-centered 5-mer on a selected molecule gains a count, it does not measure the boundary-spanning register, whose 5-mers carry a second A at −2 (Figures S13A, S13C, and S13D). A method-of-moments fit over all 256 A-centered contexts, which apportions the counts of each selected molecule among the 5-mers it carries, estimates 81 of them (Figure S13E).

The mock libraries were prepared by the same procedure as the antibody pulldowns but incubated with beads in the absence of antibody.^20^ Because enrichment is a ratio of frequencies, its count-weighted mean over all contexts in a pool is 1 by construction. The context with G at −1 and C at +1, whose Abcam enrichment lies 4.7 times as far from 1 as that of any other context, carries the leverage of the single-constant fit between the two antibody pools. Its residual was +0.41 standard deviations, and the most enriched and most depleted contexts compressed by similar factors (0.134 and 0.120). Because YAG contexts contribute to the background estimate, we also recalculated the background without them, and a pyrimidine at −1 then removed 91.6 % (Abcam) and 96.1 % (Synaptic Systems) of the GGAC score.

Antibody selection returns whole molecules, so enrichment shows that a molecule carried at least one m^6^A but not which of its A residues was methylated (Figure S13A). A random region usually carries several A residues, and every A-centered 5-mer on a selected molecule gains a count, including 5-mers that the enzyme does not use, whereas a read with a single A in the random region leaves one candidate (Figure S13B). The single-A analysis of Figure 1D therefore uses only such reads, which can contain only the four DRACH 5-mers without a second A (GGACC, GGACU, UGACC and UGACU). To use all reads, we modeled the chance that a molecule is selected as an antibody-specific background plus one contribution from each A-centered 5-mer it carries, and we estimated the contributions of all 256 5-mers in each antibody pool by the method of moments; the contributions were additive in these data. The fit separates 5-mers that occur in different combinations on different molecules, but an A at −1 or −2 creates a second A-centered window that overlaps the first on every molecule, so the fit cannot separate those 5-mers from their neighbors (Figure S13C). This applies to 12 of the 18 DRACH 5-mers, and two more, GGACA and UGACA, are separable in principle but have no valid estimate, which leaves four that are measured (Figure S13D). Across the 81 5-mers that the fit can estimate, GGACG, GGACU and GGACC ranked first to third with both antibodies, and the two pools agreed (Pearson r = 0.93; Figure S13E). Full-pool enrichment of a 5-mer with an A at −1 or −2 therefore does not measure that 5-mer’s own activity.

### Single-base m^6^A maps

Published site sets were analyzed as deposited unless a read-level reanalysis is specified. The GLORI datasets included GSE272467 (A549 chromatin-associated RNA from METTL3-depleted cells rescued with wild-type METTL3) and GSE210563, the latter covering polyadenylated RNA from HEK293T, HeLa and MEF, including HeLa siNC, siMETTL3, siMETTL14, siWTAP, siYTHDF2 and STM2457 conditions. For GSE272467, whose libraries carry no unique molecular identifiers, we used the sites called in both wild-type replicates, as Huang et al.^22^ did, and counted an intron as covered when each replicate called at least one site in it. Counted the same way in the HEK293T and HeLa libraries of GSE210563, the canonical 3′SS A was called at 31 of 24,320 and 27 of 16,540 positions, against 185 of 2,534 and 112 of 1,793 DRACH adenosines 3 to 15 nt upstream in the same introns (57- and 38-fold) and 80 of 146,135 and 76 of 99,675 YAG adenosines 3 to 200 nt upstream (2.33-fold, 95 % CI 1.54 to 3.52, and 2.14-fold, 1.38 to 3.32); against YAG adenosines 3 to 50 nt upstream the ratios were 1.62 (0.90 to 2.93) and 1.41 (0.77 to 2.59). A broader 29-series collection combined chemical mapping, crosslinking and editing assays; the 25 motif-unrestricted GLORI libraries of the cross-deposit census were GSE281749,^26^ GSE301419 (control-guide libraries of cells that express dCas13b-FTO),^27^ GSE290408 (sites already filtered by the depositors against METTL3 knockdown),^28^ GSE270643^29^ and GSE277359.^57^ The m^6^A-SAC-seq datasets included GSE162340 and GSE162356,^31^ GSE198246,^32^ GSE205710^58^ and GSE277229,^70^ and the rice and *Arabidopsis* atlases GSE243722 and GSE245738;^18^ the eTAM-seq dataset was GSE211303.^52^ Additional datasets were E-MTAB-15649, analyzed from the deposited alignments because no site table was released, GSE280563,^59^ GSE273614^60^ and GSE147440.^69^ GSE333647 was examined but excluded from the 3′SS tests, because only 20 to 23 % of its called sites are in DRACH, against 66 to 89 % in every other GLORI methylation map except the HEK293T map of GSE277359 (34 %). The companion miCLIP deposit E-MTAB-15643 was inspected but not re-analyzed because no site table was available. The METTL16-AID degron analysis used GSE296342. Non-human maps came from mouse, rice, *Arabidopsis* and *S. cerevisiae*; the nine-species comparison elsewhere in this work is a sequence census, not a methylation map. Nascent-RNA sites came from the s^4^U-GLORI site table published as the second supplementary table of Tang et al.^23^, which reports m^6^A levels for newly synthesized and steady-state chromatin-associated HeLa RNA after a 5-minute s^4^U pulse; the GSA-human deposit HRA006964 holds the reads. Its coordinates are zero-based and were converted before use, with the frame confirmed by requiring an A at every listed position on the transcribed strand. That table lists detected sites only, with no covered-but-unmethylated positions, so it was used to ask which splice-site A residues are detected. Table S2 lists platforms, cell types and treatments for the principal datasets; Table S3E records the expanded AT–AC signals individually.

When a deposit reported per-site counts, a detected site met the deposit’s own significance threshold. The GLORI site tables used for the AT–AC analysis required A+G coverage ≥ 15, A coverage ≥ 5 and a methylation ratio ≥ 0.10 in addition to that threshold.

An m^6^A-CLIP peak was scored as intronic only if it overlapped no annotated exonic region, so the intronic share shown for chromatin-associated RNA is inferred from the reported exonic fraction (Figures S3A and S3B).^3^ The four de novo motifs that Louloupi et al.^55^ derived from m^6^A peaks in nascent RNA were scored on the single-A substrates of the in vitro analysis (Figure S3C).

To recover the per-available-A denominator, every read pair of the six s^4^U-GLORI HeLa runs (GSA-human HRA006964, HRR1664501–HRR1664506: nascent poly(A)−, nascent poly(A)+ and steady-state poly(A)+, two replicates each) was trimmed with cutadapt 5.2,^84^ stitched with NGmerge (bioconda package 0.3, whose binary reports version 0.2) and processed through the authors’ GLORI-tools pipeline (commit def608b, patched to run under current Python; STAR 2.7.10b against the GRCh38 primary assembly with GENCODE v29 transcripts, at most two mismatches and unique mapping, with the transcriptome rescue step). At each reference A, reads reading A (methylated) and G (converted) were counted strand-aware at base quality ≥ 10, keeping reads with at most three A residues remaining; a site was covered at A+G ≥ 15, the rate is the fraction reading A, A/(A+G), and pooled rates sum A over A+G. Positions scored were the 302,110 canonical 3′SS A residues with offsets −500 to +500, the 5′SS A3 and A4 residues of GENCODE v46 introns and the 58,259 mapped branch-point A residues of the lariat-sequencing set, each classified by its own 5-mer, and AT–AC 3′SS A residues were scored from a ±200 nt window. 5′SS A4 and branch-point positions that lie inside an exon of any same-strand GENCODE v46 transcript (22 % and 25 % of the DRACH-compatible positions) were excluded from the per-site and direct-RNA class rates, because methylation there cannot be assigned to the unspliced precursor; rates over all annotated positions are given in the Figure 4 legend. Each library’s background is the median gene-level A fraction reported by GLORI-tools for that library on chromosomes 1 and 22 (1.7–3.1 %). Because a spliced read whose upstream exon ends in the same bases as the intron can align across the last intron positions, we also recomputed the 3′SS A level on reads that continue at least 20 nt into the intron, which gave 1.29 to 1.52 %, still below every library’s background. Removing duplicate reads by fragment start, end and strand left the 3′SS A below background in every library. Sites were also called with criteria adapted from the published ones (at least 15 reads, where the published calls required 30, and 5 reads reading A, level at least 0.10, one-sided binomial P below 0.005 against the median gene-level A fraction; the published RNA-coverage, base-quality and homopolymer filters were not applied); agreement with the published levels at the deposited sites was the acceptance gate. For the intronic share and the per-region rates, every covered A (A+G ≥ 15) on chromosomes 1 and 22 was annotated as exonic, intronic or intergenic on the transcribed strand against GENCODE v29. Each covered site was tested against its library’s background with a one-sided binomial test, P values were corrected within each library by the Benjamini–Hochberg procedure, a site was elevated at q < 0.05 and level ≥ 10 %, and reproducible when elevated in both replicates (Figure S9B). DRACH-context and non-DRACH A residues of the same position were compared as elevated-site rates per covered site–library observation and per unique position, with Fisher’s exact test and log-scale 95 % confidence intervals for the rate ratio.

Mouse GLORI+ libraries were realigned in the same way: the four MEF libraries of GSE210563 (heat shock plus and minus, two replicates each) and the eight dorsal-hippocampus libraries of GSE270643 (cytoplasmic and synaptic fractions, four each) were trimmed, aligned through the GLORI-tools pipeline against GRCm38 with GENCODE vM25, and counted at every A within ±500 nt of canonical 3′ splice sites, classified as GT–AG or GC–AG by their terminal dinucleotides, or as AT–AC when an intron of a protein-coding gene begins with AT and ends with AC, at the 5′SS +3 and +4 positions, at AT–AC 3′SS A residues confirmed in the annotation, and at 6,509 mouse branch-point A residues mapped from lariat reads^76^ and lifted from mm9 to mm10 with pyliftover 0.4.1 and the UCSC mm9ToMm10 chain, of which 99.3 % reproduce the published context. A site was covered at A+G ≥ 15, each library’s background was its median gene-level A fraction on chromosomes 18 and 19 (1.4 to 2.6 %), and per-site elevation used the binomial test and Benjamini–Hochberg correction described above. At the 12 3′ splice sites proposed as METTL16 targets, a read counted as intron-containing only if its alignment continued at least 20 nt into the intron, because a spliced read whose upstream exon ends in the same bases as the intron aligns across the last intron positions. The level at canonical 3′SS adenosines was measured separately in one MEF and two hippocampus libraries, at 300 randomly drawn covered sites each with the intron classes pooled, on reads running at least 10 nt into the intron. Only 10 to 16 % of the reads counted at those sites were intron-containing, and those reads were unconverted in 0.43 to 0.98 %, below every library’s background. Per-library values are in Figures S5A–S5E. The libraries analyzed sample neither the gonads, where METTL16 is most abundant, nor the early embryos in which two of the proposed mouse 3′ splice sites responded to its loss, so methylation of those sites there remains possible.^5^

MazF-based methods, including MAZTER-seq^39^ and m^6^A-REF-seq,^98^ detect methylation only at ACA and cannot target the canonical 3′SS AG. DART-seq detects editing of cytidines near m^6^A.^99^ In m^6^A-CLIP the site within a peak is placed at the consensus motif carrying the crosslink-induced mutation or truncation signature,^3^ so a canonical 3′SS A, which lies outside RRACU, is never a candidate site. Published m^6^A-SAC-seq deposits are filtered to DRACH by the authors’ site criteria, so our analysis of that assay used reads rather than the deposited site set. Analysis of 3′SS methylation was accordingly based on GLORI and DRS nanopore measurements, which do not impose a motif requirement. Structure-dependent conversion artifacts are not controlled in the public GLORI data, nor are PCR duplicates in libraries without unique molecular identifiers, which is why the direct RNA evidence carries weight for the 3′SS and branch-point conclusions.

In the single-A stratum of Figure S3C, DSAGR and KSAGG specify the same four 5-mers and are shown together, and DGACH specifies the same 5-mers as DRACH. In Figures 2B and S4C, replicate measurements were averaged per locus, and for the METTL16 degron (Figure S4C) the loci were then paired across auxin states within each rescue condition. The metaprofiles of Figures 2A and 2C use bins of 1–10, 11–20, 21–50, 51–100, 101–200 and 201–500 nt from the 3′SS A, starting at +3 on the exon side with exon +1 drawn alone, on fixed populations, so that the same 3′ splice sites contribute at every offset. In chromatin-associated A549 RNA these are introns of at least 500 nt and downstream exons of at least 200 nt, and in the s^4^U-GLORI libraries introns and downstream exons of at least 500 nt each, with bins of fewer than 200 covered sites not drawn. Introns that are terminal in one transcript and interior in another form a third, mixed stratum. In the survey of Figure S6, reanalyses of the same deposit count once, and each of the two T-cell union tables counts as one entry. Covered-site measurements, deposits that failed reference-build or coordinate checks, and unfinished analyses were excluded from the site-list census.

### Native single-molecule direct RNA sequencing

Native HEK293 reads came from PRJEB111137,^30^ in which the HEK293 data comprise three RNA004 flow cells, and were analyzed from the deposited modified-base BAMs. The deposit was basecalled with the dorado sup model and the 8-modification all-context model; its file names record dorado 1.0.0 for one of the three flow cells and no version for the other two. Reads were aligned to the GRCh38 analysis set without alternate contigs (GCA_000001405.15) using minimap2 -t 64 -ax splice -uf -k14 -y. An all-context model is required because a DRACH-restricted model cannot score a 3′SS A in a YAG context. The dataset assayed whole-cell polyadenylated HEK293 RNA, in which intron-containing molecules are a selected subset.

A read-level methylation detection required an ML byte ≥ 250, a lower probability bound of 250/256 (approximately 0.977) on the interval [N/256, (N+1)/256) that a SAM ML byte N encodes. For each class, the pooled rate was the number of reads detected as modified at ML ≥ 250 (sensitivity to that threshold is described below) divided by the sum of aligned observations. This rate is a modified-read fraction, not a stoichiometry.

On PRJEB82528 curlcake standards,^54^ the all-context RNA004 model detected methylated A residues in YAG contexts with 57.0 % recall and a 1.6 % per-site false-positive rate at a detection-probability threshold of 0.8, below the ML ≥ 250 threshold (a probability of about 0.98) used for calls in this study; DRACH recall was 52.8 %. The whole-transcriptome in vitro transcribed control PRJEB102644 (GM12878) provided false-positive rates by 5-mer over 851 million A observations, with every A counted in the denominator. As a fraction of read observations at the threshold used here, false-positive rates were 0.127 % across all A residues, 0.076 % at DRACH A residues, 0.321 % in YAG context and 0.416 % in the CAG core of the 3′ splice site; the seven worst 5-mers all carried a pyrimidine at −1 and G at +1, six of them with C at −1. This control was basecalled by its depositors with Dorado 0.8.1 and a four-modification model of the same all-context family rather than the eight-modification HEK293 model, so it characterizes sequence dependence without setting a formal error bound. Comparisons in the text are made within one dataset between positions matched on 5-mer context. That restriction is necessary rather than cautious: across the 18 DRACH 5-mers the model’s false-positive rate on unmethylated RNA correlates with the in vivo modified share (Spearman rho 0.70), so the 5-mers most often methylated in cells are also those the model most often reports as modified in unmethylated RNA.

Native reads therefore cannot rank 5-mers by in vivo methylation independently of the caller. The 5-mer shares of Figure 4D serve only as the within-dataset expectation of Figure S10C and not as a ranking of in vivo methylation, and every writer-dependence result in this study rests on GLORI, which has no learned sequence prior.

The minimap2 options -ax splice, -uf and -y enable spliced alignment, fix the transcript strand and carry the MM and ML modification tags into the alignments. Requiring the whole probability bin of an ML byte to exceed 0.98 would need ML ≥ 251. In a sensitivity analysis on chromosome 21, ML ≥ 251 lowered the pooled rate by 3.7 % of its value over all positions and by 1.6 % at DRACH, removed 4 of the 14 detections at 3′SS A residues and raised the DRACH-to-3′SS A contrast from 45.9-fold to 63.2-fold, so ML ≥ 250 gives the more conservative contrast. In Figure S2B, each percentage divides the covered read–A observations with a reported m^6^A probability by all covered observations in that context. The emission threshold decides which probabilities are written to the BAM and differs from the detection thresholds of Figures S2A, S2C, and S2D. Sethi et al.^4^ applied the same Dorado all-context model family (sup inosine_m6A, RNA004 chemistry) to human chromatin-associated pre-mRNA.

### Classification of spliced and unspliced reads

Reads were classified at each annotated 3′SS from their alignment across the intron–exon boundary. Both classes required at least 25 aligned nucleotides upstream and 25 from exon +1 onward, with the upstream anchor intronic for unspliced reads and exonic for spliced reads. A continuous alignment across the boundary was classified as unspliced, and an N operation matching the annotated intron identified a spliced read. Unspliced observations were counted once per 3′SS and read. Editing was scored at each adenosine from basecalled G and from Dorado’s direct inosine calls. The splicing efficiency of an intron is the spliced share of all reads classified at its 3′SS in the same libraries as the methylation calls, and introns with fewer than 10 classified reads were not scored. 3′ splice sites whose alternative 5′ splice sites assigned different motif-register classes were excluded.

The analysis population for Figure S8D included all 3′ splice sites whose annotated 5′ splice sites agreed on motif class, and we fixed that population before examining the corrected interactions. Eligibility came from annotation; a site did not need observations in both splice states. We divided the unspliced-to-spliced detection-rate ratio for lost motifs by the corresponding ratio for retained motifs. Its 95 % interval used 2,000 joint bootstrap draws over gene components (seed 867), applying the same resampling weights to both motif classes. Genes sharing an annotated 3′SS formed one component, and each 3′SS contributed once. These intervals describe variation across gene components, not biological replicates.

Sensitivity analyses kept only 3′ splice sites with a single annotated 5′ splice site and then compared upstream exons ending in CG, where splicing removes the boundary-spanning motif, with those ending in AG, GG or UG, where it is retained. That comparison fixes the terminal base rather than full splice-site strength. Both the primary all-5′SS estimate and the single-5′SS estimate are reported. Bootstrap draws that gave zero or infinite ratios were retained, and any undefined draw would have prevented an interval. In the direct RNA reads of Figures S8D and S9A, modified reads over aligned observations were 2,073 of 71,100 in unspliced and 2,709 of 804,514 in spliced RNA for retained motifs, 403 of 8,647 and 90 of 129,210 for lost motifs, and 188 of 724,747 and 1,780 of 5,927,354 for 3′ splice sites without a boundary-spanning DRACH, a class defined by its unspliced context. The decrease from unspliced to spliced RNA is therefore 8.66-fold for retained motifs and 66.9-fold for lost motifs. For pooled class rates, we also calculated a depth-unweighted per-site mean, which in Figures S8D and S9A required at least 20 observations per site and splice state, and repeated each class without the gene that contributed the most signals. In the logistic models of Figure S9C, a term is omitted where a class has no events on one side. The eCLIP meta-profiles of Van Nostrand et al.^100^ place U2AF1 and U2AF2 at the 3′ splice site and SF3B4 and SF3A3 at the branch point.

### Isoform ambiguity in the intronic reference

A position that is intronic in one annotated transcript can lie inside an exon of another. A read that covers such a position and continues through the 3′ splice site without a gap is counted as unspliced, although it could come from a precursor of the first transcript or from mature RNA of the second, so part of the methylation scored as intronic could be exonic and the splicing efficiency of the intron would be underestimated. We therefore called a position ambiguous when it lies inside an exon of another same-strand GENCODE v46 transcript that is not itself a retained-intron annotation, since such an annotation records the very retention being measured, and we called an intron overlapped when an exon of that kind overlaps any part of it.

We kept every position of the intronic reference rather than removing the ambiguous ones, because removing them would clean the reference but not the splicing signals compared with it, and we added intron overlap to the matching of Figure S10C (Quantification and statistical analysis). Three tests asked whether reads at ambiguous positions come from precursors or from mature RNA of the other transcript. The first compared the density of exon–exon junctions on reads counted unspliced at introns that another transcript’s exon spans with that on unspliced reads at introns without overlap and on spliced reads at the same introns. The second compared the junction density of reads called modified at ambiguous positions with that of unmodified reads at the same positions, in subsets defined by the kind of overlapping exon, the distance from the other transcript’s stop codon, the geometry of the overlap and splicing efficiency, against the same difference at positions that no other transcript covers. The third compared methylation at ambiguous and unambiguous positions in the HeLa s^4^U-GLORI libraries, with splicing efficiency taken from the direct RNA reads of the same introns. The results are given in Supplemental Note 2 in Document S1.

### Exon +1 at covered 3′ splice sites

We called a 3′ splice site covered when one exon of another same-strand GENCODE v46 transcript, other than a retained-intron annotation, contains both its last intronic nucleotide and exon +1 (Figure S7A). In the covering transcript the intron’s AG still precedes the exon +1 adenosine, so a boundary-spanning DRACH is intact in its mature RNA, and a short read that runs across the 3′ splice site without a gap is counted as unspliced whether it comes from the precursor or from that mRNA. In the HEK293 direct RNA reads, which are long enough to tell the two apart, we classified a read as a precursor when it rules out the mature RNA of every covering transcript, by running at least 25 nt into one of that transcript’s introns or beyond its first or last exon, or by carrying a junction that the transcript lacks. We classified a read as the covering transcript’s mRNA when it fits that transcript’s exons and has removed part of the host intron by splicing, and we compared exon +1 methylation between the two classes at the same sites (Figure S7C). The intron-retention comparison of Figure S8E and the polypyrimidine-tract comparison of Figures S8A and S8B were repeated at uncovered 3′ splice sites (Figure S7D). The results are given in Supplemental Note 4 in Document S1.

### Allele-specific m^6^A at heterozygous variants

HEK293 genotypes came from Complete Genomics whole-genome sequencing of the parental HEK293 line (ENA ERR163011),^72^ lifted from NCBI36 to GRCh38 with pyliftover and the UCSC hg18ToHg38 chain. The reference base matched GRCh38 at 99.21 % of lifted positions, and 1,562,504 heterozygous single-nucleotide variants were kept. HEK293 methylation came from the direct RNA reads described above. HG002 direct RNA came from Oxford Nanopore open data (dataset UHRR_HG002_2026.06, two RNA004 poly(A) runs and eight SQK-DRB004 libraries), basecalled by the depositors with Dorado 2.0.0 sup@v6.0.0 and the inosine_m6A_2OmeA@v1 all-context model and aligned with minimap2 -ax splice -uf -k14 -y, with genotypes from the GIAB v4.2.1 GRCh38 benchmark (2,060,752 heterozygous variants inside benchmark regions).^73^ GM12878 direct RNA came from the fourth run of PRJEB111137 (ERR17033220, RNA004, basecalled by the depositors with Dorado sup@v5.2.0 and the all-context model used for HEK293), aligned with the same minimap2 options to the GENCODE release 46 GRCh38 primary assembly, with genotypes from the GIAB v4.2.1 HG001 benchmark; its single flow cell was split in two by read name for the held-out selection. Only primary alignments were used. In each read, the target adenosine had to lie inside one gapless aligned block with 25 nt on each side, the read base at the target had to be A in transcript orientation and the base at the variant one of its two alleles, and a call required ML ≥ 250. A site was eligible when heterozygous with no second variant in its 5-mer, covered by at least 10 reads of each allele over all runs, with at most 20 % of reads carrying neither allele, and with a minor-allele fraction of at least 0.15 in the RNA reads, which confirms heterozygosity in the sequenced stock. Variants at positions −2 to +2 at which exactly one allele formed DRACH were tested. Variants that kept DRACH on both alleles, and variants 3 to 5 nt from the adenosine, served as recall controls for the caller. Sites were selected on one group of runs (a pooled call rate of at least 10 % with at least 3 calls) and scored on the other, and the two folds were summed (HEK293, ERR17033222 against ERR17033219 and ERR17033221; HG002, the RNA004 runs against the SQK-DRB004 libraries). The criteria were fixed before the data were examined. At variants at −1 or +1, the ratio had to be at least 5, with a 95 % site-bootstrap interval (2,000 resamples) excluding 1, over at least 20 sites, and the recall-control intervals had to overlap 0.5 to 2. Excluding A to G variants in transcript orientation, which A-to-I editing can mimic, left the HEK293 result unchanged. With no motif at stake, the reference and alternative alleles of HEK293 were called at equal rates (1.05, 95 % CI 0.87 to 1.28), so allelic mapping bias does not explain the difference. The three datasets were basecalled by their depositors with different model versions, and the ML ≥ 250 operating point was not recalibrated between them.

The intronic test of Figure 5B scored heterozygous variants within 5 nt of an adenosine in the HG002 direct RNA reads when the adenosine lay inside a GENCODE v46 intron on the same strand and inside no exon of any transcript other than a retained-intron model. Sites first met the eligibility rules of the allele test on all reads. Only molecules with no splice junction overlapping the introns that contain the adenosine were counted, and a site was kept when at least 10 such molecules carried each allele and the less frequent allele made up at least 15 % of them. Sites were not selected on methylation. They were grouped by whether the variant created DRACH, broke it, kept it on both alleles or lay 3 to 5 nt away, and the share of molecules called m^6^A on each allele was pooled over sites with 95 % intervals from 2,000 site bootstraps.

Figure S11B gives the allele ratio by variant position and for the recall controls in the three direct RNA sets (HEK293, HG002 and GM12878) and in HEK293T GLORI (GSE210563),^21^ whose two wild-type libraries, two libraries treated with STM2457 at 20 µM for 24 h and two libraries of in vitro transcribed RNA were trimmed with cutadapt and mapped with GLORI-tools to the A-to-G converted genome and transcriptome. GLORI reads were assigned to an allele at heterozygous variants of the parental HEK293 genome described above that lay within 5 nt of the adenosine and carried no A on either allele in transcript orientation, and an adenosine read as A was counted as methylated and one read as G as unmethylated. A read was used when both positions had a base quality of at least 10 and it carried no more than three unconverted adenosines, and a site was eligible with at least 10 wild-type reads per allele and a minor-allele fraction of at least 15 %. Sites were selected on one wild-type replicate, where both alleles together read at least 10 % A with at least three A reads, and scored on the other, and the two folds were summed. Sites with a variant at −1 or +1 whose DRACH allele read at least 10 % A with at least three A reads in the pooled wild-type libraries were then scored in the pooled STM2457 and in vitro transcribed libraries (Figure S11C).

Figure S11E assigns the adenosine of each variant at −1 or +1 with GENCODE v46 to an exon, to an intron (defined as for Figure 5B), to a position that is intronic in one transcript and exonic in another, to the first base of an exon that follows an intron, or to no annotated feature, and pools motif-creating and motif-breaking variants without selection on methylation. Intronic sites were scored on all reads and, for the direct RNA data, on intron-retaining molecules alone. For intron retention by allele (Figure S11F), a read informed an annotated intron when it covered at least 10 nt beyond each end. It counted as spliced when it carried a junction within 5 nt of both ends, and as retained when at least 25 aligned nucleotides lay inside the intron with no junction overlapping it. At exonic sites of the allele test, reads were assigned to an allele by the site variant. At intronic sites, the site variant was phased with heterozygous variants that lie inside no annotated intron on the same strand, using the direct RNA reads that carry both (at least five reads of each allele and at least 90 % concordance), and every read spanning the site’s own intron was assigned to a haplotype by those variants alone. Strata with at least five informative reads per allele and both outcomes were combined as a Mantel–Haenszel odds ratio of retention, with 95 % intervals from 2,000 site bootstraps.

To scan beyond the motif, every heterozygous variant within 70 nt of a GLORI site called in wild-type HEK293T was scored in the same way when neither allele carried an A in transcript orientation. Each site was represented by its variant with the most wild-type reads, and the direction of the allelic difference at C/U variants was tested at each position with a two-sided sign test (Figure 5C; Figure S11G).

Variants with an A allele were read by genotype. At A/C and A/U variants, a GLORI read carrying G or A at the variant was assigned to the A allele, because GLORI converts an unmethylated A to G, and A/G variants were not used. The same comparison was made in HeLa cells, with genotypes called from the GLORI− libraries and the six control libraries pooled. In the HG002 and HEK293 direct RNA reads, every substitution at positions −20 to +20 was scored for an allelic difference, with Benjamini–Hochberg correction within each dataset, and a position was reported when it reached q < 0.05 in one dataset and P < 0.05 in the same direction in another, provided that false-positive calls at unmethylated adenosines showed no bias there (Figure 5D). To test whether methylation changes allelic abundance, we regressed the log ratio of alternative to reference reads, counting every read at the variant, on the difference in m^6^A per molecule between the two alleles at 11,959 exonic heterozygous variants in the HG002 direct RNA reads, excluding A/G variants and variants in the MHC region or on chromosome X. The 95 % interval comes from a bootstrap over variants, and a bootstrap over 10-Mb genomic blocks gave 0.94 to 1.04. The joint effect of an A at −4 and a U at +4 was tested on the levels of 213,750 GLORI sites in HEK293T, each relative to the mean of its 5-mer, against the product of the two single-base effects. Sites with both reached 1.191 times their 5-mer expectation, against 1.085 and 1.097 with either alone, an interaction of 1.0004 (95 % CI 0.995 to 1.006).

### Population census of motif-changing variants

For each exon +1, 5′SS position 4, branch-point and AT–AC 3′SS adenosine, every possible single-nucleotide substitution from positions −2 to +2 was labeled by whether it created or destroyed a DRACH at the adenosine or left its status unchanged, and observed variants were extracted with BCFtools 1.16 from gnomAD v4.1.1 genomes (76,215 individuals, FILTER PASS), which include the 76,156 genomes described by Chen et al.^74^ Substitutions that break the GT or AG dinucleotide were counted separately. Selection was tested as a Cochran–Mantel–Haenszel odds ratio of observed among possible substitutions, motif-creating against motif-neutral, within strata of position, substitution, trinucleotide context, CpG status and change in MaxEntScan score.^91^ The identical test on one random adenosine per MANE Select intron and internal exon, positions with no splicing role, served as the calibration. The census of 26,028 variants includes those at AT–AC 3′SS A residues, which Figure 5F does not show.

### Database union

We pooled reported m^6^A sites from m^6^A-Atlas v2.0,^77^ m^6^AConquer,^78^ DirectRMDB^79^ and RMBase v3.0.^80^ Records were lifted to hg38 with the UCSC hg19ToHg38 chain where necessary and deduplicated by position and strand. Coverage-aware comparisons were restricted to sites that the underlying assay could in principle observe.

### Branch points

Human branch-point annotations combined a lariat-sequencing set lifted to GRCh38^35^ with a released set,^36^ yielding 86,615 positions after deduplication, both resting on the excised lariat intermediate and the branch point mapped within it.^101^ Of 86,610 usable positions, 58,259 (67.3 %) had an A in the reference sequence and entered the context-composition analysis (Figure S14A), where we scored them for DRACH compatibility (Figure S14B). GLORI signals at the branch base were interpreted with caution because the 2′–5′ branched A may resist nitrite deamination, which would lower DRACH and non-DRACH branch points alike. Native-read rates, which do not depend on deamination, were calculated from PRJEB111137. U12 and U2 branch-consensus compatibility was also assessed across nine genomes (Figure 3G).

The U2 branch-point window of the Minor Intron Database is too short to contain the last position of DRACH, so that base was read from the genome assembly on which the database was built. Each species had to reproduce the database’s own U12 DRACH count before its U2 value was reported (Figure 3G).

### AT–AC coverage matching

We compared AT–AC 3′ splice sites with controls matched by depth in unconverted GLORI libraries, which omit the deamination treatment. For the 12 human GLORI libraries with these controls, we measured coverage at each candidate base with samtools depth -Q 255 and required GLORI− depth ≥ 10 for the primary comparison. Positions were deduplicated by chromosome, position and strand, overlaps between 3′SS and comparator sets were removed, and the 39 compatible human introns represented 38 distinct 3′SS A residues. Interior DRACH A residues from the AT–AC intron set provided the rate comparator, and non-DRACH AT–AC 3′ splice sites provided the motif control. Depth strata had lower bounds of 10, 15, 25, 40, 65, 105, 170, 275, 450, 730 and 1,200 reads. We calculated Mantel–Haenszel odds ratios and one-sided exact conditional tests separately for each library; pooled counts are descriptive site–library observations. For A549 libraries, detection opportunity was estimated from an intron signal-density surrogate excluding the focal site, and zeros there were interpreted against that limited chance of detection (Figure S4F). Because every mouse GLORI+ deposit lists called sites only (Table S3F), the mouse *Tmem231* 3′SS A was scored from the raw reads of the 12 realigned mouse GLORI libraries described above, counting A and G reads at the site as for the human s^4^U-GLORI libraries, and 11 of them covered it (GSE210563 MEF, four; GSE270643 dorsal hippocampus synaptic and cytoplasmic fractions, seven). Pooled over the 11 libraries the site had 52 covering reads, of which 5 were unconverted, 4 of them in one 7-read library, and no library reached the 15-read depth of a site call (Figure 3C). The pooled count was compared with the lower human level, 41 %, allowing a 2 % unconversion floor, by a one-sided exact binomial test, and the upper bound is the one-sided 95 % Clopper–Pearson limit corrected for the same floor. The only MEF GLORI− library of GSE210563 (GSM6432603, without heat shock), which omits the deamination step and cannot report methylation, covered the site with 4 reads. *PIGB* methylation rests on one replicate, the writer-dependence evidence comes from *DSN1*, and the non-human AT–AC comparisons are underpowered (Figure S5C).

### Controls

We excluded 3′SS A residues that coincided with an annotated transcription start site before calculating rates, because a union-of-transcripts intron model can label the first nucleotide of a major mRNA isoform as a 3′SS A when a minor isoform splices to that position. Of the 39 3′SS-A positions detected in at least four libraries, 17 overlapped annotated transcription start sites, a 150-fold excess over background, and those signals were independent of the mRNA m^6^A writer in every tested perturbation arm, as expected of cap-adjacent m^6^Am^10^ rather than internal m^6^A.

For the 5-mer-matched control, we compared canonical 3′SS A residues with A residues in the identical 5-mer elsewhere in the transcriptome at matched coverage, which estimates any positional difference in m^6^A-SAC-seq remaining after sequence context is matched. The odds ratio adds 0.5 to each cell (7 of 659 observable 3′SS A residues against 18 of 1,859 matched adenosines), and its 95 % CI is the Woolf interval; without the correction it is 1.10 (exact 95 % CI 0.39 to 2.77).

### RNA-seq and 3′ splice-site usage

Short-read RNA-seq came from 15 knockdowns in ERP133481, a HeLa siRNA panel covering 305 splicing factors and regulators,^56^ and from ENCODE experiments^75^ in which METTL3 and RNPC3 were targeted with CRISPR guide RNAs and RBM15, FTO and U2AF1 with shRNAs. Reads were aligned with STAR 2.7.10b^86^ using GENCODE v46 annotations and the options --outFilterMultimapNmax 20, --outFilterMismatchNoverReadLmax 0.06, --alignSJoverhangMin 8 and --alignSJDBoverhangMin 3, and BAMs were processed with samtools 1.16.1.^88^ At each AT–AC 3′SS pair, we counted spliced reads spanning the exon–exon junction and unspliced reads crossing the intron–exon boundary. Introns were grouped by DRACH compatibility rather than by detected methylation, because a 3′SS signal is measurable only on intron-retained molecules and grouping by methylation would select for retention. Perturbation effects were assessed against a control-versus-control null calculated at each locus. The exon +1 retention analysis used companion libraries from GSE210563 (Figure S8E).

For the nearest-neighbor exon +1 comparisons, we matched each detected 3′SS to 50 controls on standardized log read depth, pyrimidine content, log intron length and log GLORI gene coverage, and perturbation comparisons also matched on the baseline value of the outcome tested (Figures S8E and S8F). We report the median, across focal 3′ splice sites, of each focal value minus the median of its controls. The reported percentile 95 % intervals used 500,000 resamples of genes (2,000 for Figure S7D), each carrying the differences of its focal 3′ splice sites, with the comparator pool and matches held fixed. For the polypyrimidine-tract comparison, every canonical 3′SS whose exon +1 A lies in a boundary-spanning DRACH (9,007 distinct sites) was scored for uridine and pyrimidine content, longest pyrimidine and uridine runs and purine interruptions over intron positions −40 to −3, from GRCh38. Sites with an exon +1 call in any HeLa or HEK293T control-state GLORI library (137 with unspliced depth ≥ 10) were each matched to their 50 nearest uncalled sites on intron retention and log read depth in the HeLa control mRNA-seq, the median paired difference was bootstrapped over sites (2,000 resamples) and tested with a two-sided Wilcoxon signed-rank test, and a logistic model of call status on each metric adjusted for retention, depth and intron length; P values were Benjamini–Hochberg-adjusted across the eight metrics and two tests.

### Comparative test in yeast

The fungal writer consensus, RGAC, requires G before the methylated A and so predicts a different outcome at branch points from DRACH. In *S. cerevisiae* and *S. pombe*, the branch A of the introns that carry the exact UACUAAC branch heptamer (222 and 644 introns) lies in UAAC, which is DRACH-compatible in 90.5 % and 86.8 % of them but RGAC-compatible in none, so the yeast writer is predicted to methylate neither the branch A nor the canonical 3′SS A. The first endogenous AT–AC intron reported in budding yeast, an unconventional intron nested inside the annotated *RPL30* intron, is spliced from an AT 5′ splice site to either of two AC 3′ splice sites.^102^ Both end in UAC, so a pyrimidine precedes the A and neither matches DRACH or RGAC. In *S. cerevisiae*, m^6^A is deposited during meiosis,^15^ and *ime4*Δ cells provide a writer-null control.

The yeast analysis used PRJEB44511 (three sporulating wild-type and three ime4Δ replicates);^53^ PRJNA1365163^71^ served only as a coverage proxy. Signal-level event alignment used f5c 1.4,^90^ and differential modification was tested with Nanocompore 1.0.4^53^ on minimap2 2.28 alignments.^89^ Validation criteria were specified before inspecting the results. Splice-site coverage was insufficient, no branch or 3′SS A in *S. cerevisiae* is compatible with the writer consensus, and the run detected none of its interior positive controls, so the local null results are uninformative rather than negative.

Synthetic RNA004 controls show that a motif-agnostic model can detect m^6^A in both splice-site contexts (Figure S2), but they cannot validate this RNA002 comparison or set its error rate on native RNA. Nanocompore’s GMM_logit_pvalue column is already Benjamini–Hochberg-adjusted, so the significance threshold was applied to it directly, and the prespecified breadth criterion required significant k-mers in at least 1,000 transcripts. The comparison recovered the expected sequence preference, with 87.7 % of the 896 significant RGAC positions favoring the wild type against 23.4 % of the 7,703 other positions, and 15,159 significant k-mers across 1,712 transcripts passed the breadth criterion, compared with 15,961 across 1,510 reported by Leger et al.^53^ PRJNA1365163 holds SEN1-background direct RNA libraries with poly(A) tails added in vitro, which we used only to project the depth and coverage attainable in sporulation-stage RNA, not to infer m^6^A. The wild-type and *ime4*Δ libraries had 0.43 and 0.38 times the read depth of this proxy, and in one replicate pair only 19 of 285 intron-containing pre-mRNA models had at least 10 reads in both genotypes, because in vitro polyadenylation captures 3′-truncated unspliced species that native poly(A) selection misses. Coverage of unspliced pre-mRNA was therefore about an order of magnitude below the projection, and the prespecified window test found no significant position among 95 3′SS A residues, 63 branch A residues or 58 interior RGAC positive controls.

### Structural modeling

We used *S. pombe* U2AF1–RNA structures PDB 7C06, 7C07 and 7C08,^7^ postcatalytic spliceosome structures 6EXN,^47^ 6BK8^81^ and 6QDV (Figure S15),^82^ and structures after branching, 5GMK^46^ and 5LJ3.^83^ Adenine and its N6 methyl group were modeled into each deposited copy, and we scanned glycosidic and methyl rotations with the base held rigid, recording steric overlap and hydrogen-bond geometry (Figures S15A–S15G). The analysis assesses geometric feasibility in deposited coordinates, not reaction rates or binding affinities.

In Figure S15D, the glycosidic torsions of both bases were each sampled over 0–360° in 2° steps from the deposited anti conformation, giving 32,400 joint states. The separation in Figure S15E follows from the law of cosines with one bond length and is exact at the measured approach angle, and the scan of Figure S15D tests whether larger rotations avoid the clash. In the human postcatalytic complex 6QDV, the unmodified pair does not reproduce the 6EXN distances (2.37 and 4.91 Å). At exon +1 of the U2AF1 structures (Figure S12), an adenine was modeled by placing N6 at the position of the guanine O6, and the N6 methyl was built in each of its two in-plane orientations with rigid geometry.

## Quantification and statistical analysis

Statistical details are given in the figure legends and in the Results where each value is reported, and the tests, thresholds and resampling units are defined in Method details and below. Where each value is reported, its n and what n counts are given in the legend, the Results or Method details, and n is distinct genomic positions, site–library observations, reads at a position, introns, genes or heterozygous variants, as stated for that analysis. Pooled rates are fractions of reads or of site–library observations, ratios compare such rates within one library or dataset, matched comparisons report the median difference from matched controls, and Figure 2B shows medians with interquartile ranges. Unless stated otherwise, intervals are 95 % confidence intervals, from the Wilson score method for proportions and from percentile bootstraps over the named resampling unit for ratios and medians. Tests were Fisher’s exact test, exact binomial tests, two-sided sign tests, two-sided Wilcoxon signed-rank tests on matched differences (Figure S8A), Mantel–Haenszel odds ratios with exact conditional tests and logistic regression, each named where its P value is given, and per-site calls in the realigned GLORI libraries used one-sided binomial tests against the library background with Benjamini–Hochberg correction. Significance was taken at P < 0.05, or at q < 0.05 for per-site calls, and the multiplicity of the main-text tests is assessed below. No statistical method was used to predetermine sample sizes, which were set by the public datasets, and randomization and blinding did not apply because no new experiments were performed. Coverage thresholds and exclusions are stated with each analysis in Method details.

Intervals on the matched intron-retention differences come from a bootstrap over genes, which resamples genes with replacement and pools their 3′ splice sites. The expectation in Figure S10C scores each site at the methylated share of the intronic reference sites (46,088 DRACH adenosines with at least 20 reads, 50 to 500 nt upstream of a 3′SS, in introns with at least 10 reads at the 3′SS) that share its 5-mer and its intron’s splicing-efficiency quintile; each 5′SS A4 is assigned to the intron whose 5′SS it occupies, and its 95 % interval comes from 2,000 replicates that redraw every reference cell binomially, the class composition multinomially and the observed count binomially. Figures 4C and 4D show 5 % samples of the intronic and exonic DRACH references (2,310 and 3,773 sites with at least 20 reads). Per-class logistic regressions of methylation on the D, R and H bases, the RNAplfold probability that the adenosine is unpaired,^92^ read depth and exon architecture are in Figure S9C. Without the splicing-efficiency matching, the branch-point ratio was the same under three treatments of read depth (Figure S9D). The association of methylation with ENCODE eCLIP peaks within 25 nt^75^ was adjusted for read depth and peak count (Figure S9E). Matching also on whether an exon of another same-strand GENCODE v46 transcript that is not a retained-intron annotation overlaps the intron (Figure S16) left exon +1 at 58 % of the expected level (95 % CI 39 to 81 %; Supplemental Note 2). Keeping only reads that prove a precursor or lie at positions that no other transcript’s exon covers, on both the signals and the reference, gave 79 % of the expected level for exon +1 (95 % CI 52 to 113 %) and 71 % for branch points (31 to 116 %). In the s^4^U-GLORI libraries, matching on the splicing efficiency of the same introns in the DRS reads moved the steady-state poly(A)+ branch-point ratio from 0.56 to 0.65 (95 % CI 0.43 to 0.96; 0.64 in the second replicate) and the exon +1 ratio from 0.83 to 0.74 (95 % CI 0.57 to 1.00; 0.73 in the second replicate). In the same libraries, exon +1 was also compared with the DRACH adenosines 2 to 150 nt upstream and 50 to 150 nt downstream of its 3′SS on reads that run unspliced through the 3′SS, after subtracting each library’s unconversion background and standardizing for the 5-mer, with a 95 % interval from 100,000 bootstrap replicates over introns. The ratio of fold decreases in Figure S8D uses 2,000 joint draws over gene components, and the perturbation medians in Figure S8F use 500,000 draws over the same components. Figure 4B plots elevated sites per 1,000 covered site–library observations, whereas the Fisher’s exact test quoted beside it is computed on unique positions, and both counts are reported.

Across the 11 P values reported in the main text and figure legends, namely the cross-species *TMEM231* contrast, the polypyrimidine-tract comparison, the three per-site motif contrasts in newly synthesized RNA and the six allelic sign tests, Holm and Benjamini–Hochberg adjustments, applied to the values as reported, leave every comparison below 0.05, the least extreme being 0.008 after Holm adjustment. The AT–AC class contrast is reported as counts, because it rests on 3 of 28 covered sites in each HEK293T replicate, and where no other transcript’s exon covers the 3′SS A, 1 of 10 compatible residues (*TSHZ3*) and none of 45 non-DRACH ones were methylated. These adjustments do not account for every test performed in the underlying analyses. As a sensitivity analysis, we removed duplicate reads, by unique molecular identifier in the HEK293T and HeLa GLORI libraries and by fragment start, end and strand in the s^4^U-GLORI and bladder libraries. Exon +2 adenosines then rose above background in five rather than six s^4^U-GLORI libraries, the per-site DRACH to non-DRACH ratios of Figure 4B rose 1.7-to 2.2-fold and the canonical 3′SS A fell from 0.18 to 0.08 elevated observations per 1,000, the bladder 3′SS A stayed within 6 % of its YAG background, and the HEK293T and HeLa allelic ratios rose by 5 to 9 %, with no conclusion changed.

Figures and Supplemental tables were generated with Python 3.11.15, numpy 2.4.3, scipy 1.17.1 and matplotlib 3.10.8, the environment of the code deposit. The upstream analyses ran in several environments, and the code deposit lists the software versions of every step. The corrected native-register readout used Python 3.9.23, NumPy 2.0.2 and SciPy 1.13.1. K.R.R. wrote the analysis, plotting and schematic code with Claude Code (Anthropic) using Claude Opus 5, Opus 5.5, Fable 5 and Fable 5.1, and the code deposit contains all of it.

