## Supplemental Figures for "The m^6^A motif is an anti-3′ splice site"

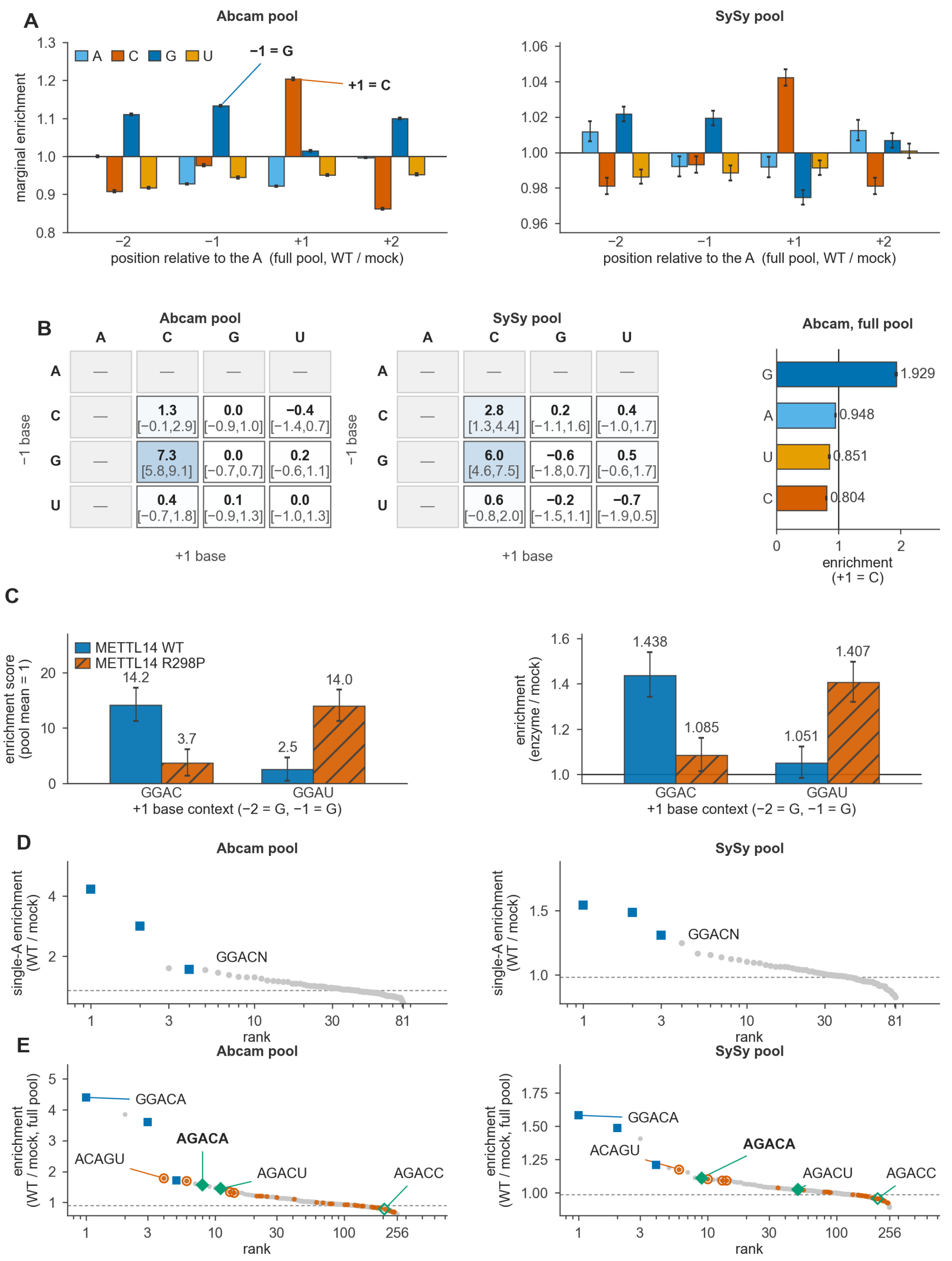

**Figure S1. Sequence specificity of METTL3–METTL14 in vitro, related to Figure 1.** IVM-seq libraries of Zhang et al. [S1], selected with the Abcam or the Synaptic Systems (SySy) antibody. Scores are enrichment over mock, not methylated fractions.

**(A)** Enrichment of each base at positions −2 to +2 in the full pool.

**(B)** Background-subtracted enrichment of single-A substrates by their −1 and +1 bases, with 95% intervals; dashes mark contexts with a second A. A pyrimidine at −1 in place of G removes 92% (Abcam) and 97% (SySy) of the GGAC score. The right plot shows the Abcam full pool at +1 C.

**(C)** The METTL14 R298P mutant reverses the +1 preference in the SySy libraries: GGAU rises from 2.5 to 14.0 while GGAC falls.

**(D)** Without motif information, GGACN ranks 1, 2 and 4 (Abcam) and 1 to 3 (SySy) among the 81 single-A 5-mers.

**(E)** All 256 A-centered 5-mers ranked by full-pool enrichment, as in Figure 1F. AGACA ranks 8 (Abcam) and 9 (SySy), and the 32 5-mers of the 3′SS register sit near the pool median. Open circles mark the ACAGN 5-mers, which share reads with GGACA.

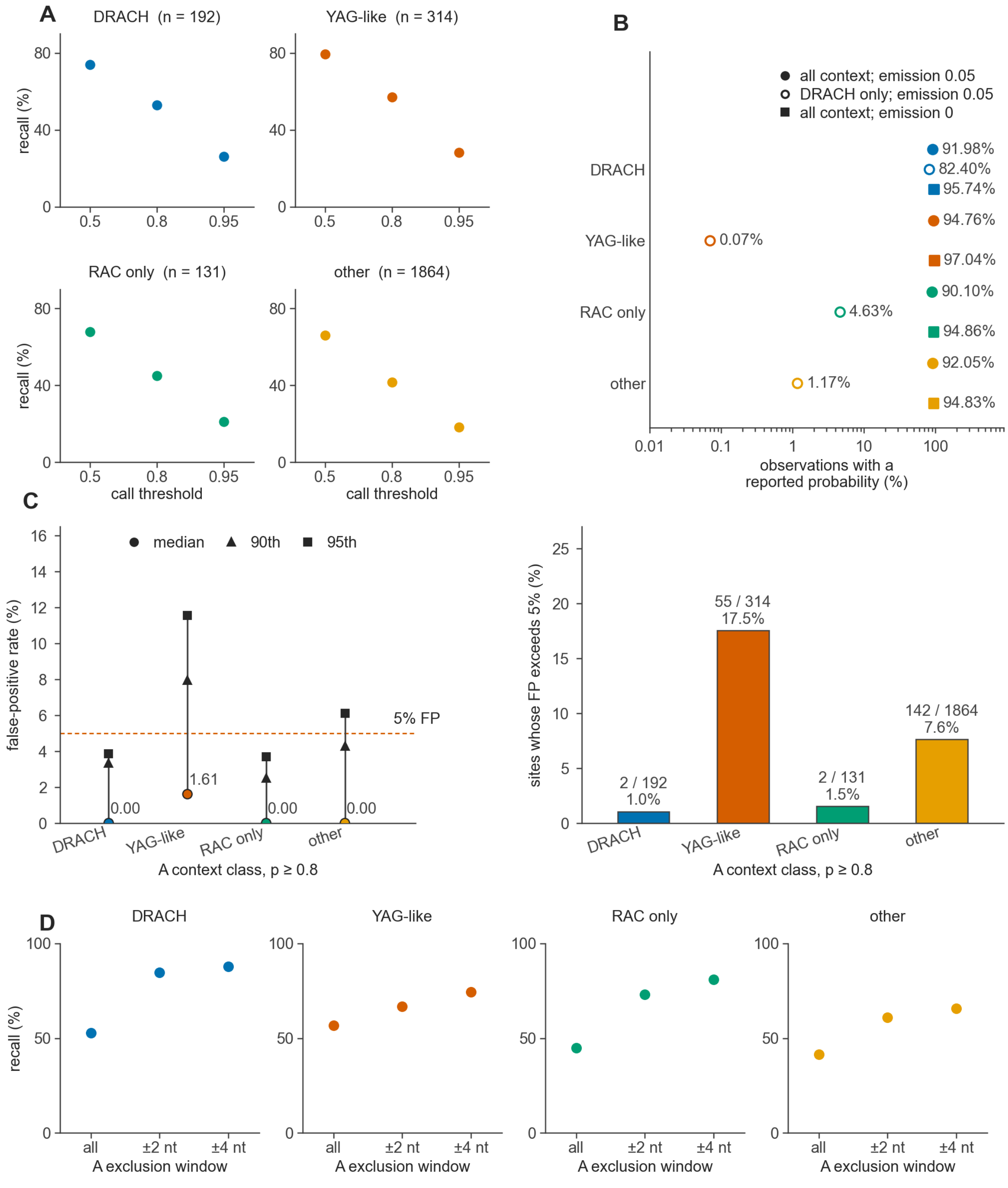

| | DRACH all | DRACH ±2 nt | DRACH ±4 nt | YAG-like all | YAG-like ±2 nt | YAG-like ±4 nt | RAC only all | RAC only ±2 nt | RAC only ±4 nt | other all | other ±2 nt | other ±4 nt |
|---|---|---|---|---|---|---|---|---|---|---|---|---|
| n | 192 | 44 | 16 | 314 | 184 | 49 | 131 | 53 | 25 | 1864 | 509 | 176 |
| FP (%) | 0.0 | 0.0 | 1.6 | 1.6 | 1.5 | 1.6 | 0.0 | 0.0 | 0.0 | 0.0 | 0.0 | 0.0 |

**Figure S2. Detection of m⁶A by the all-context nanopore model in splice-site contexts, related to Figure 2.** Synthetic curlcake RNA sequenced with RNA004 chemistry (PRJEB82528), 400 reads per sample, with a call threshold of 0.8 unless stated.

**(A)** Median site recall by context class and call threshold. At 0.8, recall is 52.8% at DRACH sites (192), 57.0% at 3′SS-like YAG sites (314) and 44.9% at branch-point-like RAC-only sites (131).

**(B)** Read observations with a reported m⁶A probability. At the default emission threshold, the DRACH-restricted model reports 0.07% of YAG-like observations against 94.76% for the all-context model, so only the all-context model can score a 3′SS A.

**(C)** False-positive rates on unmodified RNA (left, median and 90th and 95th percentiles) and the share of sites above 5% (right). At YAG-like sites the median is 1.6%, but 55 of 314 sites exceed 5%, so comparisons between conditions are more reliable than single site calls.

**(D)** Recall rises when no other A lies within ±2 nt, as at a yeast 3′SS in a polypyrimidine tract, reaching 66.7% at YAG-like and 73.1% at RAC-only sites. These controls show that the model detects m⁶A in splice-site contexts but do not bound its false-positive rate on native RNA.

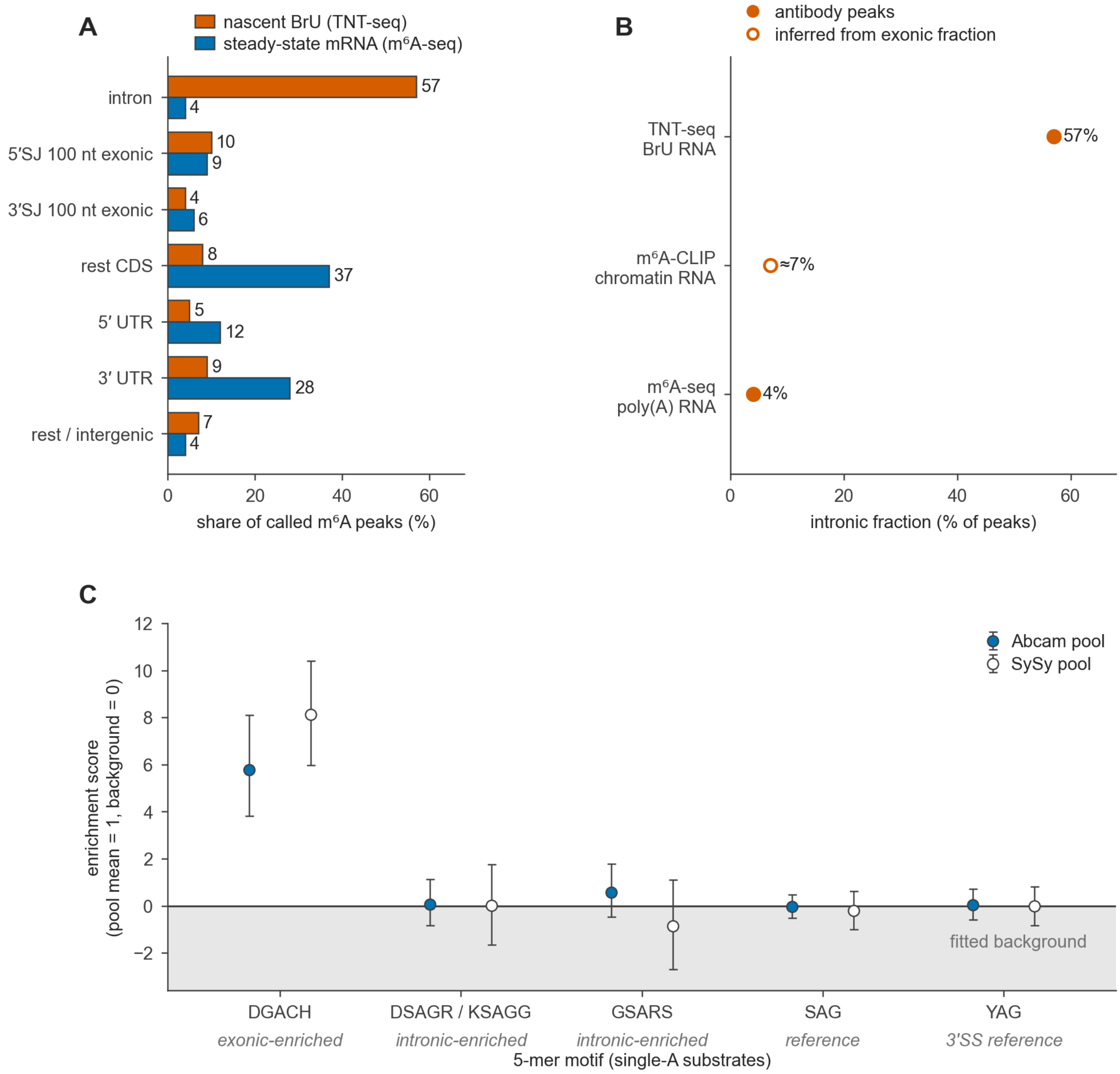

**Figure S3. Intronic m⁶A in nascent RNA, and in vitro methylation of the motifs enriched at intronic peaks, related to Figure 2.**

**(A)** m⁶A peaks by region in nascent BrU-labeled RNA (TNT-seq, 15-min pulse, HEK293T) and in steady-state mRNA [S2]. Introns hold 57% of nascent peaks and 4% of steady-state peaks, although peaks of 20 to 100 nt cannot resolve the 3′SS A.

**(B)** The intronic share of m⁶A antibody peaks across RNA pools. It falls from nascent to steady-state RNA (57% to 4%), although Ke et al. [S3] found a low intronic share in chromatin-associated RNA (open circle, inferred from their approximately 93% exonic m⁶A).

**(C)** In vitro enrichment (IVM-seq) of the four de novo motifs of Louloupi et al. [S2], each centered on its A, with 95% intervals (filled, Abcam pool; open, SySy pool). Louloupi et al. hypothesized that early intronic m⁶A lies mostly outside DRACH, but only DGACH, which matches DRACH, rises clearly above background (5.8 with Abcam and 8.1 with SySy); the SAG-core motifs enriched at intronic peaks do not.

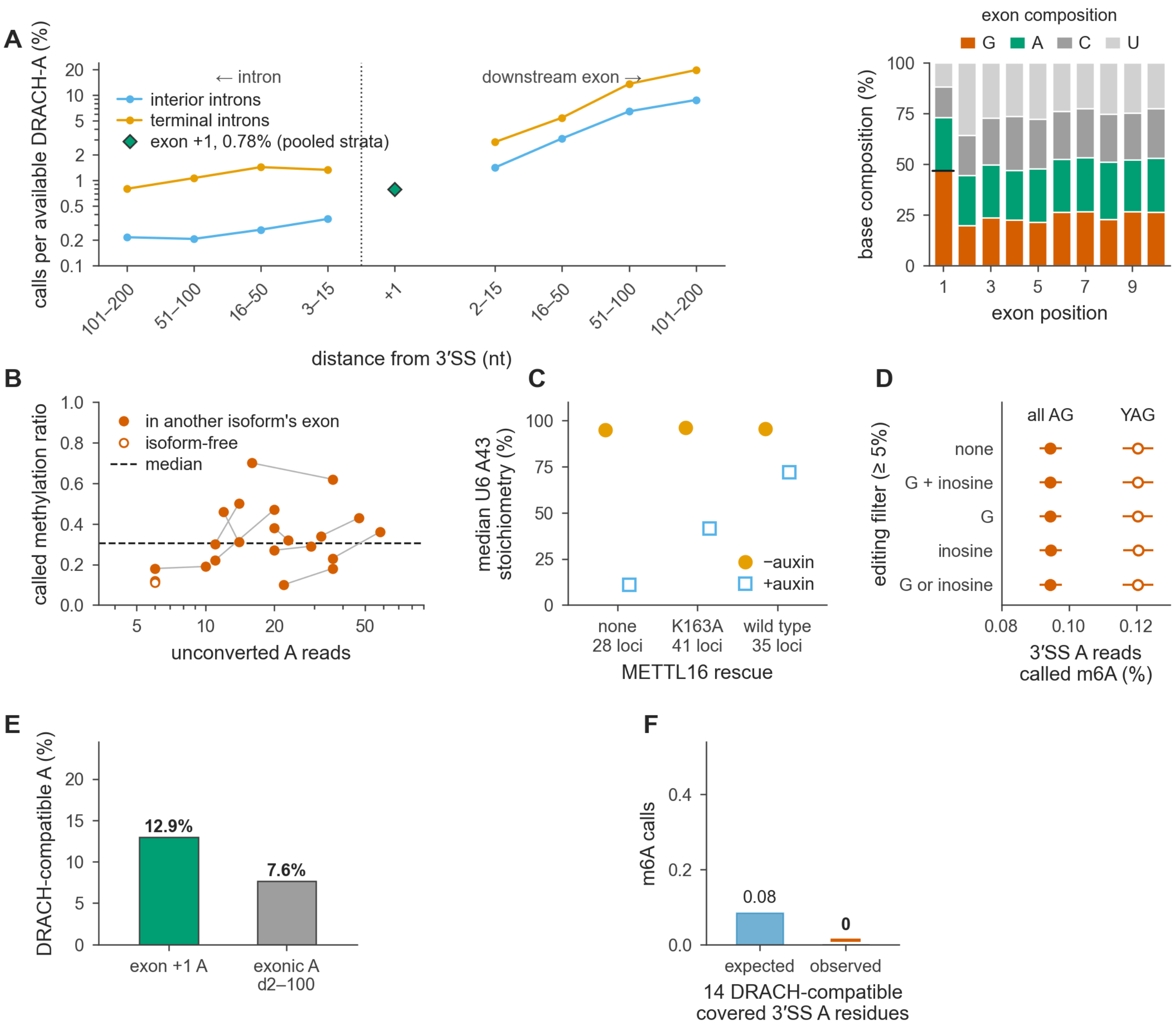

**Figure S4. Methylation near 3′ splice sites in chromatin-associated RNA, and controls for detection and editing, related to Figure 2.** Panels (A), (B), (E) and (F) use chromatin GLORI of A549 cells [S4], on $m^6A$ sites called in both replicates.

**(A)** Detected sites per available DRACH A, counting positions inside any annotated intron of a protein-coding gene on the intron side; at right, the share of G at the first exonic position (47%). Interior exons fall from 8.8% at 101–200 nt to 1.4% at 2–15 nt, the depletion that Sethi et al. [S5] found on unspliced molecules. Exon +1 (diamond) reaches 0.78% (11 of 1,408), similar to the 0.73% of exon positions +2 to +15 pooled over all intron classes (71 of 9,678), a pooled value the class-separated bins do not show.

**(B)** The 11 3′SS A signals called in both replicates are all non-DRACH (10 CAG, 1 UAG); ten lie inside an exon of another transcript, eight of them where another transcript uses an upstream acceptor, and their levels range from 0.10 to 0.70. Lines join the two replicates of each site.

**(C)** U6 A43 methylation after auxin-induced METTL16 degradation in HCT116 cells [S6], with no rescue, the hyperactive K163A mutant or wild-type METTL16 (28, 41 and 35 loci). Without rescue the median falls from 95% to 11%.

**(D)** In direct RNA of HEK293 cells [S7], $m^6A$ calls at the 3′SS A occur in 0.095% of reads at all AG acceptors (filled) and 0.120% at YAG acceptors (open). Removing every acceptor edited in at least 5% of reads, with editing scored as basecalled G, as Dorado's inosine calls, as their sum or as either one, changes neither rate by more than 0.0001 percentage points and removes at most 655 of 36,230 acceptors. Lines are 95% intervals.

**(E)** The intron's terminal AG makes 12.9% of exon +1 A residues DRACH-compatible, against 7.6% at exon positions +2 to +100.

**(F)** None of the 14 covered DRACH-compatible AT–AC 3′SS A residues is detected (0.08 expected from the intronic DRACH rate).

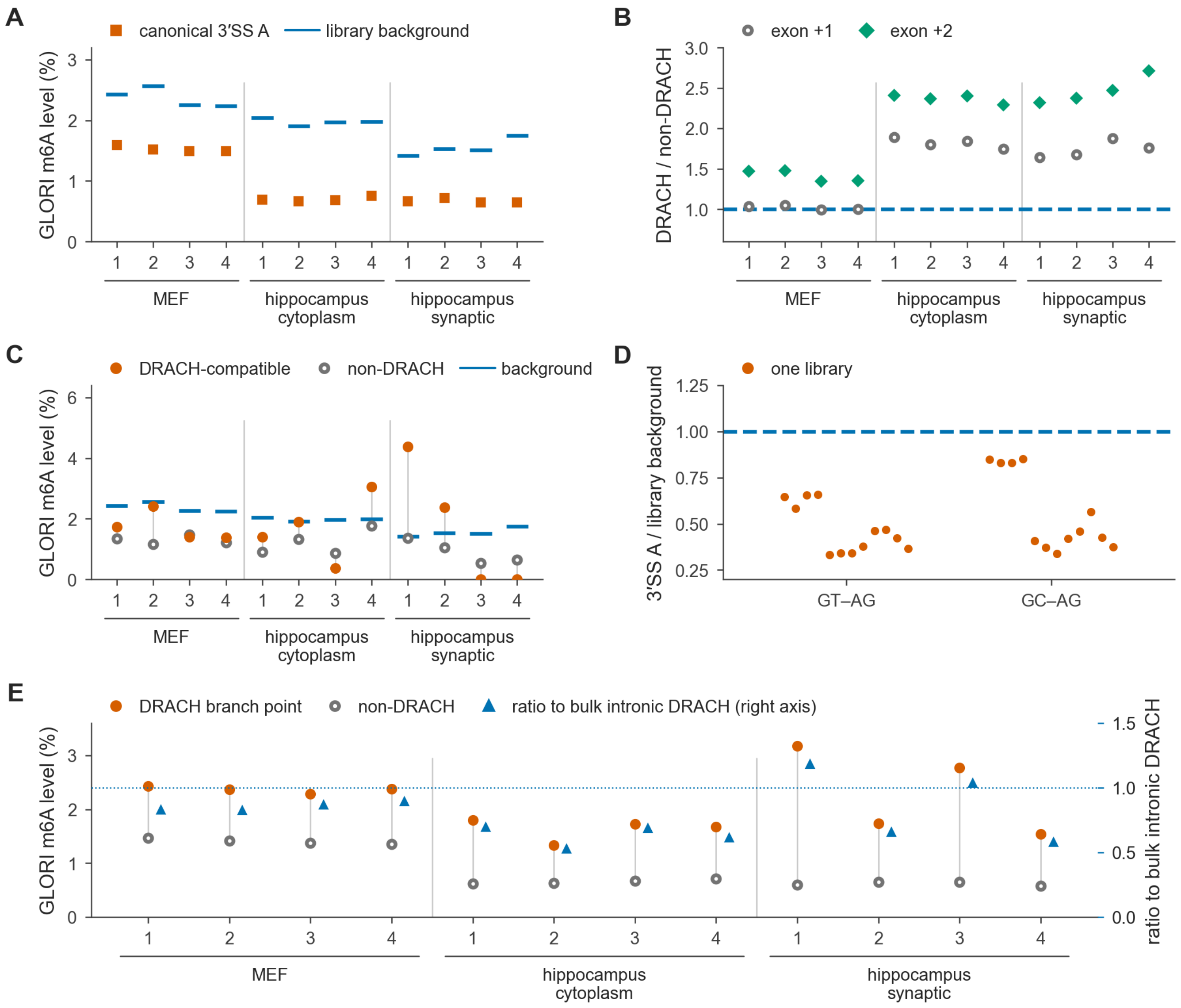

**Figure S5. The mouse GLORI census, library by library, related to Figure 2.** One column per mouse GLORI+ library, four fibroblast (GSE210563, leftmost; 1 and 2 heat-shocked) and eight dorsal hippocampus (GSE270643; cytoplasmic and synaptic).

**(A)** The canonical 3′SS A sits below background in 12 of 12, over 700,312 site–library observations.

**(B)** At exon +2, DRACH exceeds non-DRACH in 12 of 12 (median 2.34-fold), while exon +1 runs 0.99 to 1.89 and splits by tissue.

**(C)** AT–AC 3′SS A of protein-coding genes (16 DRACH-compatible and 163 non-DRACH 3′SS A residues, from the 18 and 191 introns of Figure S6C). Compatible sites exceed background in only 3 of 12 (1.73% against 1.22% pooled; 1.16% against 1.13% outside other transcripts' exons), so methylation is not established in mouse; non-DRACH sites fall below it in 12 of 12.

**(D)** In the ±500 nt scan, the 3′SS A of GT–AG and GC–AG introns sits below background in every library.

**(E)** Taggart et al. [S8] branch points, 386 DRACH-compatible ones covered. DRACH branch points exceed non-DRACH ones in 12 of 12, yet sit at or below the bulk intronic DRACH rate (triangles, right axis) in 10 of 12 (9 of 12 outside other transcripts' exons). Rates are unconverted-A fractions, not stoichiometries.

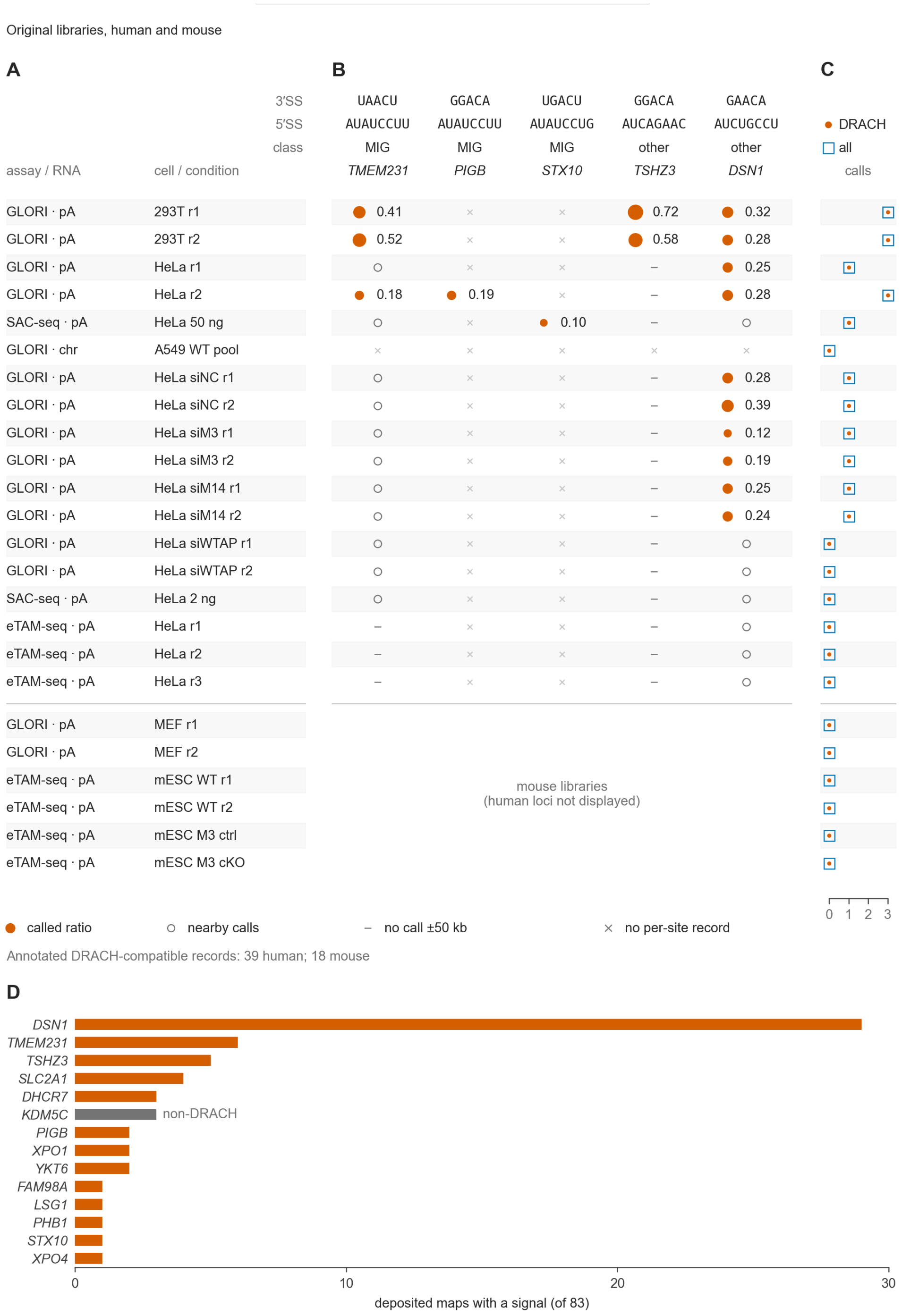

**Figure S6. AT–AC 3′ splice-site signals in the deposited m⁶A maps, related to Figure 3.** The survey covers 83 site sets (77 human and 6 mouse; Table S3D). (A)–(C) show the original libraries, and (D) summarizes the whole survey per gene.

**(A)** Assay, RNA fraction, cell type and condition of each site set.

**(B)** Signals at the 3′SS A of each gene, with the called ratio where GLORI or m$^6$A-SAC-seq reports one (circle size and value). Open circles mark signals near but not at the 3′SS, dashes no signal within ±50 kb and crosses no per-site record; none establishes coverage of the 3′SS A. MIG, minor-intron gene in the Minor Intron Database; other, class unverified.

**(C)** Signals at all AT–AC 3′ splice sites (open squares) and at the DRACH-compatible ones (dots). Protein-coding genes hold 271 AT–AC introns in GENCODE v46, 39 with a DRACH-compatible 3′SS A, and 209 in vM25, 18 of them compatible.

**(D)** m$^6$A recurrence across the deposited record, summarized per gene, with one bar per gene counting the site sets that report a signal at its AT–AC 3′SS A. Of the 83 site sets, 40 report an m$^6$A signal, giving 61 records at 14 genes, 58 at DRACH-compatible sites (vermilion), 29 of them at *DSN1*, and 3 at *KDM5C* (gray), each listed in Table S3E. An absent record means no signal, not a measured zero, and antibody-based assays place a site at the nearest consensus motif, so their records support rather than confirm methylation at a DRACH-compatible 3′SS.

pA, poly(A); chr, chromatin; r, replicate; M3/M14, METTL3/METTL14; cKO, conditional knockout.

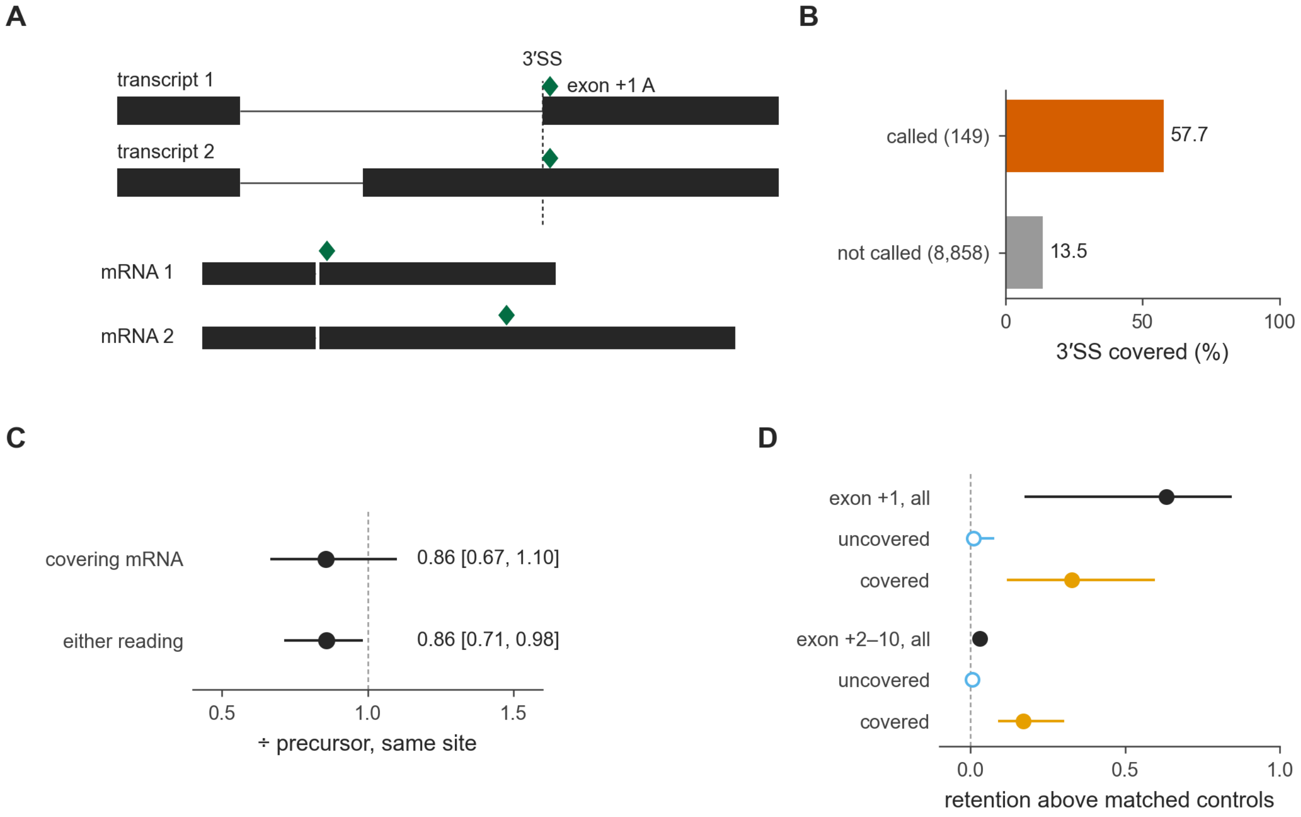

**Figure S7. Exon +1 methylation at 3′ splice sites that another transcript's exon covers, related to Figure 4.** A 3′SS is covered when one exon of another same-strand GENCODE v46 transcript, retained-intron annotations excepted, contains both its last intron base and exon +1.

**(A)** In transcript 2, which covers the 3′SS of transcript 1, the exon +1 A (green) is an internal exonic A that follows the intron's AG, so its mature RNA (mRNA 2) runs across the 3′SS without a gap.

**(B)** Share of DRACH-compatible exon +1 acceptors with a covered 3′SS, with and without an exon +1 GLORI signal in HeLa or HEK293T poly(A)+ RNA.

**(C)** HEK293 direct RNA at covered exon +1 sites: modified-read fraction of reads that fit the covering transcript's mRNA, or either reading, over reads that prove a precursor at the same site, with 95% intervals over introns.

**(D)** Intron retention of 3′SS with an exon +1 or exon +2 to +10 GLORI signal, minus 50 matched controls (median, 95% interval), by coverage.

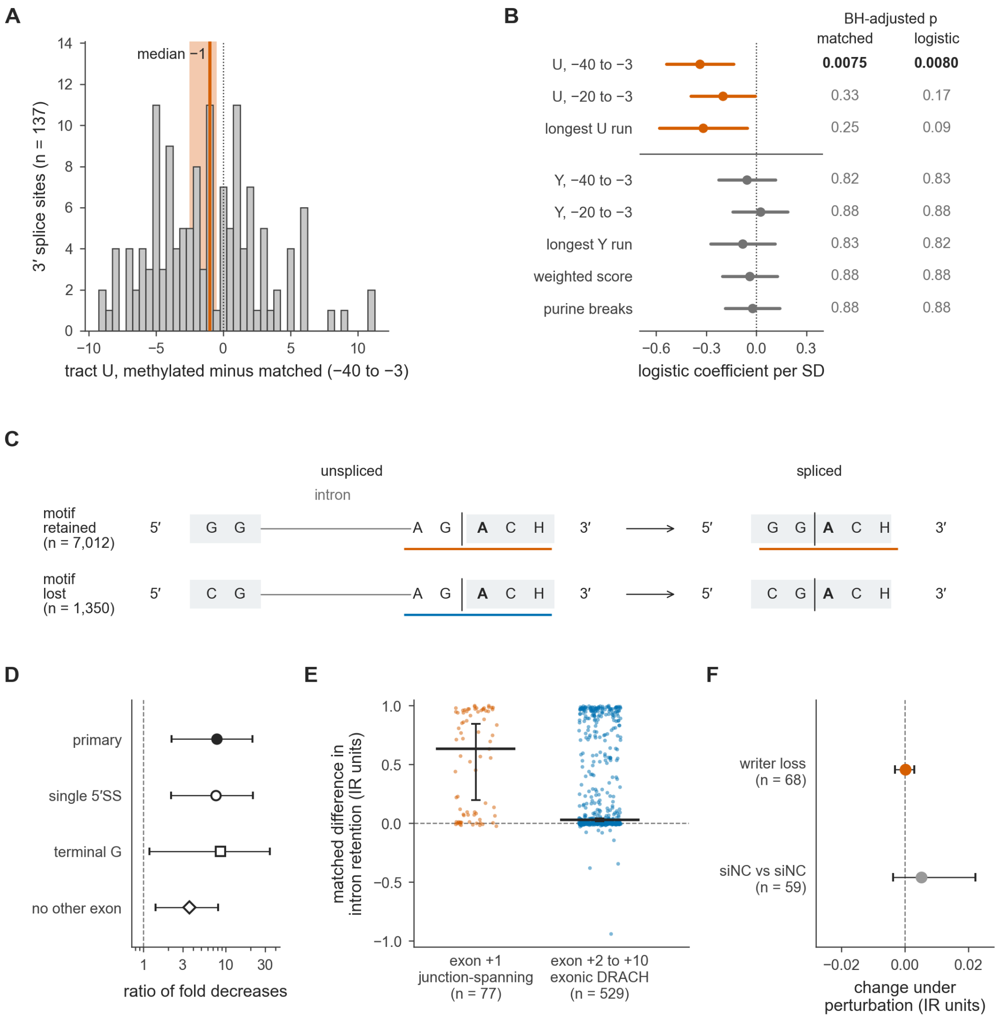

**Figure S8. Exon +1 methylation, the polypyrimidine tract, motif fate and intron retention, related to Figure 4.**

**(A and B)** Canonical 3′ splice sites whose exon +1 A lies in a boundary-spanning DRACH (9,007 sites), comparing the 137 with an exon +1 m$^6$A call in HeLa or HEK293T GLORI against the 4,616 without one whose gene is covered in HeLa GLORI.

**(A)** Uridine count over intron positions −40 to −3, each methylated site minus the median of its 50 nearest sites on intron retention and depth. The median difference is −1 (95% bootstrap interval −2.5 to −0.5; Wilcoxon signed-rank test, Benjamini–Hochberg-adjusted P = 0.0075).

**(B)** Logistic coefficient per standard deviation for each tract metric, adjusted for retention, depth and intron length, with 95% intervals and BH-adjusted p values from both tests (Y, pyrimidine). Only the uridine count over −40 to −3 survives correction in both, at −0.34 per SD (BH p = 0.0080); pyrimidine counts that include C show no difference at matched retention, so matching on pyrimidine count cannot detect the effect. Whether weaker uridine tracts give the writer access to exon +1, or slow splicing and lengthen nuclear residence, is not resolved by these data.

**(C)** Splicing replaces the intronic AG with the upstream exon’s last two bases, so GG recreates the boundary-spanning DRACH (motif retained) and CG destroys it (motif lost), which retains the motif at 7,012 of the 8,362 3′ splice sites with a spanning DRACH (84%).

**(D)** In HEK293 direct RNA reads, the modified-read fraction falls from unspliced to spliced reads 7.7-fold more where the motif is lost than where it is retained (95% interval 2.2–20.9; filled circle); the fractions are in Figure S9A. The open symbols repeat the estimate on 3′SS with a single 5′SS (7.5-fold) and, among those, on upstream exons ending in G in both classes (8.6-fold), which gives the two classes the same 5′SS −1 base. The open diamond repeats it on 3′SS that no other transcript’s exon covers (3.6-fold, 95% interval 1.4–8.1).

**(E)** Intron retention (IR) at 3′SS with an exon +1 GLORI signal, in 18 unconverted GLORI− HEK293T and HeLa libraries of GSE210563 [S9], minus 50 matched controls. Signals in a boundary-spanning DRACH at exon +1 (n = 77) sit +0.63 IR units above their controls (95% interval 0.2–0.8), against +0.03 at exon +2 to +10 (n = 529).

**(F)** Under writer loss (siMETTL3 + siWTAP, n = 68) the same sites change by +0.0002, within the siNC-versus-siNC variation (+0.0052, n = 59), so the association in (E) does not show that the mark causes retention.

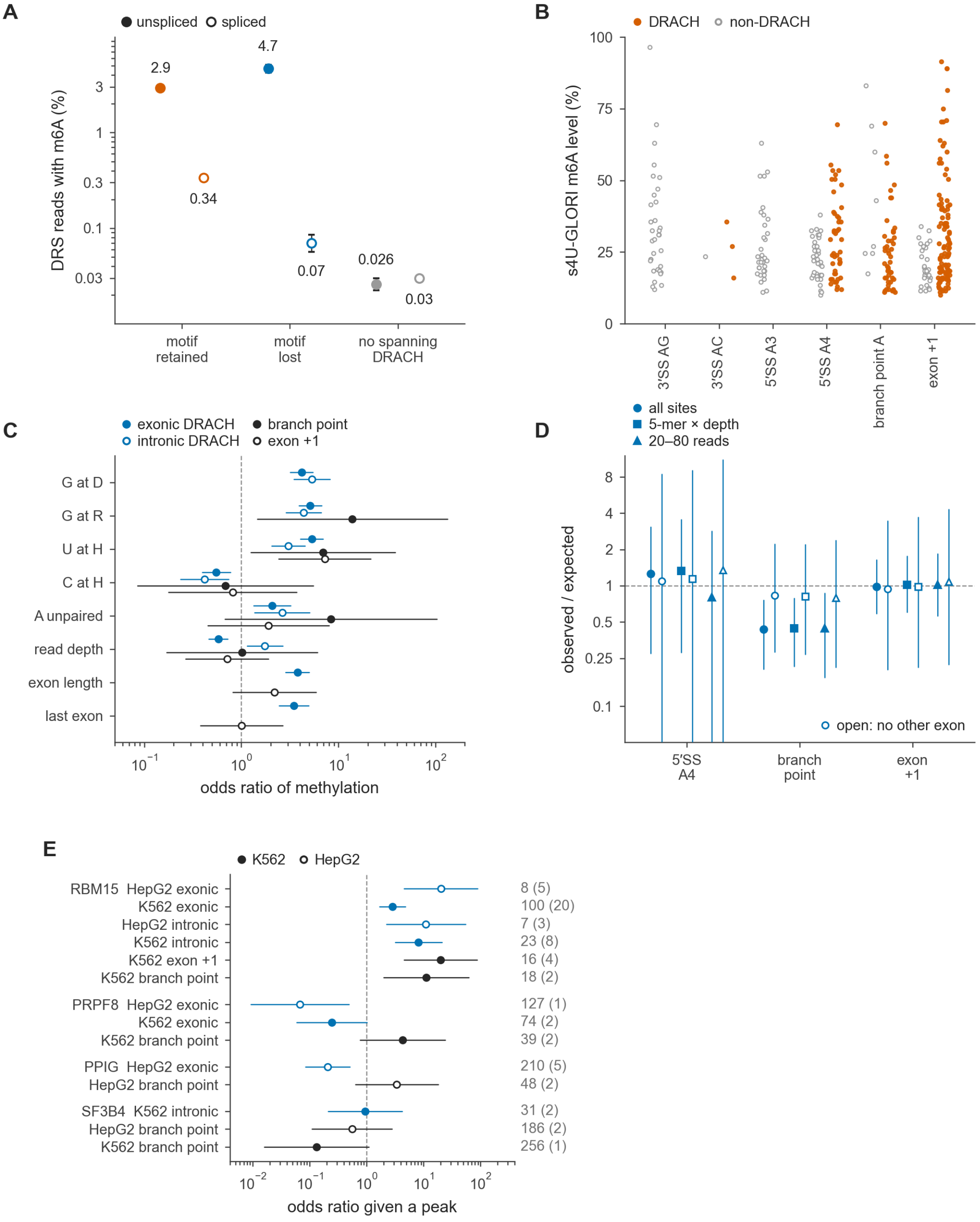

**Figure S9. Determinants of DRACH methylation at the splicing signals, related to Figure 4.** HEK293 direct RNA reads unless stated, at sites with at least 20 reads; a site is methylated when at least 20% of its reads carry modified calls.

**(A)** Exon +1 by motif fate and splice state (Wilson 95% intervals; ratio in Figure S8D).

**(B)** Methylation levels of sites elevated in both replicates of an $s^4$U-GLORI fraction [S10], DRACH filled, non-DRACH open.

**(C)** Per-class logistic regression on the D, R and H bases, the RNAplfold probability that the A is unpaired, read depth and exon architecture (odds ratios, 95% intervals).

**(D)** Each class over its 5-mer-matched expectation under three read-depth treatments, without Figure S10C's splicing-efficiency matching. The branch-point ratio is 0.44, 0.44 and 0.44, so read depth does not explain it. With the branch points and the reference both restricted to positions that no other transcript uses as exon (open symbols), it is 0.83, 0.81 and 0.80, with intervals that include 1.

**(E)** Odds of methylation given an ENCODE eCLIP peak within 25 nt [S11], adjusted for depth and peak count; labels give sites with a peak (methylated in parentheses). eCLIP comes from HepG2 and K562 cells, and peaks cover a minority of introns, so a site without a peak is not necessarily unbound.

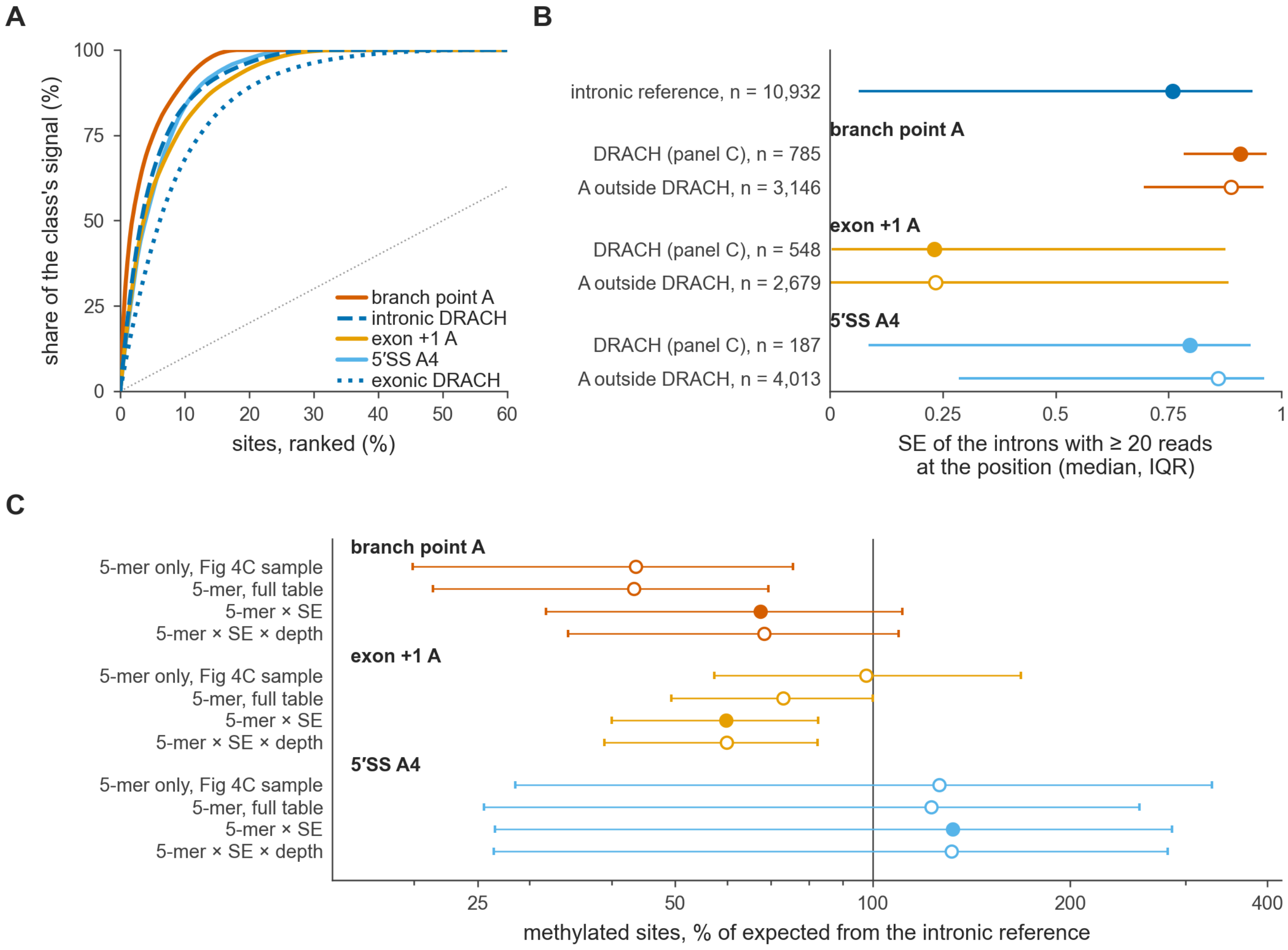

**Figure S10. Splicing efficiency, signal concentration and the splice-signal methylation comparisons, related to Figure 4.** HEK293 direct RNA; SE, the spliced share of reads at each 3′SS.

**(A)** The per-site modified-read fractions of Figure 4C as a Lorenz curve: half of each class's signal sits in 2 to 6% of its sites.

**(B)** SE of the introns contributing sites to each comparison, one value per intron; filled, the Figure 4 class; open, the same position outside DRACH. Introns with at least 20 reads at a branch point splice better than reference introns whatever the motif (medians 0.91 and 0.89 against 0.76); at exon +1 those reads must be unspliced, so the introns are retained (0.23).

**(C)** Each class against the intronic reference with 5-mer cells alone, then with SE (filled) and depth fixed; bootstrap 95% intervals. 5′SS A4 sites take the SE of their own intron. The branch point moves from 44 to 67% of expectation (32–111), exon +1 from at expectation to 60% (40–83), and 5′SS A4 reaches 133% (27–286). Binned by the same introns' DRS SE, HeLa $s^4$U-GLORI sites fall 7.9-fold in steady-state RNA, and matching on it moves the GLORI branch point from 56 to 65%. The Figure 4E gradient is 7.1-fold in all introns, 2.5-fold (1.5–3.8) where no other transcript's exon overlaps.

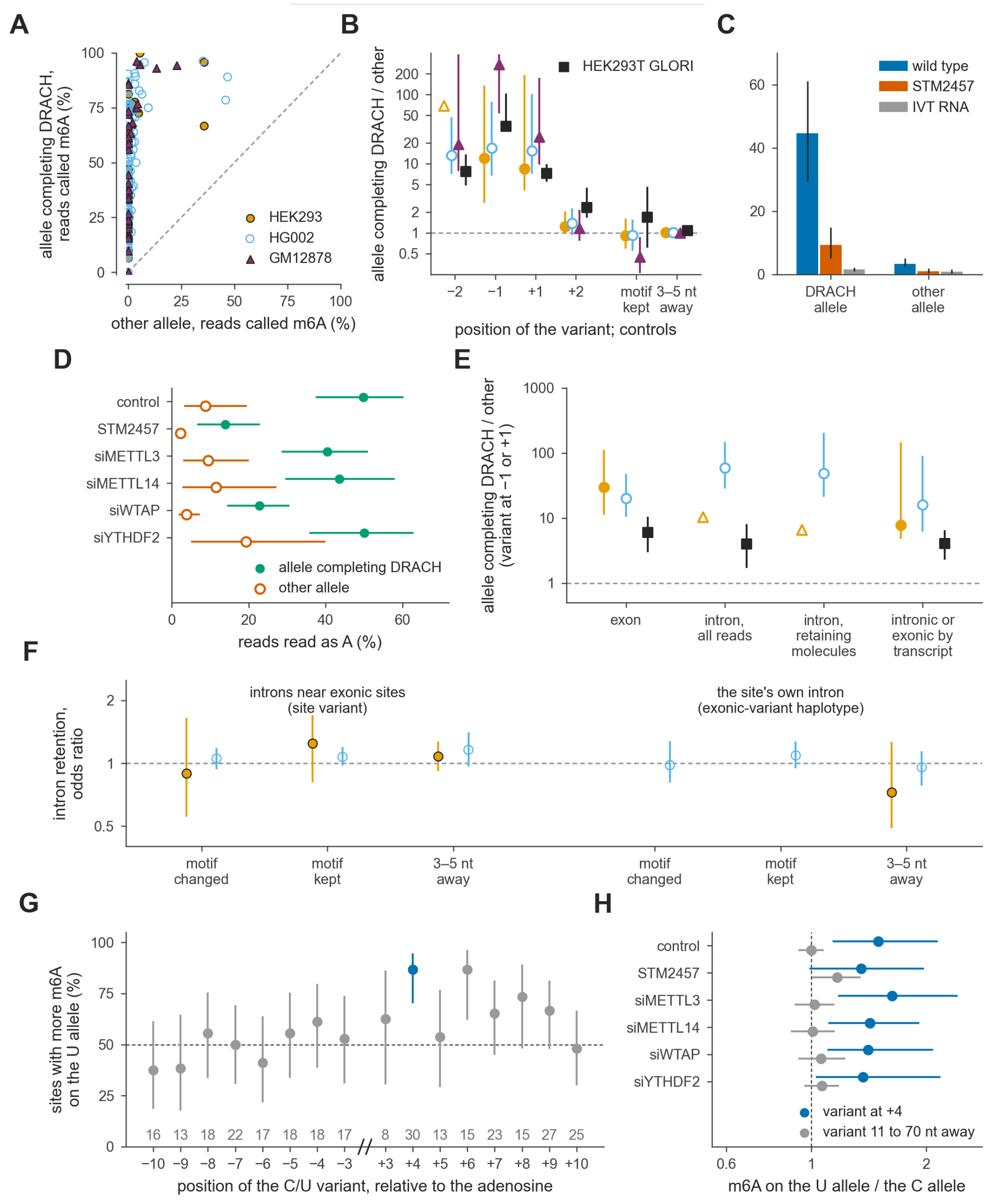

**Figure S11. A variant that completes the DRACH motif raises m⁶A on its own allele, related to Figure 5.**

**(A)** At heterozygous −1 or +1 variants, direct RNA reads called m⁶A on the allele completing DRACH against the other, at sites with five or more test reads per allele (HEK293 23 of 25, HG002 124 of 143, GM12878 42 of 43). Sites were selected on held-out runs (GM12878, halves of one flow cell) and scored on the others; pooled over all sites, 9.1-, 15.8- and 38.5-fold.

**(B)** The allele ratio by the position of the variant and for variants that keep the motif or lie 3 to 5 nt away, in the direct RNA sets and in HEK293T GLORI (54 sites at −1 or +1, 11.1-fold).

**(C)** GLORI, at the 64 sites whose DRACH allele is methylated: 44.6% in wild type, 9.4 with the METTL3 inhibitor STM2457 and 1.5 in unmodified RNA.

**(D)** HeLa GLORI, at the 138 sites whose DRACH allele is methylated in the controls: that allele reads 50% in the controls, 14 under STM2457 and 23 with WTAP depleted, while depleting the reader YTHDF2 leaves it at 50.

**(E)** The allele ratio by the location of the adenosine, for variants at −1 or +1, creating and breaking pooled, unselected; inside introns, on intron-retaining molecules alone, 48-fold (386 HG002 sites). Exon +1 has too few sites.

**(F)** Intron retention, DRACH allele over the other, at introns near exonic sites and, by exonic-variant haplotype, the site's own intron.

**(G)** In HEK293T GLORI, the share of C/U sites at which the U allele carries more m⁶A, by the variant's position, with sites at each; the peak is +4, 26 of the 30 that differ.

**(H)** The HeLa +4 U/C ratio in each arm against variants 11 to 70 nt away: 1.50 in the controls and 1.35 under STM2457, so the preference outlives the mark. Whiskers are 95% site-bootstrap intervals, Wilson in (G); open triangles in (B) and (E) are lower bounds where the other allele had no call.

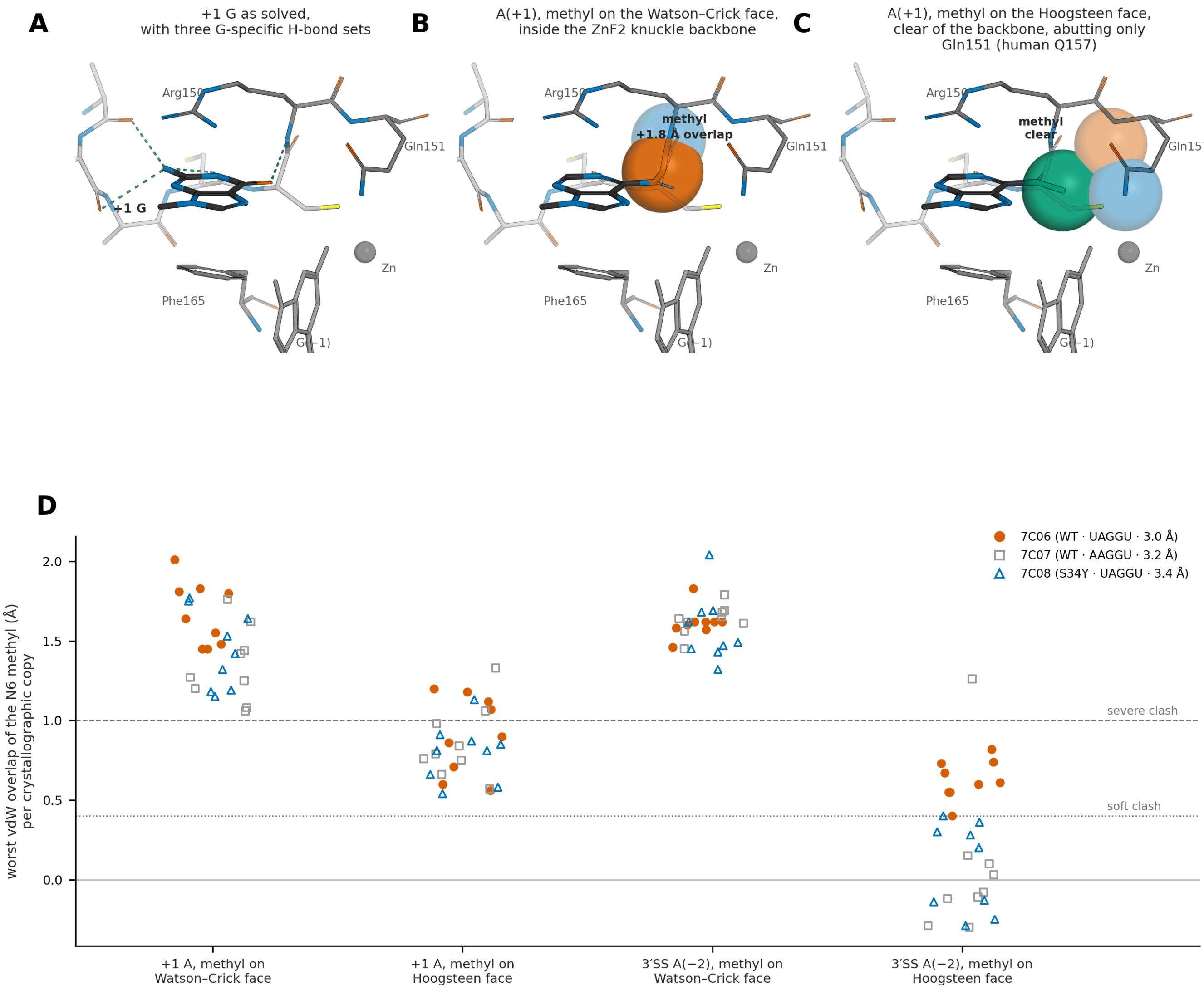

**Figure S12. U2AF1 reads the first exonic base, and m⁶A there clashes with its backbone in one orientation only, related to Figure 6.** Models in *S. pombe* U2AF1 (U2AF35) bound to a 3′ splice site (PDB 7C06, 7C07 and 7C08 [S12]; 27 copies), with an adenine at exon +1 and its $N^6$ methyl in each in-plane orientation.

**(A)** The deposited +1 G hydrogen-bonds the backbone through O6, N1 and N2 (dashes) and stacks on Phe165; an adenine loses these contacts, so the G-to-A change is the main loss of recognition.

**(B)** On the Watson–Crick face the methyl lies 1.45 Å from the Arg150 backbone nitrogen, a 1.8 Å overlap (spheres), and clashes with the backbone in 27 of 27 copies.

**(C)** On the Hoogsteen face it clears the backbone and meets only the Gln151 side chain (human Q157) and the G(−1) amino group.

**(D)** Worst methyl overlap per copy for both sites and orientations (dashed, 1.0 Å severe; dotted, 0.4 Å soft). At exon +1 the Hoogsteen-face contacts reach 0.54–1.33 Å and exceed 1.0 Å in 7 of 27 copies. Free $m^6A$ prefers the Watson–Crick orientation [S13]. The rigid-body models describe geometry, not binding affinity.

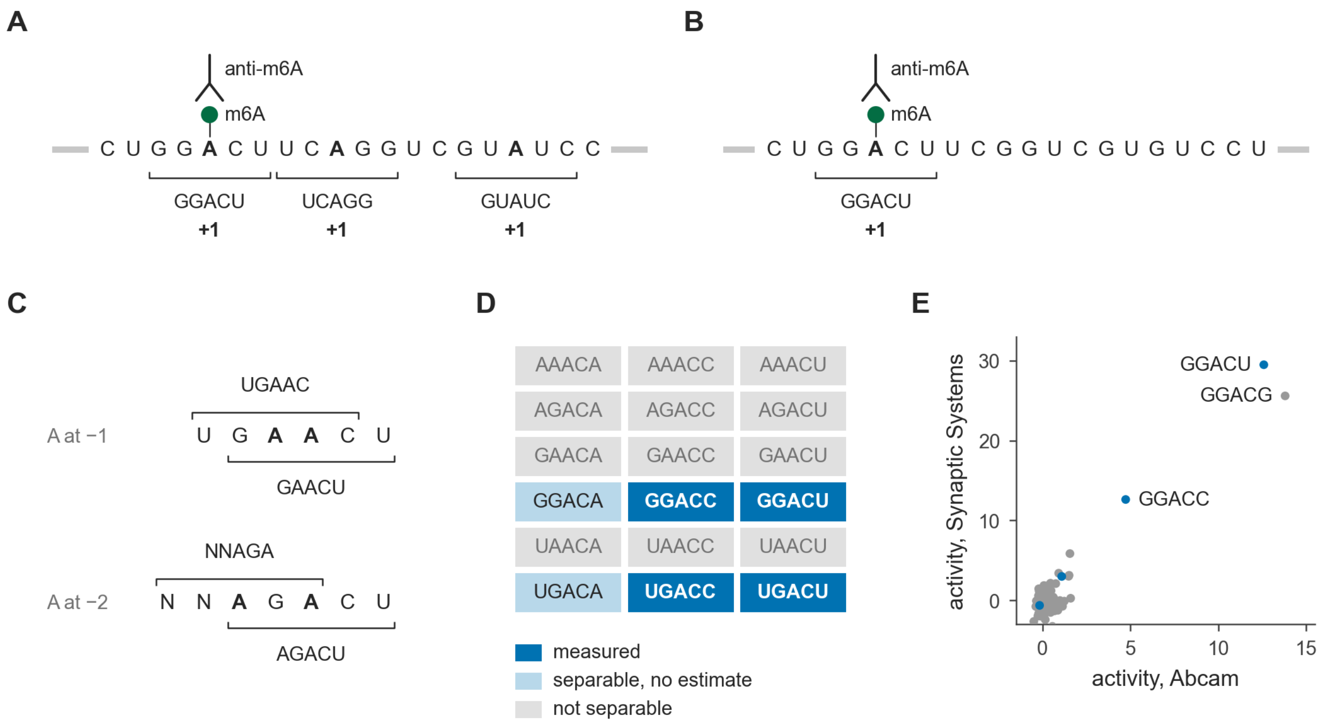

**Figure S13. Antibody selection cannot say which A of a randomized substrate was methylated, related to Figure 1.** In IVM-seq, METTL3–METTL14 methylates RNAs carrying a 20-nt random region between constant flanks (gray), and an anti-m$^6$A antibody selects methylated molecules [S1].

**(A)** Selection returns whole molecules, so every A-centered 5-mer on a selected molecule gains a count, although one A carries the mark (green).

**(B)** A molecule with a single A leaves one candidate. Such reads contain only the four DRACH 5-mers without a second A, and Figure 1D rests on them.

**(C)** A fit to all reads can apportion counts among 5-mers that occur in different combinations. An A at −1 or −2 instead places a second A-centered window on every molecule that carries the first, so no combination of reads separates the two.

**(D)** Of the 18 DRACH 5-mers, four are measured, two are separable in principle but have no valid estimate, and 12, every one with an A at −2 or −1, cannot be separated.

**(E)** Deconvolved activity of the 81 A-centered 5-mers the fit can estimate, in both antibody pools (Pearson r = 0.93). GGACG, GGACU and GGACC rank first to third in each.

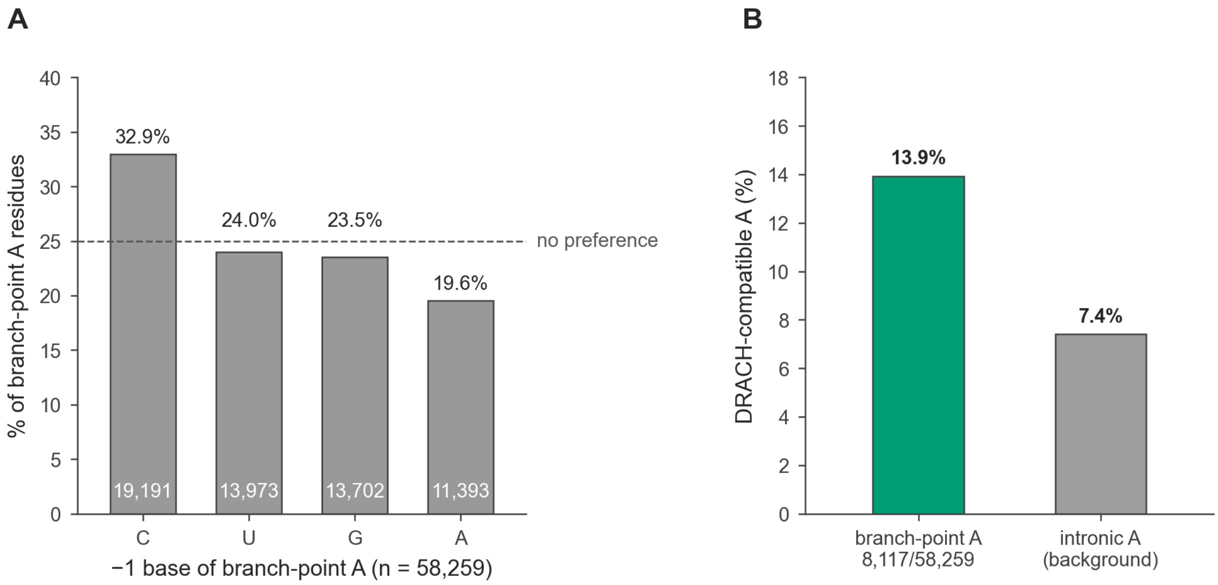

**Figure S14. Sequence context of human branch points, related to Figure 4.** Branch points mapped by lariat sequencing [S14,S15], of which 58,259 of 86,610 carry an A.

**(A)** The base before the branch A. Purines, which DRACH requires at this position, occupy 43.1%.

**(B)** DRACH-compatible contexts at 13.9% of branch-point A residues, against 7.4% of intronic A residues.

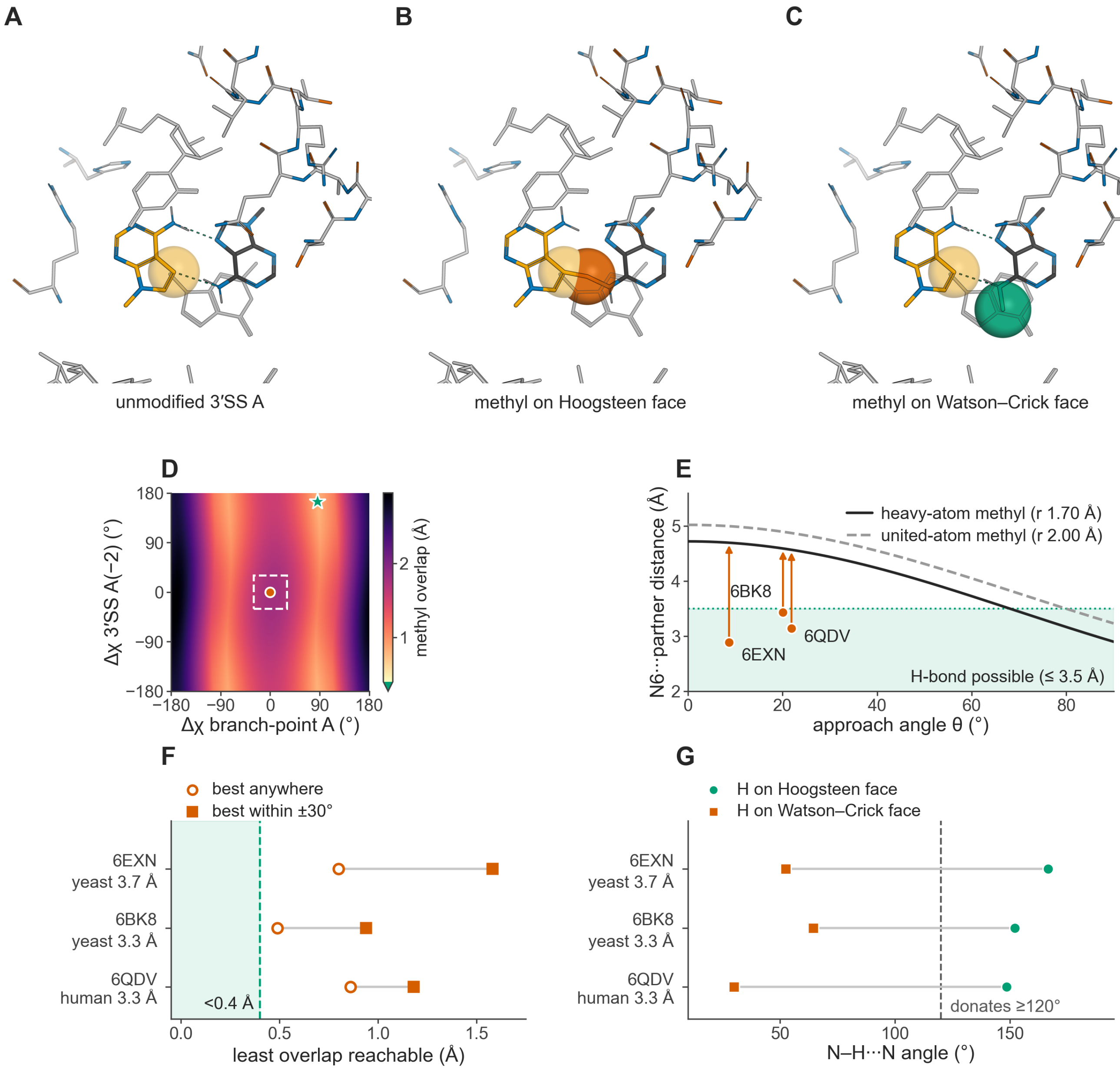

**Figure S15. m$^6$A at the docked 3′ splice site fits only in its preferred orientation, related to Figure 6.** Models in the post-catalytic P complex 6EXN [S16], with 6BK8 [S17] and the human 6QDV [S18] in (E)–(G).

**(A)** The unmodified 3′SS A(−2) and branch-point A each donate an N6–H to the other's N7 (2.89 and 2.88 Å).

**(B)** A Hoogsteen-face methyl replaces the donating hydrogen and overlaps the branch-point A by 1.80 Å.

**(C)** On the Watson–Crick face, which free m$^6$A prefers [S13], the methyl clears its neighbors and both hydrogen bonds stay intact.

**(D)** Rotating both glycosidic bonds (32,400 joint states) never brings the Hoogsteen-face methyl below the 0.4 Å threshold; the closest reaches 0.80 Å (star; circle, deposited pose; box, ±30°).

**(E)** The methyl lies 9–22° off the line joining the bases, so fitting it would push them 4.57–5.00 Å apart, beyond the 3.5 Å hydrogen-bond limit.

**(F)** No structure brings the overlap below the threshold; 6BK8 comes closest (0.49 Å).

**(G)** Only the Hoogsteen-face N–H points at the partner (148.5–166.6°), so methylation there removes the donor. The rigid-body models describe geometry, not reaction rates.

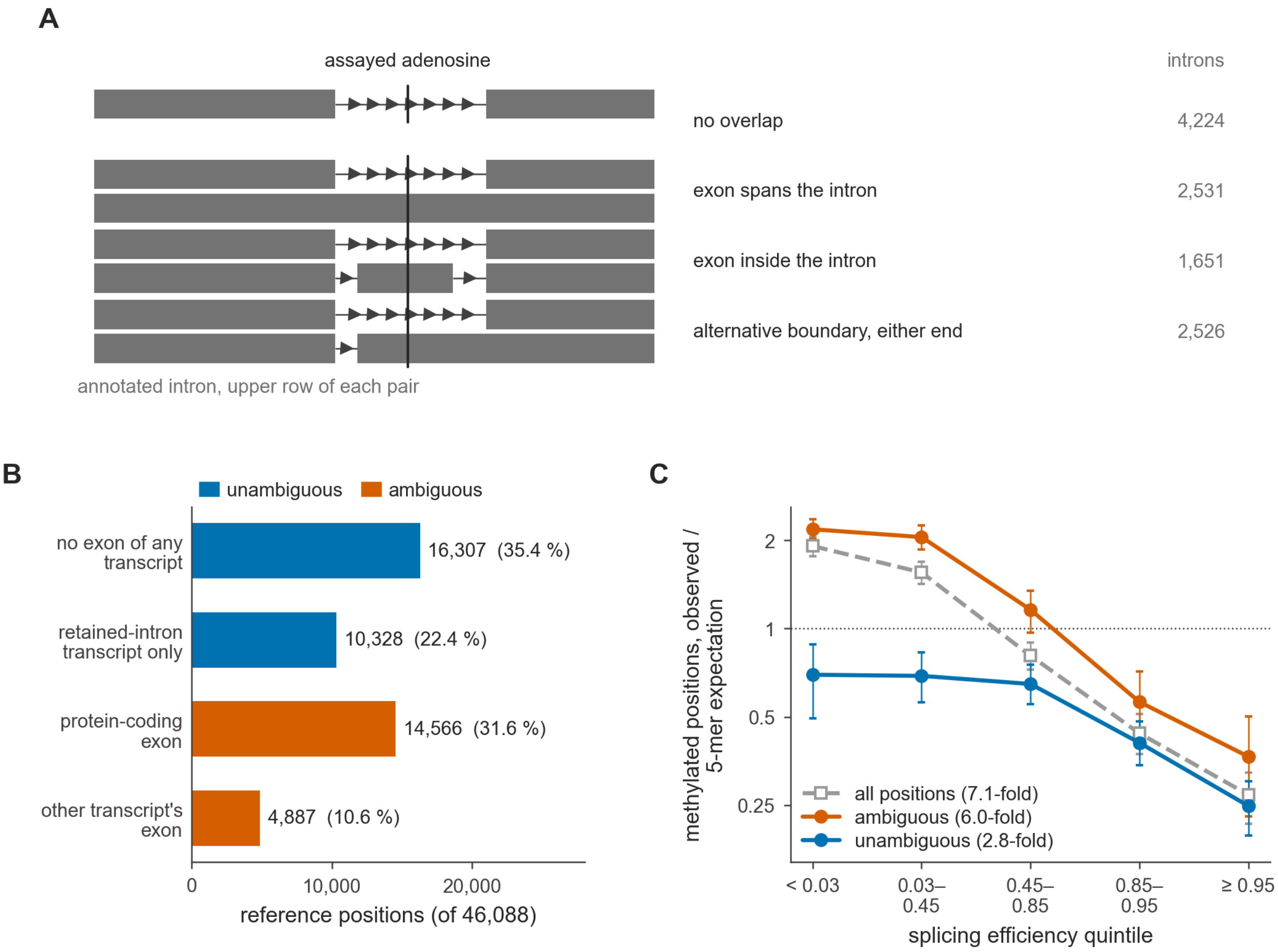

**Figure S16. Two fifths of the intronic reference positions lie inside an exon of another transcript, related to Figure 4.** HEK293 direct RNA. Reference positions are DRACH adenosines 50 to 500 nt upstream of a canonical 3′SS with at least 20 reads. A position is ambiguous when it lies inside an exon of another same-strand GENCODE v46 transcript that is not a retained-intron annotation.

**(A)** The ways such an exon can overlap an intron, with the number of the 10,932 reference introns in each. In each pair, the upper row is the annotated intron and the lower row is the other transcript; the two alternative-boundary forms share a row.

**(B)** The 46,088 reference positions by the transcripts in which they are exonic, with the 19,453 ambiguous positions (42%) in vermilion.

**(C)** Methylated share (positions called modified in at least 20% of reads) over its 5-mer expectation across the whole reference, by quintile of intron splicing efficiency, with 95% intervals from 1,000 bootstrap resamplings of introns. From the least to the best spliced quintile the share falls 7.1-fold over all positions, 6.0-fold over ambiguous and 2.8-fold over unambiguous ones.

### Supplemental notes

#### Supplemental Note 1. Historical evidence for methylation specificity and comparisons with splicing signals

The mRNA m$^6$A consensus was mapped biochemically long before the methyltransferase was identified. Nearest-neighbor analysis of mouse L-cell mRNA and hnRNA placed m$^6$A in the sequence N1-(G/A)-m$^6$A-C-N2, with G at −1 in about three-quarters of modified molecules, a purine at N1 about 90% of the time and G rarely at N2 [S19]. HeLa mRNA gave 70% Gm$^6$AC and 30% Am$^6$AC, with U, C or A following the C [S20], and avian sarcoma virus RNA yielded the same trinucleotides, with no specific sequence immediately upstream [S21]. Substrate assays and in vivo maps then refined these requirements. Harper et al. [S22] confirmed the consensus Pu[G>A]AC[A/C/U] with random copolymers and found sharply reduced methylation after single-base substitutions in defined GGACU-containing substrates; Csepany et al. [S23] found G, A or U, but never C, two positions upstream of the modified A across 12 Rous sarcoma virus sites; and Narayan et al. [S24] found methylation only within GAC or AAC, with 96% inhibition when the +1 C was replaced by U. By 1994 the preference for G over A at −1, the requirement for C at +1, the avoidance of G at +2 and the exclusion of C at −2 had all been established. Harper et al. [S22] also noted that such strict specificity was unusual for a factor acting on pre-mRNA, in contrast to splicing factors that tolerate departures from their consensus sequences, so a sequence that the spliceosome accepts may still be rejected by the methyltransferase.

Early comparisons with splicing signals centered on the branch site. Zeitlin and Efstratiadis [S25] proposed that m$^6$A might mark A residues for lariat formation because its context resembled the branch-point consensus, while noting that branch sequences lacked the invariant form then reported for methylation sites; their rabbit β-globin branch A, the penultimate A of TGCTAAC, lies in UAAC, a DRACH-compatible core. Kane and Beemon [S26] found the PuGACU methylation context similar but not identical to the branch-site consensus and proposed either that branch points could be methylated or that methylation could block the use of sequences resembling them, although they took exonic m$^6$A as evidence against a positive role in branch-site selection. Narayan et al. [S27] concluded from influenza virus experiments that internal A methylation did not promote splicing in that system. Carroll et al. [S28] mapped the predominant m$^6$A in the final intron of a bovine prolactin minigene to AGm$^6$ACU, in about 24% of molecules, and noted that methylation in a segment removed by splicing need not persist in the mature transcript. A substrate bearing a bulky N6 substituent still formed lariats, but the branching nucleotide could have been the modified A or an adjacent upstream guanosine, so that experiment neither established branching through a modified A nor tested m$^6$A [S29]. Shimba et al. [S30] distinguished the U6 snRNA A43 context, UACm$^6$AGA, from the mRNA consensus; it places C at −1 and G at +1, the METTL16-family register discussed in the main text.

Later interpretations depended on the consensus used. Martinez and Gilbert [S31] treated the 5′ splice site GURAGU, the branch site YNYURAC and the 3′ splice site YAG as unlikely methylation targets under a strict RRACH requirement. The −2 U of the branch-site consensus is allowed by DRACH but excluded by RRACH, whereas the canonical 3′SS A fails both because of its +1 G. Watabe et al. [S32] noted that no m$^6$A had been reported at the invariant AG of mammalian 3′ splice sites, alongside their demonstration of METT-10-dependent 3′SS methylation in *C. elegans*.

#### Supplemental Note 2. Isoform ambiguity in the intronic reference

Intronic positions that lie inside another transcript's exon carry most of the methylation near 3′ splice sites. Three tests indicate that the reads at these positions come from precursors rather than from the other transcript's mRNA, and the central result does not depend on them. Ambiguous positions, overlapped introns and the three tests are defined in STAR Methods, and the intronic reference is made of DRACH adenosines that lie 50 to 500 nucleotides upstream of canonical 3′ splice sites and carry at least 20 direct RNA reads in HEK293 cells.

Of the 46,088 reference positions, 19,453 (42%) are ambiguous, and three quarters of these (14,566) lie inside exons of protein-coding transcripts (Figure S16B). Of the 10,932 introns that carry reference positions, 6,708 (61%) are overlapped, and a cassette exon inside the intron accounts for only a quarter of them (1,651). In 2,531 introns an exon of another transcript spans the whole intron, and in 2,526 it crosses one end of the intron (Figure S16A). At the median, the overlapping exon covers 50% of its intron, and the median reference intron is 814 nucleotides long.

The overlap is concentrated in poorly spliced introns, where ambiguous positions make up 81% of the least spliced quintile of the reference against 18% of the best spliced one. Pooled over all reference positions, the methylated share falls 7.1-fold from the least to the best spliced quintile (Figure 4E). The fall is 6.0-fold over ambiguous positions and 2.8-fold over unambiguous ones (Figure S16C), and in every group the 95% intervals of the two extreme quintiles do not overlap, so the gradient itself is not an artifact of ambiguity. Classified by intron rather than by position, the gradient is 7.1-fold over all introns and 2.5-fold (95% CI 1.5 to 3.8) over introns that no other transcript's exon overlaps (Figure S10C).

Three observations suggest that the reads at ambiguous positions come largely from precursors of the host transcript rather than from mature RNA of the other transcript. First, mature RNA carries exon–exon junctions, yet reads counted unspliced at introns that another transcript's exon spans carry 0.92 junctions per kilobase, as unspliced reads at introns without overlap do (0.94), against 2.22 for reads spliced at the same introns. Second, if the methylation at ambiguous positions came from mature RNA of the other transcript, the reads called modified there would carry more junctions than the unmodified reads at the same positions. In every subset tested, by the kind of exon, the distance from the other transcript's stop codon, the geometry of the overlap and splicing efficiency, the junction difference between modified and unmodified reads is smaller than at positions that no other transcript covers. Both tests lose power where the overlapping exon is the other transcript's last exon, because a direct RNA read begins at the poly(A) tail and can end before it reaches a junction. Third, the higher methylation of ambiguous positions is already present in newly made RNA. In HeLa $s^4$U-GLORI libraries [S10], with splicing efficiency taken from the direct RNA reads of the same introns, ambiguous positions of the two least spliced quintiles are 5.10 times as methylated as unambiguous ones in nascent RNA without a poly(A) tail, labeled for 5 minutes (95% CI 3.60 to 7.53), 4.35 times in nascent poly(A)+ RNA and 3.55 times in steady-state poly(A)+ RNA.

Removing ambiguous positions from the reference would clean the reference but not the splicing signals compared with it, and many of those signals sit in overlapped introns (71% of exon +1 sites, 72% of 5′SS A4 sites and 49% of branch points). We therefore kept every position and added intron overlap to the matching. Matched on 5-mer, splicing efficiency and intron overlap, branch points reach 68% of the expected level (95% CI 34 to 111%), exon +1 reaches 58% (39 to 81%) and 5′SS A4 reaches 121% (26 to 268%), close to the values without this matching (67%, 60% and 133%; Figure S10C).

Ambiguity also shapes the profiles in Figure 2. In direct RNA, the share of intronic DRACH positions that lie inside another transcript's exon rises toward the 3′ splice site, from 42 to 45% at positions more than 50 nucleotides upstream to 53% at 21 to 50 nucleotides and 61% in the last 20, and at every distance these positions carry most of the methylated sites. Positions that no other transcript covers are methylated at one level all the way to the 3′ splice site (34% of the level expected from their 5-mers in the last 20 nucleotides against 32% at 201 to 500 nucleotides), so methylation still reaches the last intronic nucleotides (dashed line in Figure 2D). Pooled over all positions, the last 50 intronic nucleotides appear more methylated than exonic positions 50 to 100 nucleotides downstream (80% and 81% against 61% of the expected level), whereas positions that no other transcript covers reach 34%, little more than half the exonic level. The 3′ splice-site adenosine is unaffected, because reads misassigned from another transcript could only add methylation, and that adenosine stays at background.

At the same splicing efficiency, positions in overlapped introns remain more methylated than positions in introns without overlap. Below a splicing efficiency of 0.05 they are 3.4 times as methylated, and the ratio falls to 1.1 above 0.97. In $s^4$U-GLORI, the excess is larger at ambiguous positions within 500 nucleotides of the other transcript's stop codon, where $m^6A$ concentrates in mRNA [S33]. The observations above argue against contamination by mature RNA of the other transcript, and the cause of the excess is not known. It does not bear on the central result, because the exclusion of the canonical 3′ splice-site adenosine rests on the writer's sequence specificity and on direct measurements at that adenosine (Figures 1–3).

#### Supplemental Note 3. 3′ splice-site choice at STX10

*STX10* carries a methylatable AT–AC 3′SS beside an alternative 3′SS and first suggested that methylation might shift the choice between them. Across 58 public perturbation libraries, however, writer and adaptor knockdowns reduced the use of its methylatable 3′SS rather than increasing it, so an effect of the mark on splicing efficiency or 3′SS choice is not established. This is a descriptive pooled comparison at one of the 11 pairs observable in both arms, from 15 screened, without adjustment for locus selection (STAR Methods).

#### Supplemental Note 4. Exon +1 methylation at 3′ splice sites that another transcript's exon covers

The first exonic nucleotide of a canonical 3′ splice site can also lie inside an exon of another annotated transcript, and such covered 3′ splice sites, together with the read classes used below, are defined in STAR Methods. Covered 3′ splice sites are frequent where exon +1 is methylated. Of the 149 DRACH-compatible exon +1 acceptors with a GLORI signal in HeLa or HEK293T polyadenylated RNA, 86 (58%) are covered, against 1,193 of the 8,858 acceptors without one (13.5%; Figure S7B).

Exon +1 is also methylated more often at covered 3′ splice sites. In HEK293 direct RNA reads that cross the 3′ splice site unspliced, the modified-read fractions are 3.37% at covered and 1.87% at uncovered 3′ splice sites. These reads are long enough to test whether the covering transcript's mRNA carries the mark. At the same site, reads of the covering mRNA carry exon +1 methylation at 0.86 times the level of precursor reads (95% CI 0.67 to 1.10, 177 sites; Figure S7C), so the higher rate at covered 3′ splice sites does not come from mature RNA of the covering transcript. The excess is a property of the sites. Averaged over sites, precursor reads are modified in 6.0% of cases at covered 3′ splice sites and in 1.8% at uncovered ones.

The retention association depends on coverage. At a covered 3′ splice site, a short read that crosses the boundary without a gap can come from the precursor or from the covering transcript's mRNA, and 55 of the 77 3′ splice sites with a methylated exon +1 in the retention comparison of Figure S8E are covered. Their retention exceeds that of matched controls by 0.33 at covered 3′ splice sites and by 0.011 (95% CI 0.000 to 0.078) at the 22 uncovered ones (Figure S7D). An association between exon +1 methylation and intron retention is therefore not detectable where no other transcript covers the 3′ splice site. The polypyrimidine-tract difference of Figures S8A and S8B keeps its direction at uncovered 3′ splice sites, with a median difference of −2 uridines over positions −40 to −3 (95% CI −3.0 to −0.8, n = 58), and it carries less power than the full comparison.
